# Leveraging species-wide variation and patterns of adaptation to inform pecan crop improvement efforts

**DOI:** 10.64898/2025.12.19.695279

**Authors:** Paul P Grabowski, Chloee M McLaughlin, Joanna Rifkin, Jerry J Jenkins, Angelyn Hilton, Hormat Shadgou Rhein, Kimberly Cervantes, Motoyuki Ishimori, Avinash Sreedasyam, Chris Plott, Jenell Webber, Nolan Bentley, Gaurab Bhattarai, Paul Oladimeji Gabriel, Anna Harmon, Keith Kubenka, Clive Bock, Warren Chatwin, Patrick Conner, LJ Grauke, Jane Grimwood, Hiroyoshi Iwata, Patricia Klein, Christopher P Mattison, Cristina Pisani, Jeremy Schmutz, Joshua Udall, Xinwang Wang, Jennifer Randall, John T. Lovell

## Abstract

The genetic basis of adaptation is a fundamental question in evolutionary biology, and understanding how species will be able to adapt to changing conditions across their range has important implications for conservation and agriculture. To accurately interrogate the genetics of adaptation and assess the adaptive capacity of a species requires also characterizing the ways other mechanisms, including geographic distance and population dynamics, shape genetic variation. Pecan is an ecologically, culturally, and economically important North American tree, and a broader understanding of the genetics of environment adaptation will aid pecan conservation, breeding, and commercial management. Here, we use an expansive set of more than 700 pecan genotypes in combination with the first haplotype-resolved genome assembly for pecan to assess species-wide genetic variation and evaluate environmental adaptation across the native distribution. We identify five gene pools in pecan, with the lowest diversity in southern gene pools, and present evidence that gene pools began differentiating during multiple glacial cycles. Using complementary genotype-environment association approaches, we infer species-wide patterns of environmental adaptation. With these results, we predict mismatches in adaptation for pecan genotypes to different environments, including future environment scenarios. We see that in all locations, present-day genotypes incur some level of predicted maladaptation to simulated future environments, but current genetic diversity may provide a valuable source of resilience to future conditions through assisted migration. These results expand the understanding of environmental adaptation in pecan and provide insight into how long-lived species will be able to adapt to future conditions.

## Introduction

Understanding how adaptation shapes genetic variation is an important question in evolutionary biology (Savolainen, Lascoux, and Merilä 2013) with applications for crop breeding (Xu et al. 2006; Morris et al. 2025) and agricultural production (Rosenzweig et al. 2014; Varshney et al. 2018; Xiong, Reynolds, and Xu 2022). For long-lived plant species, fitness depends on enduring local conditions for decades. However, in the case of rapidly changing environments, local conditions may change drastically over less than a lifetime for an individual, and the alleles conferring adaptation to current conditions may not be suitable for future conditions (Aitken and Whitlock 2013). Characterizing the genetic basis of environmental adaptation can be used to determine the adaptive capacity of a species (Vanhove et al. 2021) and can help identify populations and regions that are particularly well suited or vulnerable to predicted changes in the environment (Aitken et al. 2008).

Adaptation to local conditions generates genetic differences within a species that reflect the encountered environmental variation (Leimu and Fischer 2008). However, at the genomic level, those adaptive genetic differences can be obfuscated by the other mechanisms that also shape variation in a species. For instance, gene flow decreases with geographic distance, so neutral genetic differences that accumulate between distant populations are autocorrelated with the genetic changes generated by adaptation to different environments (I. J. Wang and Bradburd 2014). Similarly, species frequently have genetic structure across their range resulting from the location of suitable refugium habitat during ice ages (Roberts and Hamann 2015), and adaptive differences between differentiated populations co-occur with the neutral difference generated during periods of allopatry (Lovell, MacQueen, et al. 2021). Characterizing the roles of different mechanisms in shaping variation helps inform expectations as to how a species is adapted to its environment and facilitates a more accurate interrogation of the genetics of adaptation and the adaptive capacity of the species.

The genetic basis of local adaptation can be effectively characterized through genotype-environment associations (GEAs), statistical correlations between genetic variants and environmental variables (Lasky, Josephs, and Morris 2023). GEAs have been widely used to investigate loci that may play key roles in environmental adaptation (Rellstab et al. 2015) and to understand how genetic variation correlates with specific environmental axes (Capblancq and Forester 2021). A rising approach is to use fitted GEA models, which relate differences in allele frequencies to variation in environments genotypes are sourced from (and assumed to be adapted to), and projecting what degree of genomic change is needed for maintained adaptation to novel environments. The resulting genomic change that is required for persisting adaptation is a measure of maladaptation, often referred to as genomic offset (reviewed in (Rellstab, Dauphin, and Exposito-Alonso 2021). Beyond forecasting maladaptation risk, GEAs can also be extended to identify optimal genotypes or varieties for particular target environments (Rhoné et al. 2020; Caproni et al. 2023; M McLaughlin et al. 2025), providing a guide for assisted migration or targeted breeding efforts.

Pecan (*Carya illinoinensis* (Wangehn.) K. Koch) is an ecologically, culturally, and economically important long-lived North American tree. Its broad distribution across environmental gradients, spanning from Central Mexico to the Midwest USA (Grauke, Wood, and Harris 2016), offers a powerful system for exploring genotype-environment associations and the genetics of adaptation. Pecan produces nutritious nuts with annual commercial production in the USA valued at more than $500M (United States Department of Agriculture, National Agricultural Statistics Service 2025a, 2025b). The history of pecan cultivation dates back thousands of years, and populations were managed and relocated by indigenous cultivators from prehistory onwards (Fritz 2016; Hal 2008; Grauke et al. 2011). Pecan yields are impacted by drought (Garrot et al. 1993), temperature stress (Sparks 2000; L. Wang et al. 2025), insect pests (Mulder, Harris, and Grantham 2012; Grauke, Wood, and Harris 2016), and disease (Gottwald and Bertrand 1988), all of which vary across its range. There is substantial phenotypic variation in how pecan responds to environmental stresses (Bock et al. 2020), and understanding the genetic basis of how pecan has adapted to environmental stress can help guide breeding and commercial decisions. The pecan research community has developed extensive genetic resources, including germplasm collections with native genotypes from across the range and contemporary and historical breeding varieties (Grauke, Wood, and Harris 2016). In addition, powerful genomic resources have been developed to facilitate studies of traits and adaptation at the genome scale (Lovell, Bentley, et al. 2021; Huang et al. 2019). High quality genome assemblies helped to reveal the complex genome structure underlying an important QTL for resistance to a destructive pest (Lovell, Bentley, et al. 2021), and applying genomic resources to the extensive germplasm collections is a powerful approach for more completely characterizing the extent of standing and potentially adaptive variation in pecans, the genetic basis of environmental adaptation, and the extent of ecological mismatch in current and future conditions.

Here, we present an improved genome for the ‘Oaxaca’ (87MX3-2.11) pecan cultivar, producing the first haplotype-resolved genome assembly in pecan. Using the updated assembly, we measure genetic variation across the range of pecan using genotypes from more than 700 samples. The results reveal that variation is distributed across five regional gene pools that diverged during the last two North American glacial periods and that contemporary patterns of genetic differences reflect processes that began more than 100k years ago. Using genetic and environmental data for more than 500 georeferenced native pecan samples, we implement complementary GEA methods to characterize patterns of environmental adaptation across the range, identify putatively adaptive loci, and elucidate the genetic architecture of environmental adaptation. We use these results to predict how pecan will respond to potential changes in the environment and which genotypes will be better adapted to current and future environment conditions. These results reveal that at all locations, future environments are expected to disrupt current adaptation, but that variation exists to minimize maladaptation to future conditions across the range, though populations in Central Mexico and parts of Texas may remain vulnerable to future environment conditions. Combined, our results can help inform priorities for pecan conservation and breeding and expand on the understanding of environmental adaptation in long-lived, wide-ranging species.

## Results and Discussion

### Resources to enable genome-informed breeding in pecan

Pecan trees are physically large and take many years to mature, which present challenges to the speed and scale of crop improvement efforts. Genome-enabled trait discovery can greatly improve the efficiency of breeding (Collard and Mackill 2008), however, genetic loci underlying traits often lie in complex genomic regions and therefore require a high-quality reference genome for accurate and precise characterization (Healey et al. 2024; Mascher et al. 2021). Despite its demonstrated use in genetic mapping (Brungardt et al. 2024; Rhein et al. 2023), the existing ‘V1’ pecan reference genome for the 87MX3-2.11 (hereon, ‘Oaxaca’, (Lovell, Bentley, et al. 2021)) pecan genotype was built with PacBio consensus long reads (CLR) technology, which requires a single-sequence build for outbred genotypes and can underrepresent repetitive sequences like centromeres.

To support future genome-informed breeding in pecan, we generated an improved and haplotype-resolved ‘V2’ ‘Oaxaca’ assembly. In short (see methods for details), we coassembled PacBio HiFi (38.56X coverage) and Hi-C (40.0X coverage) reads using HiFiAsm+HIC (Cheng et al. 2021) and scaffolded the contigs using the JUICER (Durand et al. 2016) pipeline. The resulting haplotype assemblies of 670.0Mb (HAP1) and 659.0Mb (HAP2) were polished with 55.5X Illumina 2x150 reads to correct homozygous SNPs and short indels. A total of 100 and 86 contigs were used to assemble the 16 chromosomes of each haplotype, ranging from 1 to 13 contigs per chromosome (Fig. S1), and 100% (HAP1) and 99.98% (HAP2) of the sequence is contained on the chromosome scaffolds of the assemblies, producing contig N50 values of 14.0Mb and 15.9Mb (Table S1). To maintain consistency with ongoing efforts that use our V1 genomes, we opted to ‘project’ gene models from (Lovell, Bentley, et al. 2021) onto V2, which generated 34,344 gene models. We also characterized repetitive content using EDTA (Ou et al. 2019), with 46.5% and 46.2% of the assemblies containing annotated repetitive sequence (Fig. S2, Table S2).

The Oaxaca.v2 assemblies are larger and more complete than the V1 assembly, with the assemblies containing 20.0Mb (HAP1) and 9.0Mb (HAP2) more total sequence than V1 and 5.20% (33.06Mb) and 3.44% (21.94Mb) more sequence assembled into chromosomes. The more than 5X improvement in contiguity (i.e.: number of contigs used in each assembly) and 3X improvement in contig N50 in the Oaxaca.v2 assemblies compared to Oaxaca.v1 (Fig. S1, Table S1; (Lovell, Bentley, et al. 2021)) provides a more thorough representation of the complex genome regions that complicate assembling contigs into chromosome-length sequences. For instance, the putative pericentromeric regions of V2, defined as extended regions with elevated repetitive content and <1.0% genic sequence, contain 18.17Mb (HAP1) and 14.53Mb (HAP2) more sequence than in V1, including more than 2Mb additional sequence in the putative pericentromeres of both haplotypes of Chr06 and Chr12. Furthermore, V2 is the first haplotype-resolved genome for pecan, which will facilitate more accurate analyses of structural variation and of allele-specific traits, including allele-specific expression and presence-absence variation. For example, V2 reveals a >1.7Mb inversion between HAP1 and HAP2 in sequence of Chr12 that could not be incorporated into chromosomes in V1.

To aid in characterizing genetic variation and the genetic basis of important traits in pecan, we conducted whole-genome resequencing (WGS) of 683 pecan samples using Illumina paired-end reads to a mean sequencing depth of 39.42X (Table S3), generating genotypes at more than 56.2 million single nucleotide polymorphisms (SNPs) and 4.3 million short (< 60bp) insertion-deletion polymorphisms (INDELs), across the genome. For our population genetic analysis, we downsampled to 46.4 million bi-allelic SNPs to best describe patterns of diversity across the species. The samples include 466 ‘native’ samples (trees grown from wild-origin seed collected from georeferenced native populations) and 217 genotypes related to pecan breeding in the USA (Table S3). These genotypes provide dense genotyping information for most cultivars commonly used for research and breeding and provide a powerful resource for interrogating the genomic basis of traits across diverse backgrounds. In particular, the 466 native samples represent vital diversity for adaptation across the native range, exploitation of which may prove valuable for supporting future breeding efforts.

While we sought to maximize the geographic diversity by sampling a single WGS genotype at any georeferenced position, we were also interested in within-population allele frequencies. Therefore, we also sequenced 779 additional samples, including 581 ‘native’ samples, using genotype-by-sequencing (GBS) (Table S4), obtaining genotypes at 17,920 SNPs. These ‘native’ samples represent 206 populations with unique geocoordinates, and 77 populations are represented with multiple samples (452 genotypes from these 77 populations). The GBS within-population sampling gives us the ability to more accurately estimate heterozygosity within populations.

### Genetic variation in pecan reveals differences in diversity across the range of pecan

Pecan has a large geographic range across diverse environments, with the native range spanning from Central Mexico into the Midwest USA (Wood, Grauke, and Payne 1998). As in other broadly distributed species (Murray et al. 2019; Lowry et al. 2014; J. Wang et al. 2016), pecan exhibits significant phenotypic and molecular variation across its range (Bentley, Grauke, and Klein 2019); (X. Wang et al. 2024). Despite extensive within-species variation, previous studies of pecan diversity have been limited by sampling density and resolution of genetic markers. The resources we have generated here, including a diversity panel of more than 650 pecan samples with high-density SNP markers at more than 150 million sites, form a strong foundation to more fully explore the patterns and processes that have shaped pecan genetic diversity. For example, our population structure analysis using 466 native pecan genotypes profits from vastly improved sampling of diversity in allele frequencies to identify five genetically distinct gene pools (Fig. 1A; Fig. 1B; Fig. S3; Fig. S4) varying in geographic distributions (Fig 1A) and underlying genetic structure (range of Fst between gene pools = 0.032 - 0.119; Table S5). Additionally, this more complete characterization of species-wide diversity offers a valuable opportunity to relate genetic variation to differences in the range of environments inhabited by pecan.

**Figure 1.**
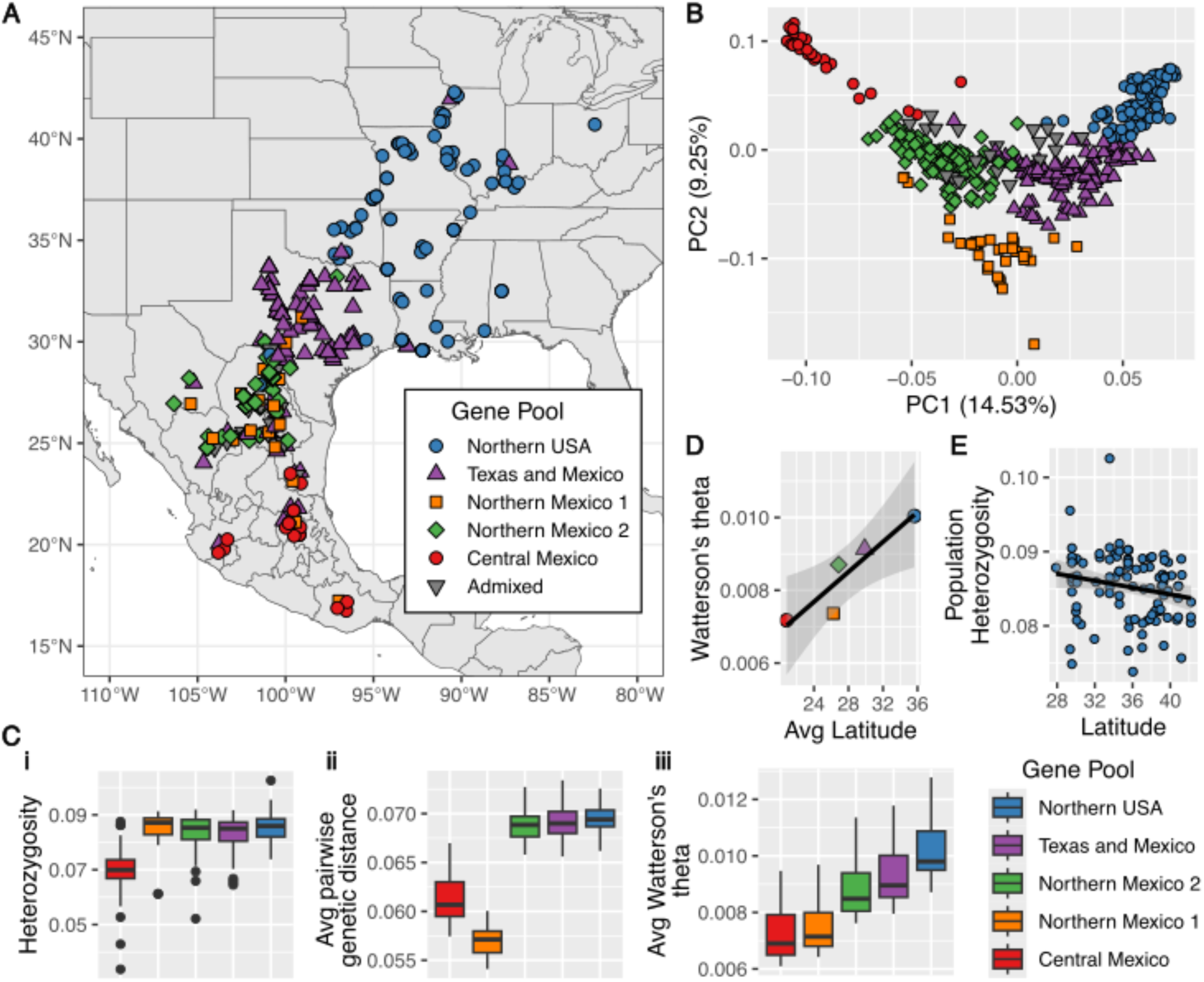
Pecan genetic variation is structured into regional gene pools with differences in genetic diversity. **A**. Geographic distribution of 466 native pecan samples colored by membership in the 5 native pecan gene pools. Genotypes without >50% ancestry from any gene pool are labeled as Admixed. **B.** Principal component analysis (PCA) using the 466 native pecan samples colored by gene pool as in (A). **C.** Genetic diversity differences between pecan gene pools. **i.** Heterozygosity **ii.** Average pairwise genetic distance. **iii.** Average chromosome-wide Watterson’s theta. **D.** Latitudinal gradient of gene pool Watterson’s theta. Points colored by gene pool as in (A). **E**. Latitudinal gradient of heterozygosity in populations of the Northern USA gene pool.

Genetic variation of many temperate plant species has been strongly shaped by glacial cycles (Hewitt 2000). In particular, the number and size of discrete refugia during glacial maxima (and effective population size therein) can have lasting impacts on the population structure and diversity of extant populations. Demographic models using the native pecan genotypes indicate the initial divergence of gene pools 9,800 to 15,700 generations ago, corresponding to an estimated time range of 147 kya - 235 kya (15 year generation time) or 196 kya - 314 kya (20 year generation time). This most ancient divergence formed the precursors to the Central Mexico gene pool, North Mexico 1 gene pool, and a third population (Table S6). The third population subsequently had an estimated divergence of 4,690 to 6,700 generations ago (70 kya - 100 kya, 15 year generation time; 94 kya - 134 kya, 20 year generation time), to form the North Mexico 2, Texas-Mexico, and Northern USA gene pools (Table S6). Importantly, our estimates from demographic models suggest overlap between the initial population divergence with the Illinoian glacial stage (132kya - 302kya) and the divergence of the Northern populations with the Wisconsin glaciation (14 kya - 109 kya, (Richmond and Fullerton 1986)). Together, these results suggest that glacial cycles played a critical role in driving range shifts and geographic isolation, ultimately shaping the underlying divergence of native pecan gene pools.

Dynamics of post-glaciation expansion and colonization can also have lasting impacts on contemporary genetic diversity. As populations emerged from glacial refugia, populations typically recolonized habitats similar to their refugial habitat (Hewitt 2000; Waltari et al. 2007) and the extent of their expansion depended on the amount and proximity of suitable habitat. The Northern USA gene pool, which has the most extensive northward expansion, was genetically distinct prior to its expansion into its current range, which suggests that the Northern USA gene pool may have been better adapted than other gene pools to colonize habitats at the northern edge of the range. In contrast, in the southern edge of the range, the post-glacial expansion of the Central Mexico and Northern Mexico 1 gene pools was more restricted, and they are only found in disjointed patches of habitat (Grauke, Wood, and Harris 2016). Diversity measures reflect these contrasting population dynamics at the different ends of the range. The Central Mexico and North Mexico 1 gene pools have lower nucleotide diversity and Watterson theta values (*p*-value < 1e-24 in two-sample Kolmogorov-Smirnov tests; Fig. 1C) than the three other gene pools, and the Central Mexico gene pool has lower heterozygosity than the other gene pools (*p*-value < 3x10-9 in two-sample Kolmogorov-Smirnov tests; Fig. 1C). Furthermore, there is a positive relationship between mean latitude of gene pools and Watterson theta values (*p*-value < 0.026; Fig. 1D). Temperate species often show a latitudinal diversity gradient, with genetic diversity decreasing at higher latitudes (Hewitt 2000), but species-wide, pecan shows a different pattern, as the more southern populations have lower diversity and lower effective population sizes, perhaps due to more limited habitat availability in their geographic regions compared to the northern part of the range. The effective migration surface also suggests geographic barriers, such as the Sierra Madre Occidental, limit gene flow in the southern portion of the range (Fig. S5). While the Northern USA gene pool overall has comparatively high diversity, within the Northern USA gene pool, there is a trend of lower heterozygosity at higher latitudes (*p*-value < 0.043 for 101 WGS localities; *p*-value < 0.047 for 85 GBS localities; Fig. 1E; Fig. S6), consistent with the expectation of higher diversity near the glacial refuge and lower diversity at the edge of an expanding population due to founder effects (Hewitt 2000; Eckert, Samis, and Lougheed 2008).

### Using genetic associations with environment to predict adaptive differences in native pecan

The variety of environment conditions across the range indicate ongoing adaptive differentiation in pecan; however, the lasting legacy of glacial refugia and post-glacial colonization on patterns of variation described above complicates efforts to interrogate the genetic basis of adaptation. For example, pecan gene pools have different geographic distributions (Fig 1A), and historical differentiation can be correlated with contemporary environmental differences that shape adaptive variation. Even within gene pools, environmental gradients can be correlated with geographic separation and genetic signatures of isolation-by-distance. As such, evaluating the genetic basis of adaptation in pecan requires disentangling the effects of historical separation and geographic distance from those of environmental differences shaping adaptation across the species.

Therefore, to characterize how environment adaptation shapes diversity in pecan, we first used partial redundancy analysis (pRDA) (Capblancq and Forester 2021) to identify loci that capture genome-wide patterns of genetic variation associated with environmental gradients, which may reflect signals of local adaptation. For this step and downstream pRDA construction, we used a smaller set of 300,000 variants, as forward selection is computationally intensive and becomes increasingly demanding with larger numbers of variants. We used forward selection (Oksanen et al. 2022) to determine which aspects of the bioclimate to include in our pRDA models, retaining 12 of the 19 standard bioclimatic variables after forward selection (Table S7), which suggest a strong influence of seasonality of temperature and precipitation gradients related to genome-wide diversity of native pecan.

Using the 12 forward-selected bioclimatic variables, we constructed pRDA models to disentangle the drivers of environment, geography, and demography on genetic variation across the range of native pecan. The combined model accounting for neutral genetic structure, environment, and geography had the best fit, explaining 11.97% of the total variation in the genetic data (p < 0.001); 44.0% of this explainable variance could be attributed to variation in bioclimate of origin alone (Table S8). Although this model illustrates a strong contribution of isolation by distance (IBD, 7.13% of explainable variance) and underlying population structure (24.88% of explainable variance) to the patterns of genetic diversity with respect to environment of origin, differences in environmental origin also clearly shape genetic variation in pecan (44.0% of explainable variance; 5.3% of total variance).

To identify loci that reflect the effect of environment on shaping genetic variation while accounting for the effects of population structure, we constructed a pRDA model including the 12 forward-selected bioclimatic variables, and the first three principal components of population structure as covariates. This approach revealed 446 variants significantly associated with bioclimate of origin (Fig. 2A) that were well-distributed throughout the genome (Fig. S7). The first axis of the pRDA accounted for 13.18% of variation and primarily described differences in precipitation patterns and seasonality of temperature. The second axis (10.28% variation described) was most related to mean temperature of the wettest quarter and precipitation seasonality. Using these 446 variants, which reflect – or more likely are linked to – variation related to environmental adaptation, we constructed an “adaptive RDA” model to represent contemporary genotype-environment associations (GEAs) (hereafter, “RDA-constructed GEA”).

**Figure 2.**
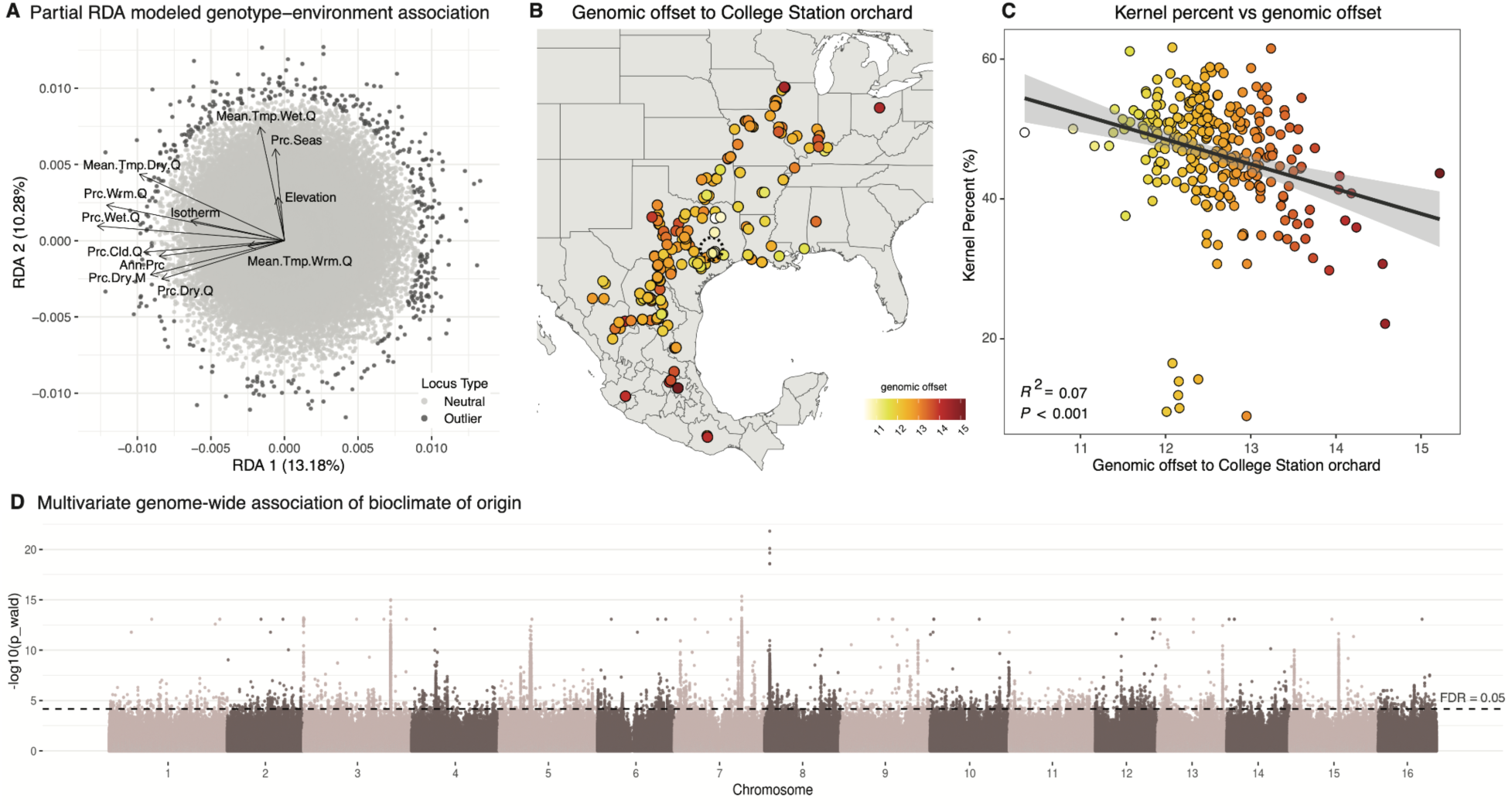
Genotype-environment associations capture signals of adaptation in native pecan. **A.** Biplot of partial RDA-constructed GEA, showing the projection of loci (points) and environmental variables (arrows) along the first two axes. **B.** Genomic offset of native pecan genotypes relative to the College Station common environment (large dashed black circle). Points are the source locations of native pecan genotypes and are colored by the genomic offset, calculated as the difference between observed and expected (predicted) optimal allele frequencies at the common garden. **C.** Relationship between kernel percent (a measure of fitness, collected in the College Station common environment) and predicted genomic offset to the College Station orchard. Points colored by predicted genomic offset to the College Station common environment. **D.** Manhattan plot for the multivariate genome-wide association with bioclimate of origin.

We validated our RDA-constructed GEA model’s ability to capture real trends in pecan adaptation by predicting adaptation to a common environment in which kernel weight was measured for 288 native pecan samples (Fig. 2B). Kernel weight is related to germination success, and higher kernel weight can be used as a measure of seed quality and fitness (Reid 1995; Thompson and Grauke 2003). For each sample in the common garden, we calculated a genomic offset to the expected genotype-environment association at the common garden (*Methods: Calculation of genomic offset*). Genomic offset is a measure of maladaptation based on genetic distance between a genotype and the optimal genotype-environment association of some target environment (Reviewed in (Rellstab, Dauphin, and Exposito-Alonso 2021)). Indeed, pecan samples with a lower genomic offset to the common garden had higher kernel percent (Fig. 2C, Fig. S8) and local genotypes had the lowest genomic offset to the conditions of the common garden (Fig. 2B, Fig. S8), suggesting that the set of variants included in the RDA-constructed GEA capture a signal of adaptation to local environments in native pecan.

### Dissecting markers linked to environmental gradients

To confirm that the identified environmentally associated loci were not sensitive to the RDA approach, we used the same set of 300k SNPs used in pRDA construction to implement the linear mixed model (LMM) GEMMA (X. Zhou and Stephens 2012) separately for elevation and the top 5 most predictive bioclimatic variables. There is an enrichment of RDA-identified SNPs (110 of 446; 24.7%) within the sites identified as highly significant across the univariate environmental GWAS (eGWAS) models (Fig. S9; P-value < 0.001) confirming that both methods capture some similar environmental associations, and also suggesting that RDA approaches may capture integrative signals that are missed when testing for associations with a single bioclimatic variable (Forester et al. 2018).

Given the distribution of RDA-identified variants, suggesting the set of sites capture genome-wide adaptive signals, we additionally performed a multivariate GWAS using a larger set of 10,619,337 SNPs and the top 5 most predictive bioclimatic variables to identify specific loci associated with environmental adaptation. The multivariate eGWAS identified significant associations at 14,604 SNPs (Fig. 2D). Despite differences in the SNP set size used in the multivariate eGWAS and RDA-identification of significant variants, the distance of adaptive RDA-identified variants to eGWAS associations is significantly closer than random sites used as eGWAS inputs (two sample Kolmogorov–Smirnov test D = 0.31624, *p* < 2.2 x 10^-16^; Fig. S10), with 23.1% (103 out of 446) of the outliers included in the RDA-constructed GEA occurring within 50kb of a significant multivariate eGWAS site. Thus, RDA-identified variants were often located near loci detected in eGWAS, supporting their putative role or linkage to sites involved in adaptive variation.

Multivariate eGWAS significant variants occurred within 414 gene models and had the top GO enrichment annotations of green leaf volatile (GLVs) biosynthetic process and response to karrikin. Both GLVs and karrikin have been linked to differences in abiotic and biotic stress response (Fang et al. 2023; Ameye et al. 2018; Cofer, Engelberth, and Engelberth 2018), consistent with the substantial variation in drought, temperature, and disease pressure across the range of pecan. Therefore these enrichments suggest they may have important roles in environmental adaptation in pecan.

### Current and Future Adaptation in Pecan

A goal in both evolutionary biology and plant breeding is to predict which genotypes are most vulnerable to changes in the environment and, in the case of environment mismatches, which genotypes are best matched to specific future conditions. Using the RDA-constructed GEA, we calculated two different estimations of genomic offset (*Methods: Calculation of genomic offset*), a measure of maladaptation, and identified the best matched genotypes to specific environment conditions. Our use of the RDA-constructed GEA provides an integrative representation of genomic-environmental associations (Forester et al. 2018), modeling how putatively adaptive diversity is distributed across environmental gradients and capturing long-term, multilocus signals. Furthermore, recent work by (Lind et al. 2024) suggests that relying solely on candidate adaptive loci may limit predictions of maladaptation, as such models may overlook patterns of neutral variation, gene flow, demographic history, and polygenic effects, all of which influence evolutionary responses.

For calculations of genomic offset, briefly, the first method calculates genomic offsets based on deviations of “loadings” in RDA-constructed genotype-environmental space (Capblancq and Forester 2021). The second method uses the RDA-constructed GEA to predict allele frequencies under a novel environment and calculates genomic offset as the Euclidean distance between observed and predicted frequencies (Gamba et al. 2025). Both methods capture important aspects of adaptive dynamics, but one measures how much genotypes deviate from the RDA-constrained multivariate adaptive space, while the other directly predicts allele frequencies using the fit RDA-constructed GEA for novel environmental conditions. Because approaches that measure maladaptation through calculation of genomic offset are still being refined, we applied both calculations to provide complementary insights.

We predicted genomic offset across the range of georeferenced native pecan under two future scenarios (SSP3–7.0 and SSP5–8.5). Spatial patterns of genomic offset, representing predicted maladaptation, varied between methods. Genomic offsets based on deviations from loadings of the RDA-constructed GEA identified the greatest predicted maladaptation in the Central and Southern Mexican populations and some samples from Texas (Fig. 3A), which is strongly correlated to environmental change (Fig. 3B) particularly decreases in annual precipitation (Fig. 3G). In contrast, genomic offsets based on deviations in predicted and observed allele frequencies identified Northern United States and Central Texas populations as having the highest genomic offsets (Fig. 3D) and were not as correlated to deviations in specific aspects of the bioclimate (Fig. 3G) though there is a relationship to seasonality of precipitation (Fig. 3E).

**Figure 3.**
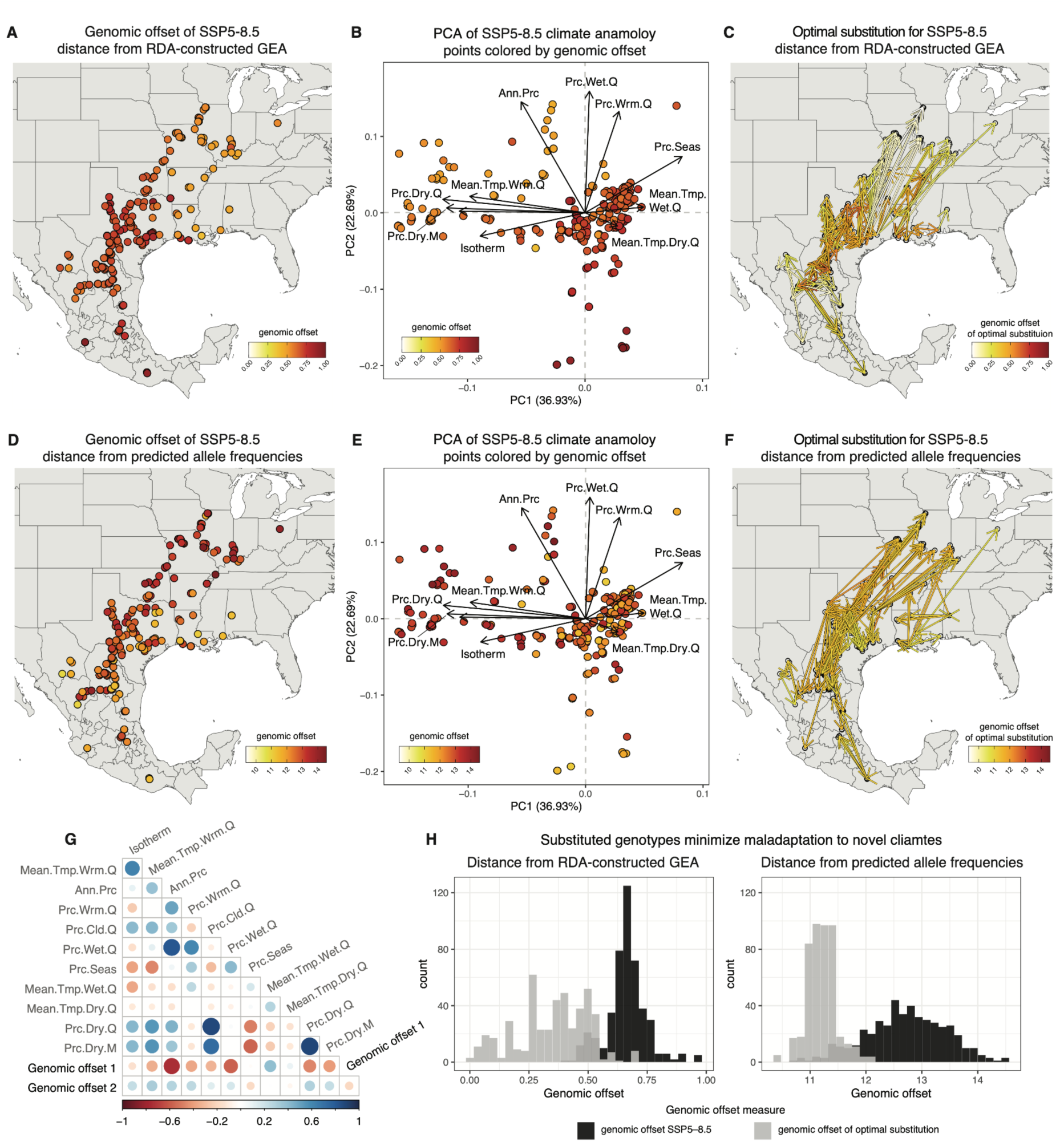
Leveraging genotype-environment associations (GEAs) to predict maladaptation and identify optimal genotypes for assisted migration. **A-C.** Genomic offset calculated from deviations in the loadings of RDA-constructed GEA space. **D-F.** Genomic offset calculated from differences in predicted and observed allele frequencies. **A,D:** Genomic offset for native pecan genotypes under the SSP5-8.5 emission scenario for their source location. **B,E.** PCA of environmental variable change at source locations of native pecan genotypes for the SSP5-8.5 environment anomaly. Points represent pecan samples and are colored by genomic offset. **C,F:** Optimal substitution, defined as the pecan genotype that minimizes genomic offset to the novel environment under the SSP5-8.5 emission scenario. For each target location, arrows connect the optimal source genotype to the migration destination (arrowhead) and are colored by the remaining genomic offset of a substitution**. G.** Correlation plot comparing environmental variables included in the climate anomaly PCA with genomic offset estimates from the two approaches: distance from RDA-constricted GEA (genomic offset 1) and distance from predicted allele-frequencies (genomic offset 2). **H.** Histograms of genomic offset values from both calculations before (black) and after (gray) optimal substitutions.

We next leveraged our GEA models to identify the native pecan samples that minimize genomic offset to, and therefore represent the pecan genotype best matched to, environmental conditions at locations across the range for current conditions and two future scenarios (Fig. 3; Fig. S11). Despite discrepancies in predicted range-wide genomic offsets under future emission scenarios between the two methods used to calculate genomic offset, we see similar patterns in the genotypes that are selected as optimal substitutions for future emissions across both methods (Fig. 3C, Fig. S11). For the most extreme emission scenario (SSP5-8.5), pecan genotypes that minimize maladaptation in the Northern portion of the range come from present-day locations that are farther South (Texas). Further, though no single genotype consistently minimized the genomic offset to future conditions across the range, the identification of genotypes that are optimal for substitutions to multiple nearby migration destinations suggest contemporary pecan genotypes that may provide valuable sources of resilience. For almost every location, existing genetic diversity is predicted to minimize genomic offset to future conditions (99.57% RDA loadings-based; 99.36% predict-based), suggesting that assisted migration of contemporary genotypes may improve the resilience of populations (Fig. 3H). However, after selecting best-fit genotypes for future environments, high quality substitutions that minimize genomic offset (RDA-loadings based prediction) are most common for the Northern portion of the range (Fig. 3C). In contrast, substitutions to central Mexico and parts of Texas have higher remaining genomic offset, suggesting limited diversity among contemporary pecan genotypes suited to these conditions and suggesting these regions as particularly vulnerable to future environmental change.

Pecan breeding has traditionally used cultivars derived from local, wild trees (Grauke, Wood, and Harris 2016), in part because they are adapted to the local environmental conditions. This is evident in the genetic variation contained in the breeding genotypes in this data set, where almost all USA breeding material is within the two gene pools predominantly found in the USA (Fig. 4A). Similarly, our offset estimates show that local genotypes are the best adapted for locations of commercial pecan production (Fig 4B, D). However, pecan trees live for more than 80 years, and our results indicate that genotypes with alleles adapted to a location may not be best adapted to that same location in less than the lifespan of a tree (Fig. 3). For several locations of commercial production, genomic offset predictions indicate that local genotypes will have a higher degree of maladaptation to environment conditions projected for within 75 years from present (Fig 4C, E), less than the lifetime of a pecan tree, and substituting non-local variation may be a management strategy for increasing pecan productivity. Since the USA breeding material primarily contains just a subset of species-wide genetic diversity (Fig 4A), capturing adapted alleles from other gene pools may be a productive avenue for improving certain traits. Breeding decisions include many considerations, including consumer and grower preferences in addition to selection of adaptive traits to improve productivity. Information about the adaptive fit and adaptive alleles will be valuable information as breeders and growers make decisions about what varieties to develop and where to grow them in the future.

**Figure 4.**
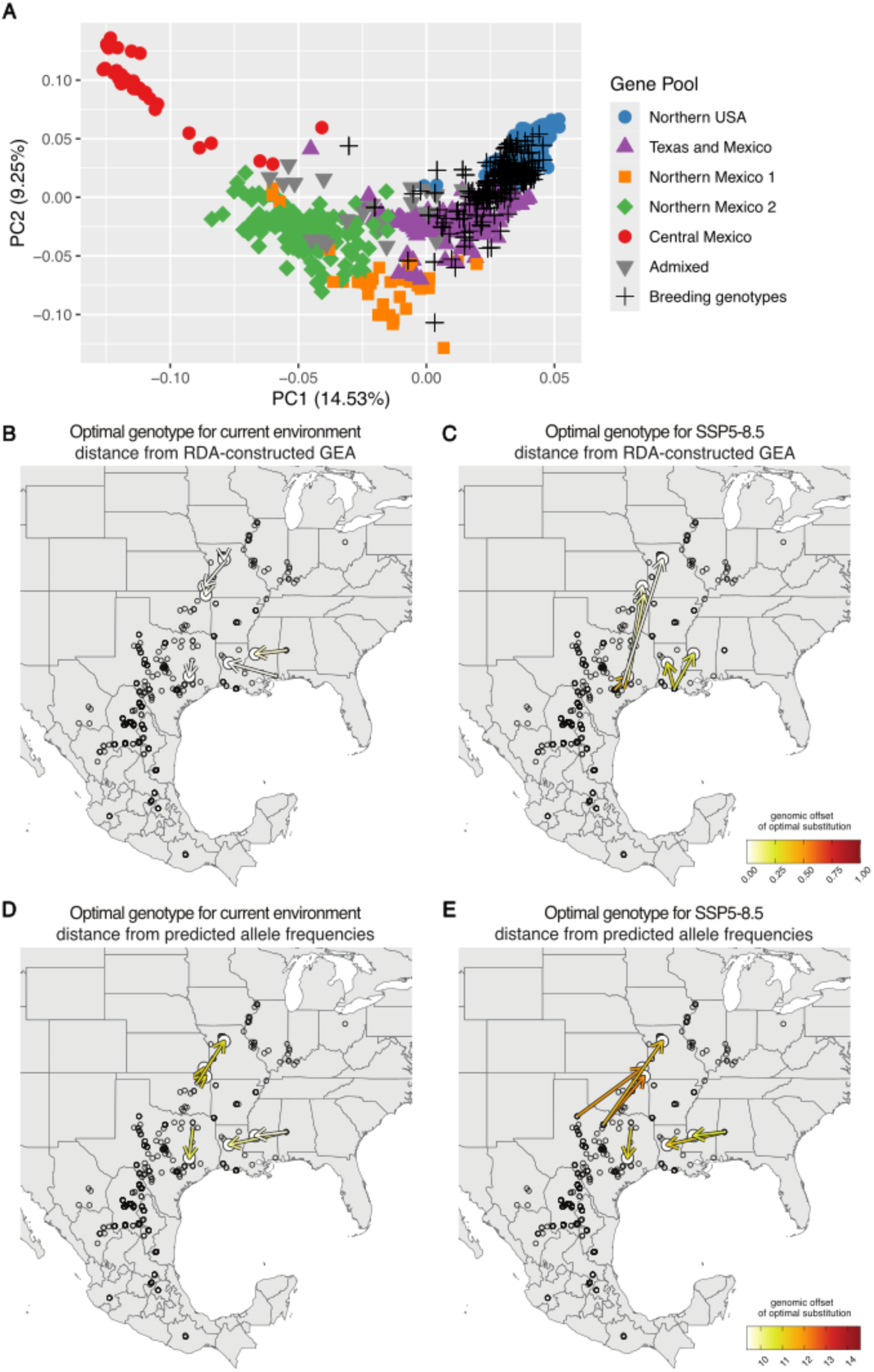
Genome offset predictions point to the benefit of using expanded genetic variation for future pecan production. **A.** Principal component analysis of 466 native and 217 breeding pecan genotypes. Values for breeding material genotypes were projected onto principal component axes calculated using the native samples to capture the relationship of breeding genotypes with the native gene pools. **B-E** Optimal native genotype predictions for 6 commercial growing locations within the native range pecan. Arrows connect the optimal source genotype to the destination (arrowhead) and are colored by the remaining genomic offset of a substitution. **B,D:** Optimal genotype prediction for current environmental conditions. **C,E:** Optimal genotype predictions for SSP5-8.5 environment anomaly. **B,C:** Genomic offsets calculated as distance from RDA-constricted GEA. **D,E:** Genomic offsets calculated as differences in predicted and observed allele frequencies

## Methods

### Genome Assembly

We sequenced *Carya illinoinensis* (var. ‘Oaxaca’ 87MX3-2.11) using a whole genome shotgun sequencing strategy and standard sequencing protocols. Sequencing reads were collected using Illumina and PACBIO platforms. Illumina and PACBIO reads were sequenced at the HudsonAlpha Institute in Huntsville, Alabama. Illumina reads were sequenced using the Illumina NovoSeq6000 platform, and the PACBIO reads were sequenced using the REVIO platform using the Circular Consensus Sequencing (CCS) protocol. Two 400bp insert 2x250 Illumina fragment libraries (195.21x per haplotype) was sequenced along with one 2x150 HiC library (120.70x per haplotype) (Table S9). Prior to assembly, Illumina fragment reads were screened for PhiX contamination. Reads composed of >95% simple sequence were removed. Illumina reads <50bp after trimming for adapter and quality (q<20) were removed. The final read set consists of 1,789,923,358 reads for a total of 195.21x of high-quality Illumina bases. For the PACBIO sequencing, the total raw sequence yield was 34.70 Gb, with a total coverage of 25.71x per haplotype (Table S10).

The version 2.0 HAP1/HAP2 assemblies were generated by assembling the 5,092,472 PACBIO CCS reads (25.71x per haplotype) using the HiFiAsm+HIC assembler (Cheng et al. 2021) and subsequently polished using RACON (Vaser et al. 2017). This produced initial assemblies of both haplotypes. The initial HAP1 assembly consisted of 161 scaffolds (161 contigs), with a contig N50 of 14.1 Mb, and a total genome size of 671.5 Mb (Table S11). The initial HAP2 assembly consisted of 158 scaffolds (158 contigs), with a contig N50 of 12.1 Mb, and a total genome size of 674.7 Mb (Table S12).

Hi-C Illumina reads from *C. illinoinensis* (var. ‘Oaxaca’ 87MX3-2.11), were separately aligned to the HAP1 and HAP2 contig sets with Juicer (Durand et al. 2016), and chromosome scale scaffolding was performed with 3D-DNA (Dudchenko et al. 2017). No misjoins were identified in the HAP1 or HAP2 assemblies. The contigs were then oriented, ordered, and joined together into 16 chromosomes per haplotype using the HiC data. A total of 105 joins were applied to the HAP1 assembly, and 90 joins for the HAP2 assembly. Chromosomes were numbered using the version 1.0 *C. illinoinensis* cv. Pawnee assembly available from Phytozome (Goodstein et al. 2012). Each chromosome join is padded with 10,000 Ns. Contigs terminating in significant telomeric sequence were identified using the (TTTAGGG)_n_ repeat, and care was taken to make sure that they were properly oriented in the production assembly. The chloroplast genome was assembled using OatK (C. Zhou et al. 2025). The remaining scaffolds were screened against bacterial proteins, organelle sequences, GenBank nr and removed if found to be a contaminant. After forming the chromosomes, it was observed that some small (<20Kb) redundant sequences were present on adjacent contig ends within chromosomes. To resolve this issue, adjacent contig ends were aligned to one another using BLAT (Kent 2002), and duplicate sequences were collapsed to close the gap between them. A total of 21 adjacent contig pairs were collapsed in the HAP1 assembly and 21 in the HAP2 assembly.

Finally, homozygous SNPs and INDELs were corrected in the HAP1 and HAP2 releases using ∼55.5x of Illumina reads (2x150, 400bp insert) by aligning the reads using bwa mem (Li 2013) and identifying homozygous SNPs and INDELs with the GATK’s UnifiedGenotyper tool (McKenna et al. 2010). A total of 389 homozygous SNPs and 5,200 homozygous INDELs were corrected in the HAP1 release, while a total of 301 homozygous SNPs and 4,920 homozygous INDELs were corrected in the HAP2 release. The final version 2.0 HAP1 release contains 670.0 Mb of sequence, consisting of 100 contigs with a contig N50 of 14.0 Mb and a total of 100% of assembled bases integrated into chromosomes. The final version 2.0 HAP2 release contains 659.0 Mb of sequence, consisting of 86 contigs with a contig N50 of 15.9 Mb and a total of 99.98% of assembled bases in chromosomes.

Completeness of the euchromatic portion of the version 2.0 assemblies was assessed by aligning the existing version 1.0 *C. illinoinensis* cv. Lakota annotated primary transcripts to the version 2.0 releases. The aim of the completeness analysis is to obtain a measure of completeness of the assembly, rather than a comprehensive examination of gene space. We retained genes that aligned at greater than 90% identity and 85% coverage. The screened alignments indicate that 98.22% of the primary transcripts aligned to the HAP1 release, and 98.28% aligned to the HAP2 release.

### Genome Annotation

To annotate the phased Oaxaca.v2 genomes, we applied a spliced alignment-based liftover approach using coding DNA sequences (CDS) from publicly available unphased ‘Oaxaca’, ‘Pawnee’, ‘Elliott’, and ‘Lakota’ reference genomes (Lovell, Bentley, et al. 2021). GENESPACE (v1.3.1) (Lovell et al. 2022) was used to cluster protein sequences from these varieties into hierarchical orthogroups, and the longest representative protein from each orthogroup was selected. Corresponding CDS were extracted from reference annotations and aligned to the phased Oaxaca.v2 genomes using GSNAP (v2023-10-10, (Wu et al. 2016) with the options `--npaths=1, --no-chimeras,` and `--gff3-fasta-annotation=1`. The resulting GFF3 file was sorted using gff3sort (v1.0.0, (Zhu et al. 2017), and protein sequences were generated with gffread (v0.12.7, (Pertea and Pertea 2020), filtering out sequences shorter than 30 amino acids.

Transposable elements were annotated using EDTA v. 2.21 (Ou et al. 2019) and then using panEDTA (Ou et al. 2024). We first ran EDTA v. 2.2.1 on each haplotype individually, using coding sequence as additional evidence. We then constructed a shared TE library using panEDTA, and finally reannotated each haplotype using this unified library.

### Germplasm Collection

The genotypes used in this study come from *ex-situ* germplasm collections at the USDA-ARS National Plant Germplasm System (NPGS) - BRW worksites in Texas: the Brownwood worksite in Brown County, Texas, and the College Station worksite in Burleson County, Texas. Trees in this collection originate from two general types of propagative source material: wild-origin seed collected from across the geographic range of pecan or hickory species (primarily grown on their own roots, (Grauke et al. 2011; Grauke, Payne, and Wood 1989), or asexually propagated graftwood taken from pecan or hickory cultivars or native trees (primarily grafted on pecan seedling rootstocks). Approximately 450 named cultivars and 3,000 native seedlings are represented between both NPGS worksites. Trees grown from seeds collected from native populations are considered accurate representations of the local genetic variation since both parents are from the locality from which the seeds were collected.

### WGS Sequencing and Genotyping

Young tender spring leaf tissue was collected from mature pecan trees in the USDA-ARS NPGS – BRW worksites in College Station and Brownwood, TX in 2019 and 2021. For the first collection in 2019, young tender spring leaf tissue was flash-frozen in liquid nitrogen for 440 pecan accessions and stored at −80℃ for DNA extraction. DNA was extracted using a Qiagen Dneasy Plant Mini Kit (QIAGEN) following the manufacturer’s instructions. Illumina libraries were constructed using standard protocols (Illumina TruSeq PCRfree Catalog #21105962) and were sequenced using the Illumina NovoSeq 6000 platform. For the second collection in 2021, young tender spring leaf tissue was collected and lyophilized for 380 pecan and hickory trees. DNA was extracted using a Qiagen Dneasy Plant Mini Kit (QIAGEN) following the manufacturer’s instructions. Illumina libraries were constructed using standard protocols (Illumina TruSeq PCRfree Catalog #21105962) and sequenced on the Illumina HiSeq 4000 platform. Illumina libraries were mapped to the Oaxaca.v2_HAP1 assembly using bwa-mem2 (Vasimuddin et al. 2019) and duplicate reads were removed using picard “MarkDuplicates” (v.3.0.0, http://broadinstitute.github.io/picard). We use samtools “mpileup” (v1.17, (Danecek et al. 2021)) to generate a VCF file of variates followed by “mpileup2cns” in varscan (v2.4.4, (Koboldt et al. 2012) to call genotypes.

For GBS libraries, fresh leaves were sampled and stored at −20C until DNA was extracted using DNeasy Plant Mini Kit (QIAGEN). DNA was processed following (Baird et al. 2008). DNA was digested using the restriction enzymes *Eco*RI and *Bgl*II and divided into 4 total sequencing libraries. Illumina libraries were prepared using standard protocols and sequenced at Clockmics (Osaka, Japan) on an Illumina HiSeqTM and HiSeq 2000. Demultiplexed libraries were mapped to the Oaxaca.v2_HAP1 assembly using bwa-mem2 (Vasimuddin et al. 2019) and the VCF of variants was generated with samtools “mpileup” (v1.17, (Danecek et al. 2021)) followed by “mpileup2cns” in varscan (v2.4.4, (Koboldt et al. 2012) to call genotypes. Raw sequences are registered at the NCBI Short Read Archive (SRA) under BioProject ID: PRJNA614625.

### Whole Genome Alignment and Inversion Analysis

We used minimap2 (Li 2021) for whole genome alignments of Oaxaca.v1, Oaxaca.v2_HAP1, and Oaxaca.v2_HAP2. SyRI (Goel et al. 2019) was used for inversion detection and to compare the amount of sequence in pericentromeric regions, defined as regions with high repetitive content and <1.0% gene sequence.

### Population genetic analysis

To assay diversity and population structure in pecan we first used plink (v1.9, (Purcell et al. 2007) to perform principal component analysis (PCA) with genotypes from native pecan populations using more than 6 million SNPs that were LD-pruned and with minor allele frequency > 0.007. We then used bcftools (Danecek et al. 2021) to randomly select 500k genome-wide SNPs from the native pecan genotypes and used that subset of SNPs with ADMIXTURE (Alexander, Novembre, and Lange 2009) to test the fit of the genotype data to K=2 through K=12 genetic populations. We used the cross-validation (“--cv”) option to select the best fit of the data. We calculated IBD-based genetic distance between native pecan samples with plink (v1.9, (Purcell et al. 2007) and used pairwise distance values to evaluate relative levels and variation of nucleotide diversity between gene pools. Diversity patterns using pairwise genetic distance and window-based pi calculations in pixy ((Korunes and Samuk 2021); see below) show similar patterns across gene pools, and we present the pairwise genetic distance results to better represent the variation in how individuals contribute to the gene pool level of nucleotide diversity (Fig. S12). Heterozygosity was calculated using plink (v1.9, (Purcell et al. 2007) for whole genome sequencing (WGS) and GBS genotypes, and mean heterozygosity of samples with the same geo-reference information and same genotyping method was used for comparisons of heterozygosity and latitude. We also used pixy (Korunes and Samuk 2021) to measure nucleotide diversity, Watterson’s theta, and Fst across natural pecan samples and within each gene pool.

We used dadi (Gutenkunst et al. 2009) to estimate the divergence times of gene pools. For each scenario, we tested models for divergence with no migration and with or without exponential growth using both standard and log-based optimization methods. We ran five replicates of each model and optimization method and summarized results using the output with the highest likelihood from each combination of model and optimization method. The input genotypes were all SNPs on Chr01 after repetitive regions were removed. To estimate divergence time for the three most differentiated gene pools, Central Mexico, North Mexico 1, and Northern USA, we generated frequency spectra using 20 samples from each of the gene pools with 80% or more ancestry based on the results from ADMIXTURE. The sample numbers for this scenario were limited by the number of samples with North Mexico 1 ancestry. For the gene pools with the lowest differentiation, North Mexico 2, Texas-Mexico, and Northern USA, we generated frequency spectra using 50 samples from each gene pool.

To estimate patterns of migration in native pecan, we used Fast Estimation of Effective Migration Surfaces (FEEMS) (Marcus et al. 2021; Petkova, Novembre, and Stephens 2016) and modeled the estimated effective migration surface for both the GBS and WGS datasets. To estimate the GBS effective migration surface, we used 19,944 LD-pruned SNPs filtered for MAF > 0.01 of *n* = 581 samples georeferenced to 206 unique locations. To estimate the WGS effective migration surface, we used 50,038 LD-pruned SNPs filtered for MAF > 0.01 of *n* = 466 samples georeferenced to 266 unique locations. For the estimation of lambda, the smoothing parameter, we used the suggested cross validation method for each migration surface separately (Marcus et al. 2021; Petkova, Novembre, and Stephens 2016).

### Environment data

For the set of georeferenced native pecan samples, we extracted the 19 standard bioclim variables from WorldClim V2.1 (Fick and Hijmans 2017). Data are representative of long-term historical averages (1970–2000) in environment and were generated at 30 sec spatial resolution. Future environment scenarios were projected using the Coupled Model Intercomparison Project (CMIP6) and shared socioeconomic pathways (SSPs) developed by the IPCC. This phase of this model integrates new emission and land-use scenarios compared with previous models (Eyring et al. 2016; O’Neill et al. 2016; Stouffer et al. 2017; Wyser et al. 2020). We further gathered WorldClim v2.1 bias-corrected future bioclimate data averaged from five CMIP6 models (ACCESS-CM2, CMCC-ESM2, IPSL-CM6A-LR, MIROC6, MPI-ESM1-2-HR) for the time period 2081-2100 and two emission scenarios (SSP3-7.0 and SSP5-8.5 (Gidden et al. 2019)) at 30 sec spatial resolution.

### Redundancy analysis

To identify loci with significant variation explained by environmental variation, which we hypothesize may capture signals of native pecan local adaptation, we used partial redundancy analysis (pRDA) methods as described in (Capblancq and Forester 2021). Redundancy analysis captures linear relationships between genetic markers and explanatory variables, modeling this variation along orthogonal axes that summarize patterns of covariation between genetic and environmental data. We used forward selection (R/vegan::ordistep(), (Oksanen et al. 2022)), to determine and exclude highly collinear or non-predictive bioclimate variables from pRDA construction. pRDA models were built using population allele frequencies (population defined as samples from the same geocoordinates) of 300,000 LD-pruned SNPs, filtered for MAF > 0.05 as response variables and the explanatory variables of bioclimate and elevation, while accounting for the first three PCs of population structure as covariates. Our goal here was to account for population structure without overcorrecting for geographic signals, which are likely correlated to the environment (high false negative rate when explicitly correcting for geography (Forester et al. 2018)). Following methods described in (Capblancq et al. 2018), we identified SNPs as candidates under selection (RDA-identified variants) through the *radapt* function. Briefly, outlier loci are selected based on the extremeness of their loading along a Mahalanobis distance distribution, calculated between each marker (SNP) and the center of the first two pRDA axes. Each SNP is given a *p*-value derived from this distance and corrected for the inflation factor using a chi-squared distribution with two degrees of freedom. From this methodology, we identified 446 putatively environmentally adaptive SNPs which were used in construction of the adaptive RDA models for further exploration of current genotype-environment associations (GEAs).

### Environmental GWAS

To assess whether the top loci selected by pRDA are unique to the method, we further implemented Genome-wide Efficient Mixed Model Association (GEMMA; (X. Zhou and Stephens 2012) and compared the significant loci as identified by GEMMA and pRDA. We additionally implemented a multivariate GEMMA model for the top 5 most predictive forward-selected bioclimatic variables.

### Calculation of genomic offset

Genomic offset (sometimes referred to as genomic vulnerability) is one metric used to characterize maladaptation with a genomic context (reviewed in ref. (Rellstab, Dauphin, and Exposito-Alonso 2021)). The distance between current and expected genotype-environment associations under novel environments is representative of the genomic offset, or the genetic shift required in a population to adapt to the future environment. Comparing the current genotype-environment association captured by RDA models, and the projected genotype-environment association for some target condition (ex: common phenotyping environment, future emission scenarios, substitution locations) we made several measurements of genomic offset to summarize predicted maladaptation. For all genomic offset calculations, we used two approaches. Both approaches calculate an Euclidean distance between the accession’s RDA-modeled GEA (representing current genotype-environment relationships) and the expected genomic composition for the target environment (representing the projected optimal genotype-environment relationship for the target location). The first method calculates expected genomic composition from deviations in the constrained RDA ordination space (extrapolated from RDA loadings; (Capblancq and Forester 2021)) and the second quantifies the difference between observed and expected allele frequencies (calculated using R/vegan::predict; (Gamba et al. 2025)) under the future environment. Both measures of genomic offset are extrapolated from current genotype-environment relationships represented in the adaptive RDA model (weighted by the contribution of loci included in the model to represent current adaptation) and indicate what amount of genetic change would be required for adaptation to the common environment. For the analysis of common garden kernel phenotypes, we calculated genomic offset for the 288 native pecan samples with phenotype data (see Methods: Kernel phenotypes) as predicted to the common environment (30.525 N, 96.4238 W) using the two methods described above.

We extended our RDA-constructed GEA models to predict genomic offset in native pecan under two emission scenarios: SSP3-7.0 and SSP5-8.5. Here, genomic offset was calculated as the Euclidean distance between the current RDA-predicted GEA and for the future projected GEA of each emission scenario. We again calculated genomic offset using the two methods as described above.

### Pecan nut kernel phenotypes

To confirm our RDA-constructed GEA models captured real differences in current pecan adaptation across the range of environments to which native pecan is local, we used phenotypic data of a multiyear-multitree average of kernel weight and kernel percent for 288 native pecans grown in the National Plant Germplasm System - BRW repository in College Station, TX, USA and/or Brownwood, TX, USA. These averages represent a median of 4 years and a mode of 1 year per accession. Kernel weight (g) was calculated using a destructive measurement of 5 nuts per year by cracking the nut with an Inertia Nut Cracker, carefully extracting the kernel from the shell, and weighing the contents in grams.

### Identification of pecan genotypes for novel environmental conditions

The existing genetic diversity of pecan may be a source of resilience to changing environments through assisted migration. To evaluate this potential, we identified the current pecan genotype with the lowest genomic offset to specific environmental conditions (range-wide and orchard sites under current and future emission scenarios), representing the genotype with the lowest predicted maladaptation based on current genotype-environment associations captured by the RDA-constructed GEA. Genomic offsets were calculated using both methods described in *Methods: Calculation of genomic offset.* For each method, the most optimal substitution was defined as the genotype with the minimum Euclidean distance to the target climate of the area (M McLaughlin et al. 2025). The remaining genomic offset represents the gap between the substituted genotype and the predicted genotype for adaptation to the projected conditions. High remaining genomic offset indicates that no existing genotype is well matched to the future climate of the location, suggesting potential vulnerability.

### Locations of Commercial Production

The locations of commercial production were identified as the locations of large commercial or research orchards in the 6 states within the native range of pecan with the highest annual commercial pecan production: Kansas, Louisiana, Mississippi, Missouri, Oklahoma, and Texas.

## Supporting information

Data Set 1

Data Set 2

## Data Accessibility

Data files will be available on the Pecan Toolbox website (https://pecantoolbox.nmsu.edu/). These include the assembly and annotation files for Oaxaca.v2_HAP1 and Oaxaca.v2_HAP2 and the genotypes used for this analysis as Variant Call Format (VCF) files. The VCF using GBS genotypes is available as SI Dataset 1. The Kernel phenotype data is available as SI Dataset 2. All raw sequencing reads have been deposited in the NCBI SRA database under BioProject accessions PRNJA680537, PRJNA1375741 and PRJNA1375742.

## Acknowledgements

The authors would like to acknowledge Dr. Bruce Wood and Dr. Jerry Payne for their role in establishing the pecan provenance collections that serve as the foundation of this study. We acknowledge the World Climate Research Programme for coordinating and promoting CMIP6, thank the climate modeling groups for their model output, and appreciate the Earth System Grid Federation (ESGF) and multiple funding agencies for archiving the data and providing access. We acknowledge Pecan Toolbox for hosting the data from this work for public access.

The work was also funded in part by U.S. Department of Agriculture National Institute of Food and Agriculture (SCRI-2016-51181-25408) “Coordinated Development of Genetic Tools for Pecan” and U.S. Department of Agriculture National Institute of Food and Agriculture (SCRI-2022-51181-38332) “Trees for the Future: Coordinated Development of Genetic Resources and Tools to Accelerate Breeding of Geographic Adapted Pecan Trees”, and by the U.S. Department of Agriculture – Agriculture Research Service National Programs through CRIS project 3091-21000-046-000-D (Crop Germplasm Research Unit, TX). The agency was not involved in the study design, collection, analysis, interpretation of data and the writing of this article. However, this manuscript was approved by the agency before submission for publication. This article reports on the results of the research only. This material is based upon work supported by the Center for Bioenergy Innovation (CBI), U.S. Department of Energy, Office of Science, Biological and Environmental Research Program under Award Number ERKP886.

## Notes

https://pecantoolbox.nmsu.edu/

### Competing Interest Statement

The authors have declared no competing interest.

