## Supporting Information for "Leveraging species-wide variation and patterns of adaptation to inform pecan crop improvement efforts"

#### Supplemental Figures

**Fig S1**

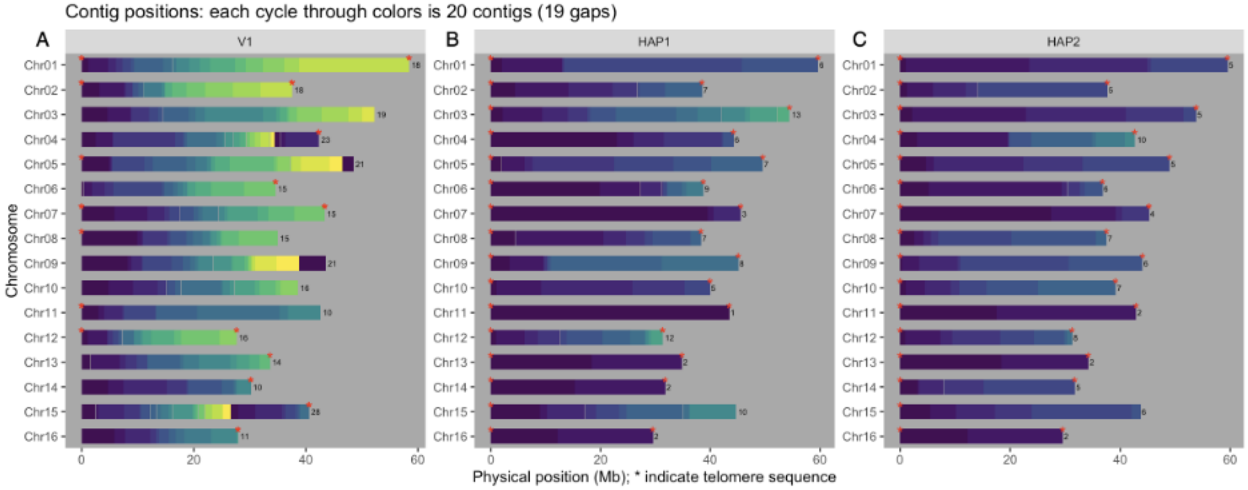

**Fig. S1: Contig Map of 'Oaxaca' genome assemblies**
Comparison of the number and length of contigs used to assemble each chromosome for the 'Oaxaca' V1 assembly (**A**) and the phased 'Oaxaca' V2 HAP1 (**B**) and HAP2 (**C**) assemblies. Red stars indicate the presence of identifiable telomeric sequence.

**Fig S2**

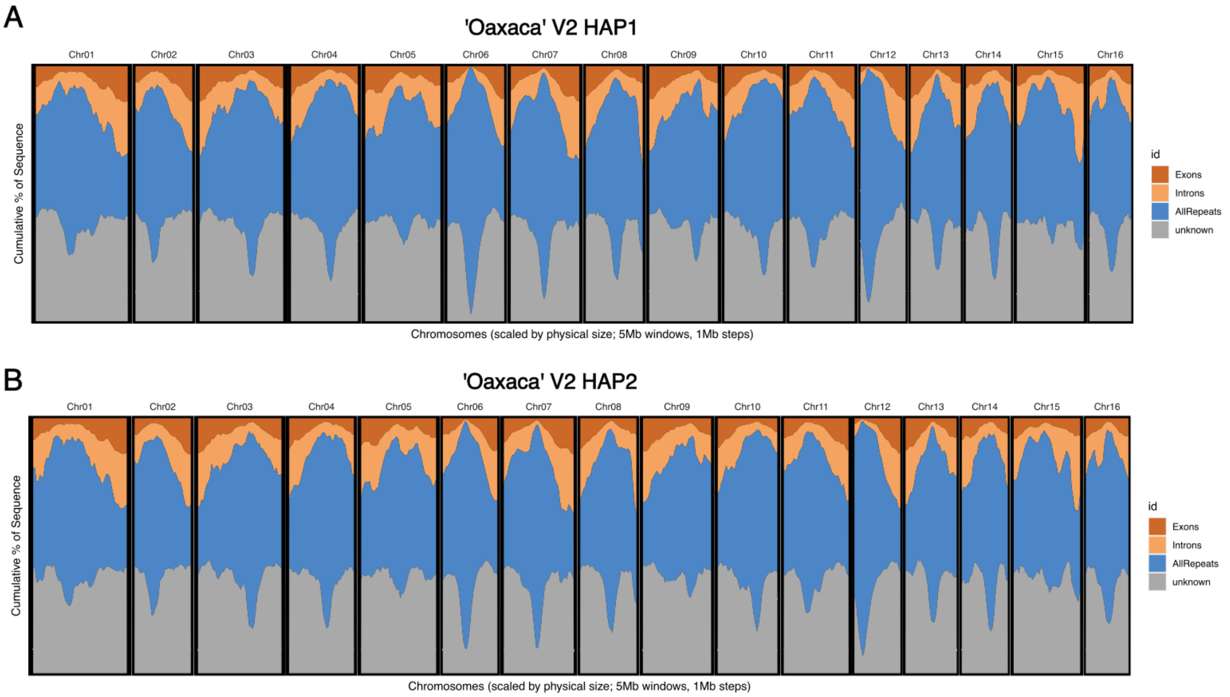

**Fig. S2: Gene and repeat density across 'Oaxaca' V2 Assemblies**
Density of exon, intron, repetitive, and unannotated sequence in 5Mb windows across the 'Oaxaca' V2 HAP1 (**A**) and HAP2 (**B**) assemblies.

**Fig S3**

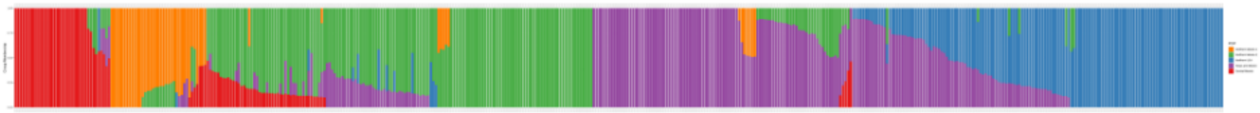

**Fig. S3: ADMIXTURE ancestry membership across native pecan for K=5**
Membership in the 5 pecan gene pools from K=5 ADMIXTURE results for the 466 native pecan samples genotyped using Illumina whole genome resequencing. Colors correspond to gene pool ancestry: Red: Central Mexico; Orange: North Mexico 1; Green: North Mexico 2; Purple: Texas and Mexico; Blue: Northern USA.

**Fig S4**

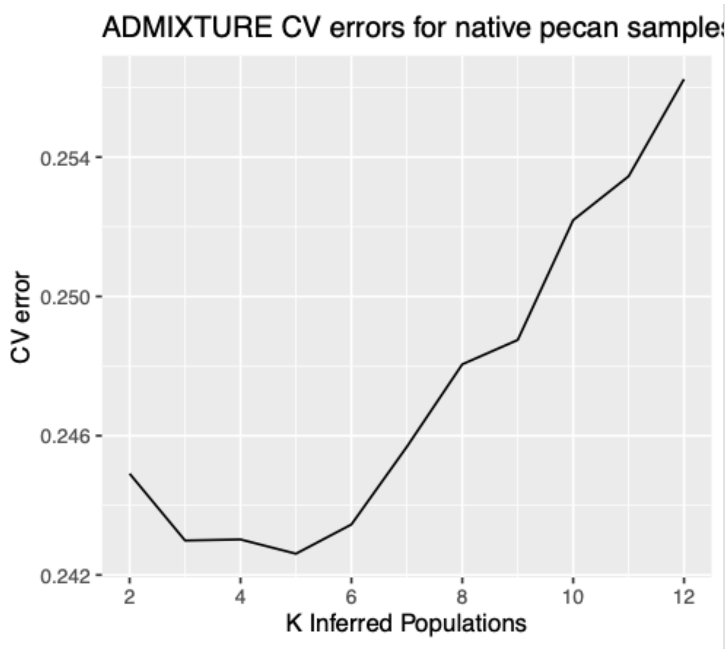

**Fig. S4: Cross-validation results for ADMIXTURE tests**
Cross validation results from ADMIXTURE tests of the number of populations ranging from 2 to 12 using the 466 native pecan samples genotyped using Illumina whole genome resequencing. K=5 has the lowest cross-validation error.

**Fig S5**

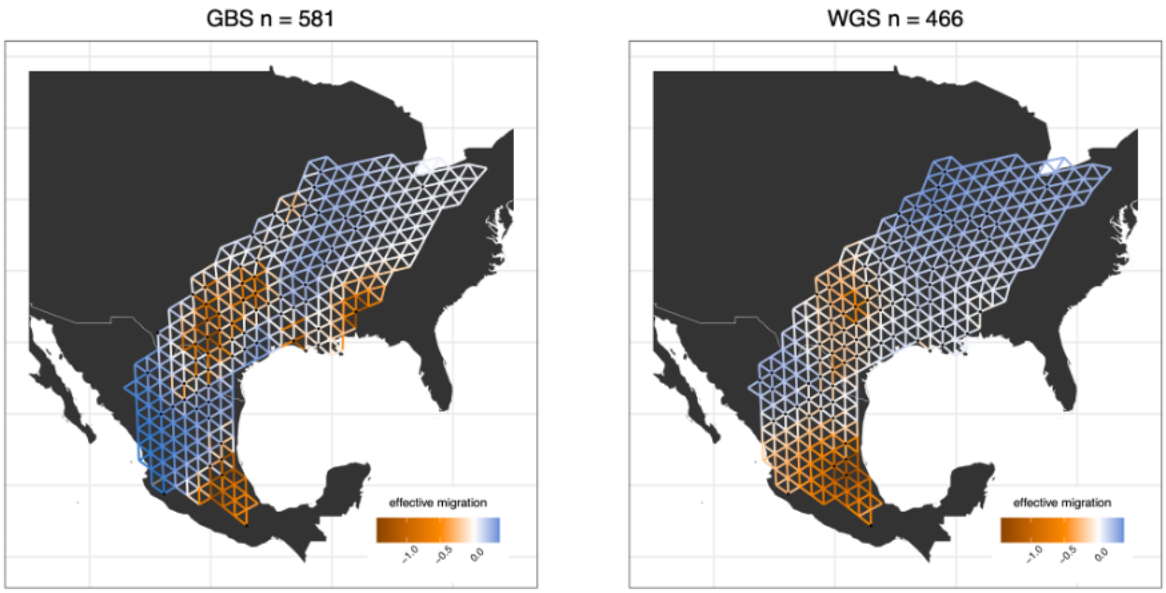

**Fig. S5: Estimated effective migration among native pecans.**
Estimated effective migration surfaces among native pecan populations calculated with FEEMS (Marcus et al. 2021) and using the GBS dataset (left, n=581) and WGS dataset (right, n=466). High gene flow (blue) and low gene flow (orange) are indicated by colored edges in the 'wire-

mesh' graph. Graph grid resolution corresponds to cell spacing (distance between mid-points of adjacent cells) of 110 km.

**Fig S6**

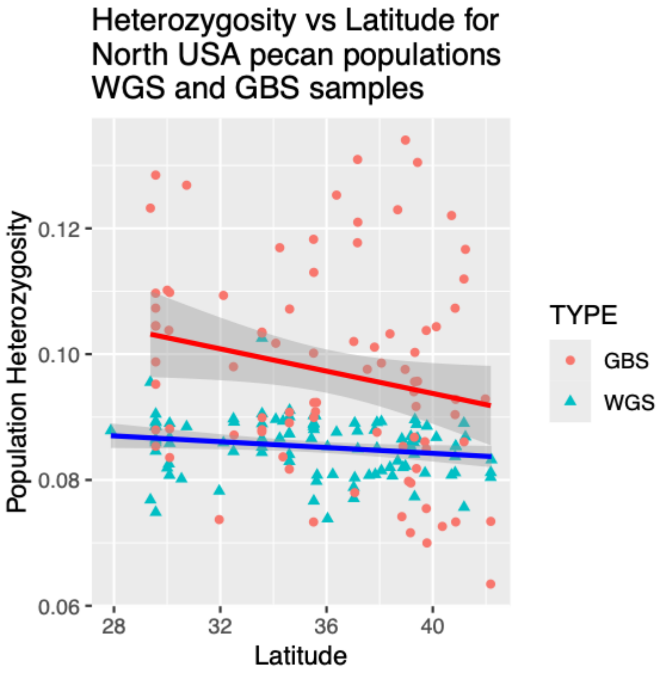

**Fig. S6: Latitudinal gradient in heterozygosity in Northern USA populations**
Relationship of latitude and heterozygosity in the Northern USA gene pool for both WGS and GBS genotypes. Heterozygosity was measured as mean heterozygosity of samples with the same geo-reference source information and same genotyping method.

**Fig S7**

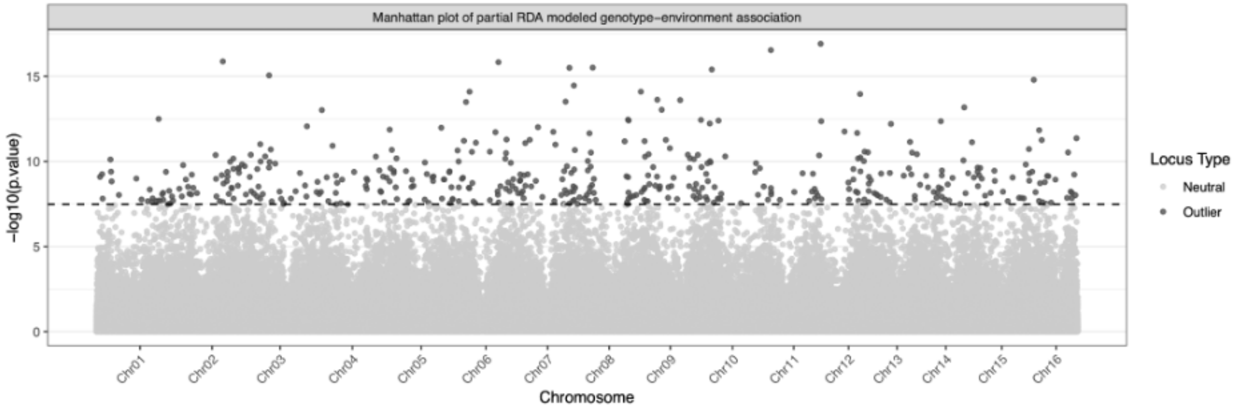

**Fig. S7: Genome-wide distribution of partial RDA (pRDA)-constructed genotype-environment association (GEA) sites.**

Location in 'Oaxaca' V2 HAP1 of the 300k SNPs used for pRDA construction and the significance of the pRDA-modeled association with bioclimate of origin plot as a "Manhattan plot". The same SNPs are represented as a “biplot” in Fig. 2A.

**Fig S8**

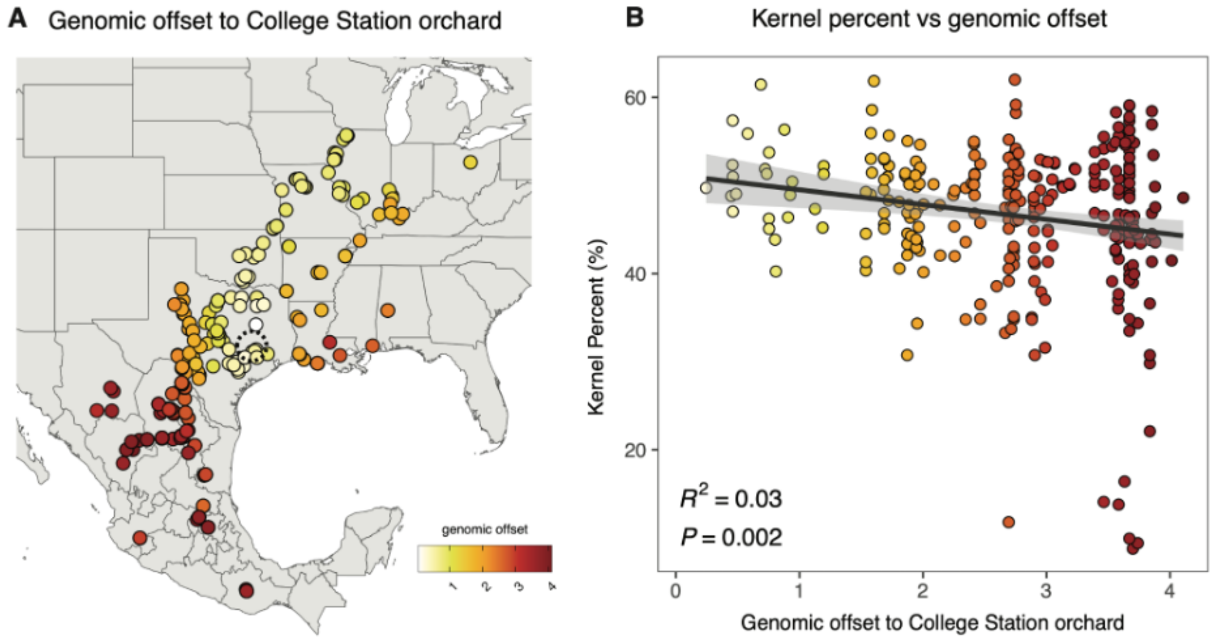

**Fig. S8: Genomic offset to common garden using RDA loadings**
**A.** Genomic offset to common garden in College Station, TX (dashed circle) for native pecan samples. Genomic offset calculation derived from loadings of RDA-constructed GEA. **B.** Kernel percent compared to genomic offset, as calculated in (A). **A,B** Similar to Fig 2B,C except using a different calculation of genomic offset.

**Fig S9**

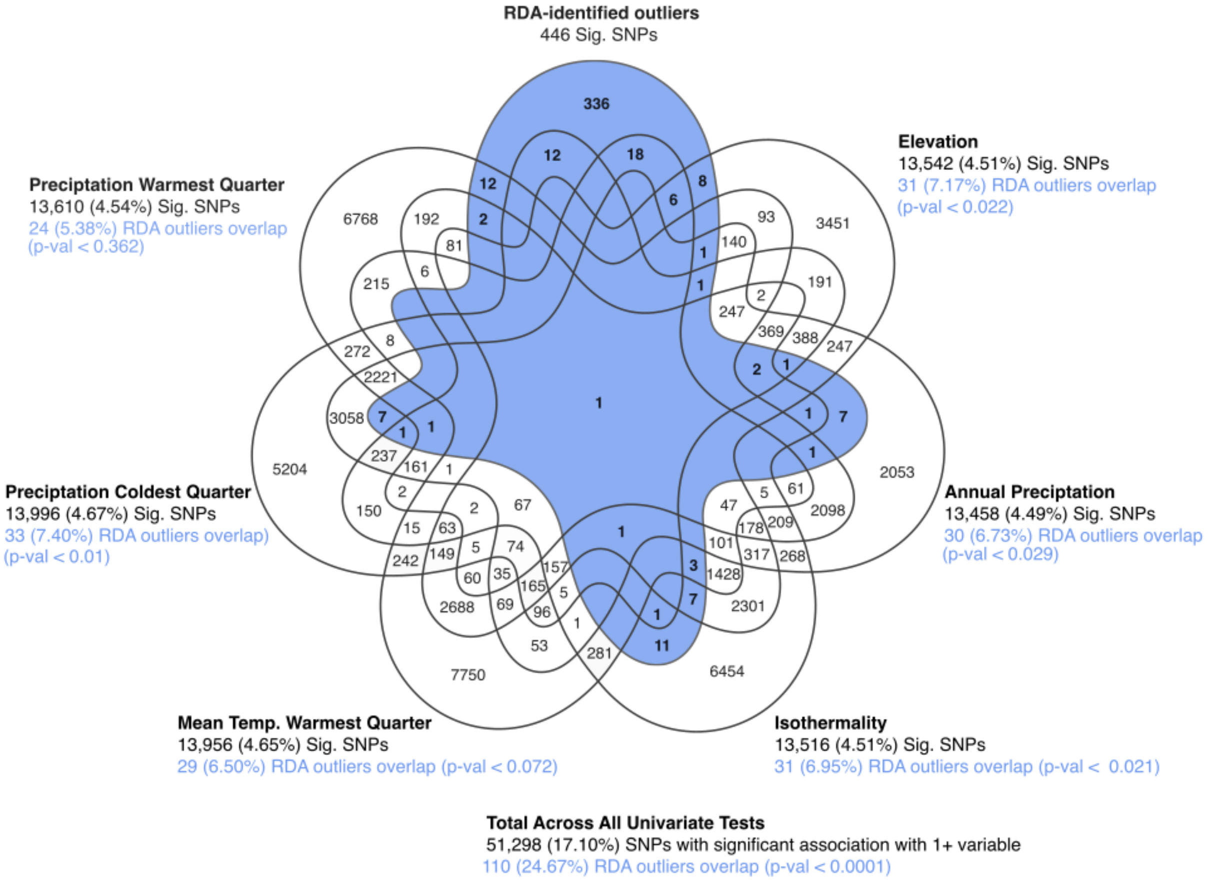

**Fig S9: Overlap of RDA outliers and univariate eGWAS SNPs**

Venn diagram showing the overlap of RDA-identified outlier SNPs and the SNPs with significant (raw p-value < 0.05) associations in univariate eGWAS. The number of SNPs with associations for each eGWAS model is in black. The number of RDA outliers that overlap with eGWAS associations is in blue; p-value is from a binomial test for enrichment. The eGWAS use the same set of 300,000 SNPs that were used to construct the pRDA and identify the 446 RDA outliers.

**Fig S10**

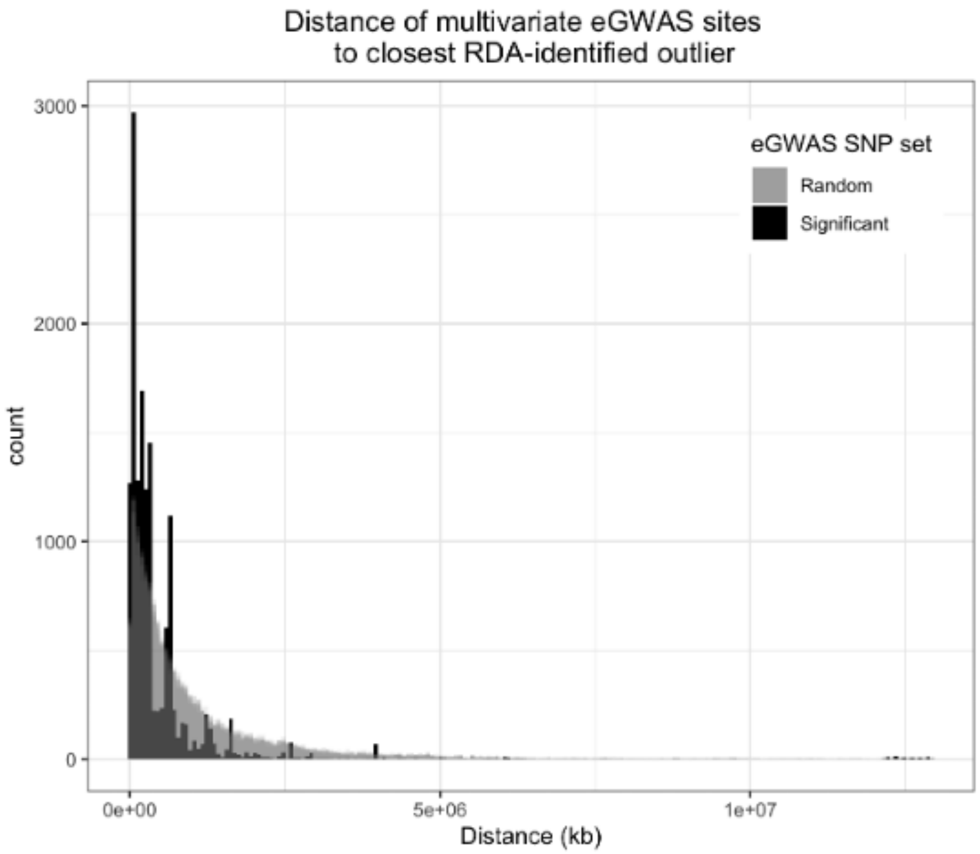

**Fig S10: Distance of RDA-identified variants to eGWAS associations.**
Histograms showing the distance of the RDA-identified variants to the nearest significant multivariate eGWAS associations (black) compared to the distance of the RDA-identified variants to randomly-selected SNPs (gray) from the multivariate eGWAS SNP set. Random sets are shown as the mean of 10 replicates with high transparency.

**Fig. S11**

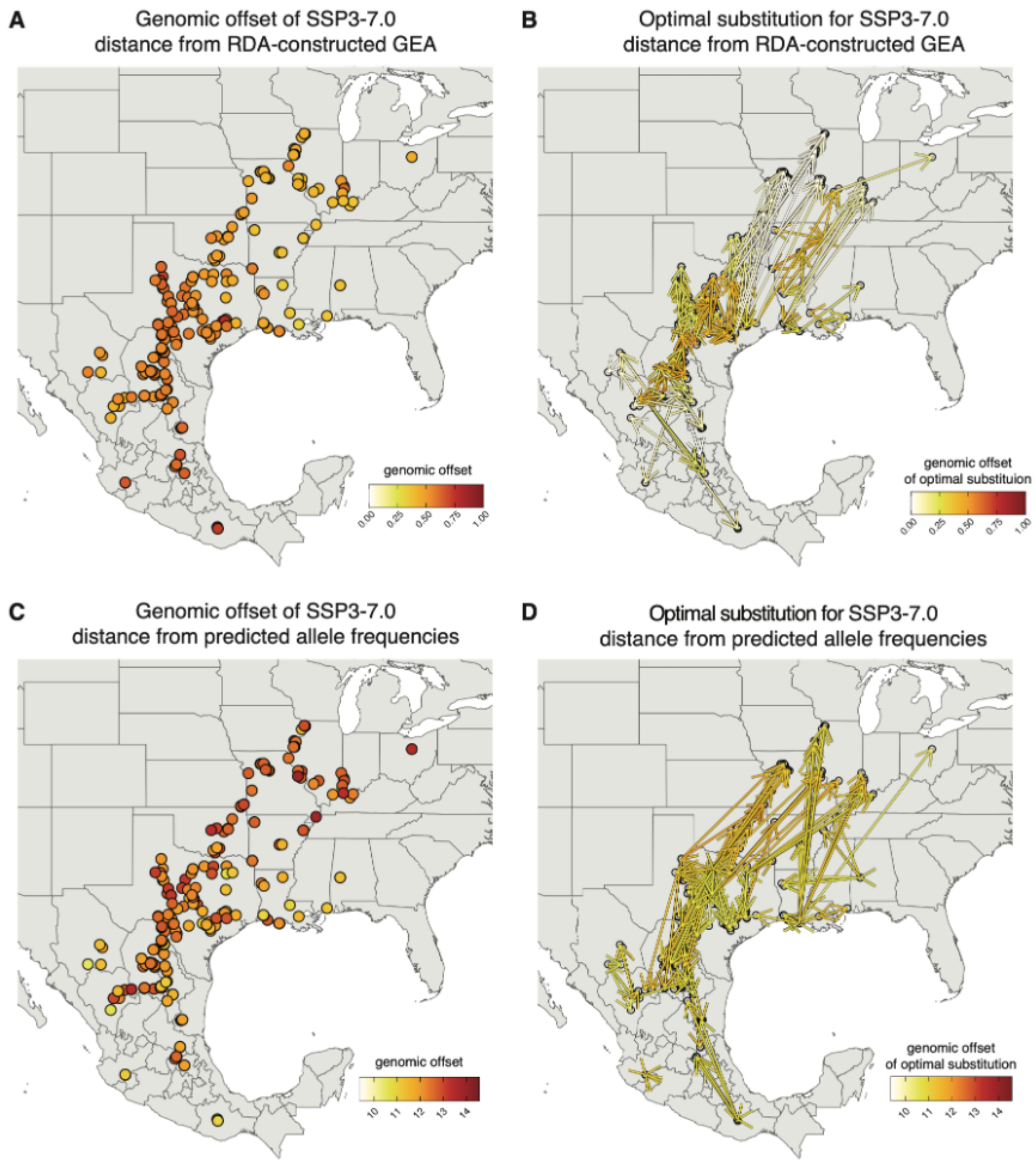

**Fig S11: Genomic offset and optimal substitutions across the native range for SSP3-7.0 scenario**

**A,B.** Genomic offset calculated from deviations in the loadings of RDA-constructed GEA space. **C,D.** Genomic offset calculated from differences in predicted and observed allele frequencies. **A,C:** Genomic offset for native pecan genotypes under the SSP3-7.0 emission scenario for their source location. **B,D:** Optimal substitution, defined as the pecan genotype that minimizes genomic offset to the novel environment under the SSP3-7.0 emission scenario. For each target location, arrows connect the optimal source genotype to the migration destination (arrowhead) and are colored by the remaining genomic offset of a substitution.

**Fig S12**

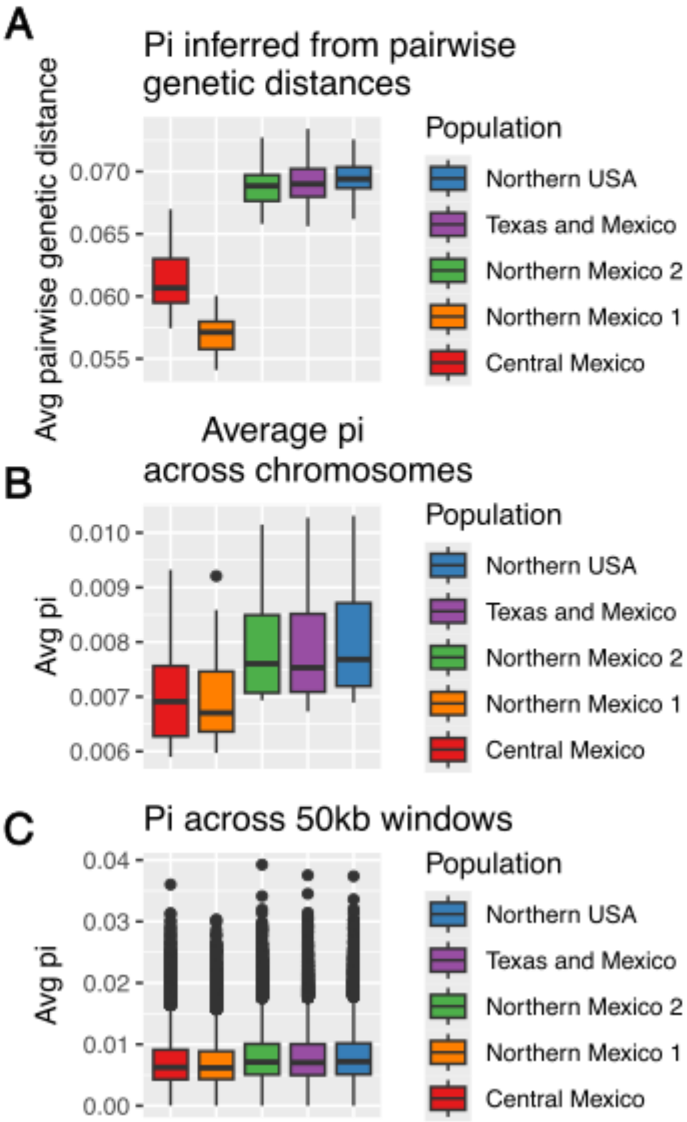

**Fig. S12: Comparison of methods for representing gene pool nucleotide diversity**
Comparison of different methods for representing nucleotide diversity (pi) in the pecan gene pools. **A.** Pairwise genetic distances for all samples assigned to each gene pool. Boxplots represent values from each pairwise comparison. **B.** Nucleotide diversity (pi) calculated in pixy (Korunes and Samuk 2021) using 50kb windows, averaged across each chromosome. Boxplots represent average pi of each chromosome. **C.** Nucleotide diversity (pi) calculated in pixy using 50kb windows. Boxplots represent average pi value in each of the 50kb windows across the genome.

### Supplemental Tables

**Table S1: 'Oaxaca' V2 Assembly Statistics and Comparisons**

| Assembly | Assembly Size (Mb) | Scaffold N50 (Mb) | Contig N50 (Mb) | N scaffolds | N contigs |
|---|---|---|---|---|---|
| Oaxaca V1 HAP1 | 670.0 | 44.3 | 14.0 | 16 | 100 |
| Oaxaca V1 HAP2 | 659.0 | 42.9 | 15.9 | 17 | 86 |
| Oaxaxa V1[1] | 650.0 | 42.3 | 4.4 | 298 | 552 |
| Pawnee V1[1] | 674.3 | 44.7 | 26.5 | 16 | 34 |
| Elliott V1[1] | 656.7 | 41.3 | 4.4 | 431 | 829 |
| Lakota V1[1] | 669.0 | 41.6 | 3.7 | 261 | 499 |

[1] (Lovell et al. 2021)

**Table S2: 'Oaxaca' V2 Assembly Sequence Content**

| Assembly | Genome Size (Mb) | Exons (Mb) | Introns (Mb) | Repeat Content (Mb) | Unannotated (Mb) |
|---|---|---|---|---|---|
| Oaxaca' V2 HAP1 | 670.0 | 39.37 | 74.31 | 311.63 | 244.65 |
| Oaxaca' V2 HAP2 | 659.0 | 39.18 | 75.41 | 304.44 | 239.81 |

**Table S3: Pecan samples genotyped using Illumina whole genome resequencing**

| Plant ID | Native Sample | ADMIXTURE Genepool[1] | Latitude | Longitude | VCF ID[2] | Illumina Sequencing Depth |
|---|---|---|---|---|---|---|
| 10-ILL-IA-4.1 | TRUE | NORTH_USA | 41.228 | -91.113 | IYQF | 57.3095 |
| 1950-12-0090 | FALSE | NA | NA | NA | 1950-12-0090 | 27.484 |
| 1964-06-0233 | FALSE | NA | NA | NA | 1964-06-0233 | 25.537 |
| 1972-02-0009 | FALSE | NA | 28.593 | -98.991 | 1972-02-0009 | 31.5326 |
| 1972-06-0012 | FALSE | NA | 28.593 | -98.991 | 1972-06-0012 | 27.8965 |
| 1975-08-0005 (Pueblo) | FALSE | NA | NA | NA | 1975-08-0005 | 29.0241 |
| 1980-08-0017 | FALSE | NA | NA | NA | IYQM | 78.2259 |
| 1996-01-0295 (Zuni) | FALSE | NA | NA | NA | 1996-01-0295 | 29.399 |
| 1996-02-0001 | FALSE | NA | NA | NA | 1996-02-0001-7 | 30.7877 |
| 1997-34-0017 | FALSE | NA | NA | NA | IYQL | 46.1453 |
| 2016-13-0025 | FALSE | NA | NA | NA | IYQG | 47.9131 |
| 86-IL-1-3.4 | TRUE | NORTH_USA | 39.086 | -90.616 | IXIJ | 42.4894 |

| | | | | | | |
|---|---|---|---|---|---|---|
| 86-KS-2-4.4 | TRUE | NORTH_USA | 37.062 | -95.060 | 86-KS-2-4-4 | 29.5402 |
| 86-KS-2-5.5 | TRUE | NORTH_USA | 37.025 | -95.046 | 86-KS-2-5-5 | 27.8546 |
| 86-MO-1-1.4 | TRUE | NORTH_USA | 39.758 | -93.608 | IXSE | 47.2712 |
| 86-MO-1-5.5 | TRUE | NORTH_USA | 39.707 | -93.305 | 86-MO-1-5-5 | 32.8824 |
| 86-TN-1-1.1 | TRUE | NORTH_USA | 36.377 | -89.501 | IXIK | 41.807 |
| 86-TX-1-3.1 | TRUE | TX_MX | 29.486 | -97.466 | IXDF | 53.4122 |
| 86-TX-1-4.5 | TRUE | TX_MX | 29.490 | -97.466 | IXIL | 37.8607 |
| 86-TX-1-5.4 | TRUE | TX_MX | 29.495 | -97.460 | IXDG | 44.4934 |
| 86-TX-2-1.5 | TRUE | TX_MX | 28.918 | -99.795 | IXSF | 44.3284 |
| 86-TX-2-2.5 | TRUE | TX_MX | 28.910 | -99.781 | IXSG | 45.6454 |
| 86-TX-3-1.2 | TRUE | ADMIX | 29.230 | -100.482 | IXSH | 57.629 |
| 86-TX-4-5.5 | TRUE | TX_MX | 30.025 | -101.169 | IXIM | 40.6233 |
| 86TX5-1.102 | TRUE | TX_MX | 31.352 | -100.207 | IXSK | 53.9544 |
| 86-TX-5-1.3 | TRUE | TX_MX | 31.352 | -100.207 | 86-TX-5-1-3 | 30.1723 |
| 86-TX-5-2.3 | TRUE | NORTH_MX_1 | 31.354 | -100.209 | IXSI | 48.6399 |
| 86-TX-5-3.3 | TRUE | TX_MX | 31.357 | -100.209 | IXSJ | 40.629 |
| 86-TX-5-4.1 | TRUE | TX_MX | 31.482 | -100.471 | IXDH | 48.4913 |
| 86-TX-5-5.5 | TRUE | TX_MX | 31.481 | -100.467 | IXIN | 49.7485 |
| 87-MX-1-1.2 | TRUE | TX_MX | 21.667 | -99.539 | IXDI | 49.6145 |
| 87-MX-1-1.6 | TRUE | TX_MX | 21.667 | -99.539 | IXSL | 43.5047 |
| 87-MX-1-2.2 | TRUE | CENTRAL_MX | 21.672 | -99.539 | IXIP | 45.8619 |
| 87-MX-1-2.4 | TRUE | CENTRAL_MX | 21.672 | -99.539 | IXDJ | 43.5538 |
| 87-MX-1-4.1 | TRUE | CENTRAL_MX | 21.678 | -99.543 | IXDK | 33.2626 |
| 87-MX-2-1.1 | TRUE | TX_MX | 19.891 | -103.586 | IXIQ | 44.6248 |
| 87-MX-2-2.1 | TRUE | CENTRAL_MX | 19.893 | -103.588 | IXSM | 51.5357 |
| 87-MX-2-3.1 | TRUE | CENTRAL_MX | 19.899 | -103.596 | IXIR | 43.1709 |
| 87-MX-2-4.3 | TRUE | CENTRAL_MX | 19.901 | -103.596 | IXIS | 33.6737 |
| 87-MX-3-1.5 | TRUE | NORTH_MX_1 | 17.047 | -96.790 | IXDL | 39.4073 |
| 87MX3-2.11 | TRUE | CENTRAL_MX | 16.951 | -96.756 | IIMT | 157.969 |
| 87-MX-3-2.2 | TRUE | CENTRAL_MX | 16.951 | -96.756 | 87-MX-3-2-2 | 26.3596 |
| 87-MX-3-2.4 | TRUE | CENTRAL_MX | 16.951 | -96.756 | IXDM | 47.9078 |

| | | | | | | |
|---|---|---|---|---|---|---|
| 87-MX-4-2.1 | TRUE | CENTRAL_MX | 20.484 | -99.222 | IXDN | 52.0544 |
| 87-MX-4-2.6 | TRUE | CENTRAL_MX | 20.484 | -99.222 | IXDP | 54.0525 |
| 87-MX-4-3.3 | TRUE | CENTRAL_MX | 20.485 | -99.220 | IXIT | 33.7886 |
| 87-MX-4-3.6 | TRUE | CENTRAL_MX | 20.485 | -99.220 | IXDQ | 33.77 |
| 87-MX-4-5.5 | TRUE | CENTRAL_MX | 20.486 | -99.221 | IXDR | 44.3729 |
| 87-MX-5-1.7 | TRUE | CENTRAL_MX | 23.405 | -99.381 | IXDS | 43.9833 |
| 87-MX-5-3.7 | TRUE | CENTRAL_MX | 23.404 | -99.382 | IXSN | 46.2141 |
| 87-MX-5-4.6 | TRUE | TX_MX | 23.369 | -99.479 | IXDT | 40.6341 |
| 87-MX-5-5.2 | TRUE | CENTRAL_MX | 23.367 | -99.475 | IXSP | 47.3862 |
| 87-MX-5-5.5 | TRUE | CENTRAL_MX | 23.367 | -99.475 | IXIU | 40.5012 |
| 89-ILL-ALA-3.1 | TRUE | NORTH_USA | 32.497 | -87.716 | 89-ILL-ALA-3-1 | 33.6637 |
| 89-ILL-ALA-3.4 | TRUE | NORTH_USA | 32.497 | -87.716 | IXDU | 47.0526 |
| 89-ILL-ALA-4.1 | TRUE | NORTH_USA | 32.497 | -87.716 | IXSQ | 49.1915 |
| 89-ILL-CHAMP-1.1 | TRUE | TX_MX | 32.309 | -100.800 | IXDW | 45.8898 |
| 89-ILL-CHAMP-1.8 | TRUE | TX_MX | 33.049 | -100.823 | 89-ILL-SCHAMP-1-1 | 27.7841 |
| 89-ILL-CHAMP-2.2 | TRUE | TX_MX | 32.309 | -100.801 | IXDX | 56.9192 |
| 89-ILL-CHAMP-4.1 | TRUE | TX_MX | 32.320 | -100.746 | IXDY | 43.789 |
| 89-ILL-CONCH-2.1 | TRUE | TX_MX | 31.790 | -100.416 | 89-ILL-CONCH-2-1 | 29.7659 |
| 89-ILL-CONCH-2.3 | TRUE | TX_MX | 31.481 | -100.467 | IXDZ | 49.0895 |
| 89-ILL-CONCH-4.1 | TRUE | TX_MX | 31.482 | -100.471 | IXEA | 46.1566 |
| 89-ILL-CONCH-4.5 | TRUE | TX_MX | 31.790 | -100.416 | 89-ILL-CONCH-4 | 24.8908 |
| 89-ILL-CONCH-4.8 | TRUE | TX_MX | 31.790 | -100.416 | 89-ILL-CONCHO-4 | 32.7673 |
| 89-ILL-DEV-1.12 | TRUE | NORTH_MX_2 | 29.967 | -101.149 | IXEB | 56.7958 |
| 89-ILL-DEV-1.9 | TRUE | NORTH_MX_2 | 29.967 | -101.149 | IXIW | 41.4233 |
| 89-ILL-DEV-3.1 | TRUE | NORTH_MX_2 | 29.982 | -101.168 | IXEC | 59.8132 |
| 89-ILL-ED-1.6 | TRUE | NORTH_MX_1 | 29.842 | -100.160 | IXED | 51.0279 |
| 89-ILL-ED-6.3 | TRUE | TX_MX | 29.792 | -100.415 | IXIX | 45.3936 |
| 89-ILL-HAYES-1.1 | TRUE | TX_MX | 32.797 | -101.010 | IXIY | 43.8488 |
| 89-ILL-HAYES-3.2 | TRUE | TX_MX | 32.796 | -101.006 | IXIZ | 56.3203 |

| | | | | | | |
|---|---|---|---|---|---|---|
| 89-ILL-IL-4.1 | TRUE | NORTH_USA | 39.301 | -90.595 | IXJA | 40.7301 |
| 89-ILL-ILL-6.3 | TRUE | TX_MX | 41.978 | -90.552 | IXJC | 45.9845 |
| 89-ILL-LA-1.6 | TRUE | TX_MX | 29.784 | -93.087 | IXJD | 50.0262 |
| 89-ILL-LA-3.4 | TRUE | TX_MX | 29.785 | -93.086 | IXEE | 47.9059 |
| 89-ILL-LIPAN-3.2 | TRUE | TX_MX | 31.357 | -100.209 | IXJE | 44.8527 |
| 89-ILL-LIPAN-3.3 | TRUE | TX_MX | 31.357 | -100.209 | IXEF | 48.9478 |
| 89-ILL-LMO-2.4 | TRUE | NORTH_MX_2 | 29.290 | -100.425 | IXSS | 35.6102 |
| 89-ILL-LMO-2.5 | TRUE | TX_MX | 30.130 | -100.422 | 89-ILL-CONCHO-2 | 25.5284 |
| 89-ILL-LMO-4.6 | TRUE | TX_MX | 29.290 | -100.424 | IXJF | 51.8601 |
| 89-ILL-OK-1.7 | TRUE | NORTH_USA | 34.090 | -96.960 | IXJG | 48.0391 |
| 89-ILL-OK-5.8 | TRUE | NORTH_USA | 34.361 | -96.635 | IXEG | 48.2632 |
| 89-ILL-SBRA-2.4 | TRUE | TX_MX | 33.214 | -96.109 | 89-ILL-SBRA-2-4 | 28.7702 |
| 89-ILL-SBRA-2.8 | TRUE | TX_MX | 33.214 | -96.109 | IXEH | 34.0565 |
| 89-ILL-SBRA-4.4 | TRUE | TX_MX | 32.746 | -96.101 | IXEI | 62.4147 |
| 89-ILL-SBRA-6.1 | FALSE | NA | NA | NA | IXEJ | 51.1223 |
| 89-ILL-SFE-1.6 | TRUE | NORTH_USA | 29.371 | -100.884 | 89-ILL-SFE-1-6 | 29.8264 |
| 89-ILL-SFE-3.5 | TRUE | ADMIX | 29.361 | -100.890 | IXEK | 35.4969 |
| 89-ILL-STATON-1.1 | TRUE | TX_MX | 32.571 | -100.905 | IXJH | 38.0485 |
| 89-ILL-STATON-2.5 | TRUE | TX_MX | 32.573 | -100.907 | IXJI | 39.7376 |
| 89-ILL-UV-3.1 | TRUE | TX_MX | 29.531 | -100.006 | IXEL | 70.0077 |
| 89-ILL-UV-4.1 | TRUE | TX_MX | 29.530 | -100.005 | IYPI | 53.2192 |
| 94-ILL-ARK-1.3 | TRUE | NORTH_USA | 34.610 | -92.185 | IXEN | 43.0999 |
| 94-ILL-ARK-2.4 | TRUE | NORTH_USA | 34.604 | -92.189 | IXEP | 49.8704 |
| 94-ILL-ARK-3.9 | TRUE | NORTH_USA | 34.599 | -92.192 | IXJJ | 33.8267 |
| 94-ILL-ARK-4.6 | TRUE | NORTH_USA | 34.598 | -92.192 | IXJK | 39.3568 |
| 94-ILL-ARK-5.1 | TRUE | NORTH_USA | 34.593 | -92.195 | IXJL | 44.3379 |
| 94-ILL-ELK-2.2 | TRUE | NORTH_USA | 36.040 | -94.052 | IXEQ | 52.0228 |
| 94-ILL-ELK-3.1 | TRUE | NORTH_USA | 36.035 | -94.039 | IXER | 40.1459 |
| 94-ILL-RR-1.8 | TRUE | NORTH_USA | 33.573 | -94.203 | IXES | 42.06 |
| 94-ILL-RR-2.2 | TRUE | NORTH_USA | 33.571 | -94.207 | IYPJ | 50.4862 |

| | | | | | | |
|---|---|---|---|---|---|---|
| 94-ILL-RR-3.7 | TRUE | NORTH_USA | 33.569 | -94.210 | IXEU | 39.9736 |
| 94-ILL-RR-4.6 | TRUE | NORTH_USA | 33.567 | -94.207 | IXEW | 40.9658 |
| 94-ILL-RR-5.6 | TRUE | NORTH_USA | 33.566 | -94.209 | IXEX | 33.2471 |
| 94-ILL-RR-6.7 | TRUE | NORTH_USA | 33.569 | -94.204 | IXEY | 55.7702 |
| 94-ILL-STF-1.6 | TRUE | NORTH_USA | 35.522 | -90.431 | IXEZ | 47.851 |
| 94-ILL-STF-2.5 | TRUE | NORTH_USA | 35.522 | -90.432 | IXFA | 47.288 |
| 94-ILL-STF-3.8 | TRUE | NORTH_USA | 35.511 | -90.446 | IXJM | 38.9798 |
| 94-ILL-STF-4.5 | TRUE | NORTH_USA | 35.511 | -90.446 | IXFB | 47.678 |
| 94-ILL-STF-5.4 | TRUE | NORTH_USA | 35.511 | -90.444 | IXJN | 59.6897 |
| 97-ILL-CAT-1.1 | TRUE | NORTH_USA | 29.571 | -92.239 | IXST | 44.2549 |
| 97-ILL-CAT-11.3 | TRUE | NORTH_USA | 29.569 | -92.204 | IYPL | 45.1805 |
| 97-ILL-CAT-12.2 | TRUE | NORTH_USA | 29.569 | -92.204 | IXJQ | 43.5113 |
| 97-ILL-CAT-13.9 | TRUE | NORTH_USA | 29.569 | -92.203 | IXJR | 47.0283 |
| 97-ILL-CAT-14.13 | TRUE | NORTH_USA | 29.569 | -92.203 | IXJS | 44.6441 |
| 97-ILL-CAT-15.15 | TRUE | NORTH_USA | 29.569 | -92.203 | IXJT | 33.841 |
| 97-ILL-CAT-16.2 | TRUE | NORTH_USA | 29.569 | -92.203 | IXJU | 40.8507 |
| 97-ILL-CAT-7.17 | TRUE | NORTH_USA | 29.570 | -92.224 | IYPK | 49.759 |
| 97-ILL-CAT-8.5 | TRUE | NORTH_USA | 29.571 | -92.223 | IXJX | 41.3798 |
| 97-ILL-CAT-9.4 | TRUE | NORTH_USA | 29.570 | -92.222 | IXJY | 46.1807 |
| 97-ILL-LA-1.12 | TRUE | NORTH_USA | 30.065 | -93.351 | IXJZ | 35.6111 |
| 97-ILL-LA-WB1.14 | TRUE | NORTH_USA | 30.094 | -93.391 | IXKB | 37.5828 |
| 97-ILL-LA-WB2.3 | TRUE | NORTH_USA | 30.095 | -93.392 | IXKC | 35.9231 |
| 97-ILL-LA-WB3.1 | TRUE | NORTH_USA | 30.095 | -93.392 | IXSU | 41.0516 |
| 99-ILL-SC-2 | FALSE | NA | 33.033 | -80.371 | IXYI | 51.7405 |
| A-93 | FALSE | NA | NA | NA | IXSW | 45.2708 |
| Allbritton | TRUE | NORTH_USA | 32.120 | -93.520 | IXYJ | 38.2884 |
| Alley | FALSE | NA | 30.433 | -88.533 | IXSX | 45.9175 |
| Altman | TRUE | TX_MX | 30.769 | -98.264 | IYPQ | 65.8999 |
| Ames 25281.1 | FALSE | NA | NA | NA | Ames-25281-1 | 27.2641 |
| Amling | FALSE | NA | NA | NA | IXSY | 51.9473 |
| Anson Jones | TRUE | TX_MX | 30.322 | -96.147 | IXFC | 40.7148 |

| | | | | | | |
|---|---|---|---|---|---|---|
| Apache | FALSE | NA | NA | NA | IXKE | 35.3262 |
| Apalachee | FALSE | NA | NA | NA | Apalachee | 29.9381 |
| Aquilla | FALSE | NA | 31.832 | -97.194 | IXSZ | 53.1246 |
| Baker | FALSE | NA | 32.599 | -93.290 | IXTA | 45.6669 |
| Barraza 3 | FALSE | NA | 32.277 | -91.714 | Barraza-3 | 28.6386 |
| Barton | FALSE | NA | NA | NA | IXTB | 67.804 |
| Bean | FALSE | NA | NA | NA | Bean | 28.6698 |
| Bennett | FALSE | NA | 28.594 | -98.991 | Bennett | 30.3678 |
| Best's Early | TRUE | NORTH_USA | 39.288 | -90.552 | IXYK | 51.184 |
| Big Boy | FALSE | NA | 29.389 | -98.459 | IXTC | 47.3861 |
| Big Sandy | FALSE | NA | NA | NA | Big-Sandy | 25.924 |
| Biggs | FALSE | NA | 32.442 | -97.794 | Biggs | 27.2218 |
| Bolds | TRUE | NORTH_USA | 37.601 | -86.990 | Bolds | 28.5083 |
| Bolton | FALSE | NA | NA | NA | Bolton | 26.0761 |
| Bradley (FL) | FALSE | NA | NA | NA | IYPU | 54.9281 |
| Brake | FALSE | NA | 35.780 | -78.638 | Brake | 29.2213 |
| Branch | FALSE | NA | 30.400 | -88.583 | Branch | 27.6344 |
| Bridges | TRUE | TX_MX | 33.213 | -97.761 | Bridges | 27.8518 |
| Brooks | FALSE | NA | 33.291 | -84.459 | Brooks | 27.9793 |
| Brown's Leaning Tree | TRUE | TX_MX | 29.531 | -100.006 | IXYL | 46.9862 |
| Bryce | TRUE | NORTH_USA | 39.756 | -93.560 | IXYM | 36.7019 |
| Brzozowski-Vlasek | TRUE | TX_MX | 29.523 | -97.491 | IXYN | 47.8761 |
| Buchel 1 | TRUE | TX_MX | 29.098 | -97.285 | IXTD | 55.9061 |
| Burkett | TRUE | TX_MX | 32.374 | -99.163 | IXTE | 52.8258 |
| Busseron | TRUE | NORTH_USA | 38.835 | -87.449 | IXYP | 43.3991 |
| Byrd | FALSE | NA | NA | NA | Byrd-58 | 31.4009 |
| C. Boles | FALSE | NA | NA | NA | C-Boles | 29.2223 |
| Caddo | FALSE | NA | NA | NA | IWRX | 40.4061 |
| Candy | FALSE | NA | 30.429 | -88.766 | IXTF | 55.8007 |
| Canton | TRUE | NORTH_USA | 40.132 | -91.530 | IXYQ | 41.0657 |
| CapeFear | FALSE | NA | 34.690 | -77.979 | IXYR | 37.8944 |

| | | | | | | |
|---|---|---|---|---|---|---|
| Carden | TRUE | TX_MX | 31.713 | -96.163 | IXYS | 42.1617 |
| Carlson #3 | TRUE | NORTH_USA | 41.176 | -91.000 | Carlson-3 | 26.4112 |
| Carman | FALSE | NA | 32.339 | -91.024 | IXYT | 53.8879 |
| Carmichael | TRUE | TX_MX | 29.655 | -97.368 | IXTG | 56.8363 |
| Carole Leigh | FALSE | NA | 30.431 | -87.745 | Carole-Leigh | 26.2255 |
| Carter | FALSE | NA | 30.411 | -88.828 | IXKF | 39.5415 |
| Centennial | TRUE | NORTH_USA | 30.004 | -90.776 | IXKG | 33.7747 |
| Champ 15 | FALSE | NA | 30.283 | -96.096 | Champ-15 | 26.8306 |
| Cherokee | FALSE | NA | NA | NA | Cherokee | 26.4458 |
| Cherryle | FALSE | NA | 30.476 | -88.342 | IXTH | 52.7595 |
| Chetopa (K-112) | TRUE | NORTH_USA | 37.023 | -95.046 | Chetopa-K-112- | 26.4003 |
| Chickasaw | FALSE | NA | NA | NA | Chickasaw | 29.0831 |
| Chief | TRUE | NORTH_USA | 37.806 | -88.262 | IXYU | 53.6946 |
| Choctaw | FALSE | NA | NA | NA | IWRZ | 53.1788 |
| Clark | TRUE | TX_MX | 31.105 | -98.505 | IXTI | 43.9477 |
| Colby | TRUE | NORTH_USA | 38.679 | -89.309 | IXTJ | 34.9009 |
| Comanche | FALSE | NA | NA | NA | IWSA | 42.0327 |
| Cooper | FALSE | NA | 32.015 | -93.342 | IXTK | 55.6739 |
| Cordele Dwarf | FALSE | NA | 31.963 | -83.783 | Cordele-Dwarf | 25.5375 |
| Cordele Dwarf Sdlg 1 | FALSE | NA | NA | NA | Cordele-Dwarf-Sdlg-1 | 29.5401 |
| Cordele Dwarf Sdlg 2 | FALSE | NA | NA | NA | Cordele-Dwarf-Sdlg-2 | 28.9044 |
| Creek | FALSE | NA | NA | NA | Creek | 30.1535 |
| Curtis | FALSE | NA | 30.600 | -87.012 | IXYW | 50.3746 |
| Davis | TRUE | NORTH_USA | 30.540 | -88.688 | Davis | 26.6131 |
| Deakle's Special | FALSE | NA | 30.507 | -88.234 | Deakle-s-Special | 28.2931 |
| Delmas | FALSE | NA | 30.368 | -88.554 | IXKH | 34.4801 |
| Dependable | FALSE | NA | 30.411 | -88.828 | IXTM | 33.1172 |
| Desirable | FALSE | NA | 30.411 | -88.828 | IXTN | 61.5741 |
| Die Guet | TRUE | TX_MX | 29.411 | -98.905 | IXTP | 36.7664 |

| | | | | | | |
|---|---|---|---|---|---|---|
| Dixie | FALSE | NA | 30.790 | -87.780 | Dixie | 29.0648 |
| Dooley | TRUE | NORTH_USA | 35.622 | -96.014 | Dooley | 26.9172 |
| Dowds | FALSE | NA | 39.065 | -88.748 | Dowds | 28.426 |
| Duck Creek 1 | TRUE | TX_MX | 33.684 | -100.939 | IXYX | 35.6414 |
| Duck Creek 16 | TRUE | TX_MX | 33.684 | -100.939 | IXKI | 35.1568 |
| Duck Creek 25 | TRUE | TX_MX | 33.683 | -100.939 | Duck-Creek-25 | 30.153 |
| Dumbell Lake Small | TRUE | NORTH_USA | 40.863 | -91.072 | IXYZ | 37.939 |
| E Z Peel | FALSE | NA | 30.283 | -96.096 | E-Z-Peel | 26.7136 |
| El Mart | FALSE | NA | 31.630 | -99.510 | El-Mart | 28.6369 |
| Elgin | FALSE | NA | 29.955 | -97.370 | IXTQ | 37.5026 |
| Elliott | FALSE | NA | 30.631 | -87.056 | IPZM | 146.136 |
| Esnuel | FALSE | NA | 31.000 | -87.500 | Esnuel | 26.8963 |
| Evans | TRUE | TX_MX | 29.867 | -96.817 | IXTR | 49.3374 |
| Evers | TRUE | NORTH_MX_2 | 33.206 | -97.080 | IXTS | 46.5189 |
| Excel | FALSE | NA | 31.302 | -82.247 | IXZB | 55.9331 |
| F.W. Anderson | FALSE | NA | 37.229 | -120.248 | F-W-Anderson | 26.2292 |
| Family Use | TRUE | TX_MX | 31.196 | -98.718 | IXTT | 36.3442 |
| Fangue | TRUE | TX_MX | 29.945 | -96.120 | IYPG | 60.3165 |
| Farley | FALSE | NA | 30.800 | -85.020 | IXTU | 59.4038 |
| Fat Cat | FALSE | NA | NA | NA | Fat-Cat | 27.6776 |
| Fayette | FALSE | NA | 29.899 | -96.872 | Fayette | 31.8658 |
| Fifth Row | FALSE | NA | NA | NA | Fifth-Row | 26.9728 |
| Fisher | TRUE | NORTH_USA | 38.479 | -89.678 | Fisher | 29.311 |
| Flack | FALSE | NA | 37.320 | -87.511 | Flack | 25.363 |
| Forey | FALSE | NA | 30.125 | -91.833 | Forey | 25.9568 |
| Forkert | FALSE | NA | 30.411 | -88.828 | IXTW | 37.8907 |
| Foster | TRUE | TX_MX | 29.461 | -97.658 | IXTX | 55.1448 |
| Freeman | TRUE | TX_MX | 32.816 | -95.664 | IXTY | 41.5168 |
| Frisbie | TRUE | NORTH_USA | 39.754 | -93.550 | IXTZ | 39.3165 |
| Fritz (TX) | FALSE | NA | 30.283 | -96.096 | IXZD | 73.2257 |
| Frotscher | FALSE | NA | 29.975 | -91.756 | Frotscher | 27.4333 |

| | | | | | | |
|---|---|---|---|---|---|---|
| Frutoso | TRUE | NORTH_MX_2 | 25.420 | -102.172 | IXZE | 36.7354 |
| Gafford | FALSE | NA | 31.788 | -86.453 | IXZF | 66.4517 |
| Gay | FALSE | NA | 29.465 | -96.539 | Gay | 27.526 |
| Gibson | FALSE | NA | 40.025 | -90.997 | Gibson | 26.6007 |
| Giles | TRUE | NORTH_USA | 37.062 | -95.060 | IXZG | 49.2903 |
| Gloria Grande | FALSE | NA | 33.493 | -80.851 | Gloria-Grande | 28.4857 |
| Goose Pond | TRUE | NORTH_USA | 39.326 | -92.959 | IXZH | 63.2582 |
| Gormely | TRUE | NORTH_USA | 35.433 | -96.305 | IXUA | 39.8652 |
| Govett | TRUE | TX_MX | 29.568 | -97.885 | IXUB | 45.7763 |
| GraCross | FALSE | NA | 30.335 | -96.111 | GraCross | 28.0948 |
| Graf | FALSE | NA | NA | NA | Graf | 29.6482 |
| GraKing | FALSE | NA | 34.011 | -95.509 | IXZI | 44.8227 |
| Grandma's Native | FALSE | NA | NA | NA | Grandma-s-Native | 28.3441 |
| GraPark Giant | FALSE | NA | 32.746 | -96.998 | IXUC | 38.4359 |
| GraTex | FALSE | NA | 26.645 | -97.452 | IXZJ | 57.3977 |
| GraZona | FALSE | NA | 33.422 | -111.822 | IXZK | 44.4648 |
| GRE 14-20 | FALSE | NA | NA | NA | GRE-14-20 | 33.4231 |
| Green Island Beaver | TRUE | NORTH_USA | 42.177 | -90.301 | IXZL | 54.4353 |
| Green Island Hackberry | TRUE | NORTH_USA | 42.182 | -90.299 | IXZM | 41.3682 |
| Greenriver | TRUE | NORTH_USA | 37.898 | -87.491 | IYPS | 66.0588 |
| Guenther | FALSE | NA | 29.703 | -98.124 | IXZP | 48.2802 |
| Guidry | FALSE | NA | NA | NA | Guidry- | 27.8226 |
| Halbert | TRUE | TX_MX | 31.824 | -99.391 | IXZQ | 50.4547 |
| Halsly | FALSE | NA | 31.406 | -99.230 | Halsly | 26.1181 |
| Hark | FALSE | NA | 40.943 | -90.505 | IXZR | 35.9937 |
| Harper | FALSE | NA | NA | NA | IYPW | 64.009 |
| Harris Super | FALSE | NA | 33.944 | -90.945 | IXZS | 68.9237 |
| Hayes | TRUE | NORTH_USA | 35.645 | -96.833 | Hayes | 28.094 |
| Hensel | TRUE | TX_MX | 29.893 | -96.871 | IXUD | 33.5778 |
| Hirschi | TRUE | NORTH_USA | 38.064 | -94.238 | IXZT | 44.5833 |

| | | | | | | |
|---|---|---|---|---|---|---|
| Hodge | TRUE | NORTH_USA | 39.174 | -87.635 | IXZU | 42.1361 |
| Hollis | TRUE | TX_MX | 31.100 | -98.515 | IXUE | 37.7649 |
| Hopi | FALSE | NA | NA | NA | Hopi | 29.3881 |
| Houma | FALSE | NA | NA | NA | Houma | 26.0314 |
| Hughes | FALSE | NA | 30.411 | -88.828 | IXZW | 45.0206 |
| Humble | TRUE | TX_MX | 29.087 | -99.875 | IXUF | 55.4833 |
| Iago | FALSE | NA | 31.630 | -99.510 | IXZX | 53.3296 |
| Ideal_(Bradley) | TRUE | TX_MX | 31.196 | -98.718 | IXUG | 36.7636 |
| Jack Doby | FALSE | NA | 30.009 | -97.159 | Jack-Doby | 30.2539 |
| Jackson | FALSE | NA | 30.411 | -88.828 | Jackson | 28.0144 |
| James(LA) | FALSE | NA | 32.340 | -91.024 | IXUH | 45.5013 |
| Jefferson GTV | FALSE | NA | NA | NA | Jefferson-GTV | 28.1782 |
| Jenkins | FALSE | NA | 34.151 | -90.782 | Jenkins | 27.6983 |
| Jersey | TRUE | TX_MX | 31.250 | -98.596 | IXUI | 42.4107 |
| Jin Hua 1 | FALSE | NA | NA | NA | Jin-Hua-1 | 30.6471 |
| Johnson(KS) | TRUE | NORTH_USA | 37.176 | -94.820 | IXUJ | 40.4965 |
| Jubilee | FALSE | NA | 30.417 | -87.667 | IXUK | 49.2552 |
| Jumbo Joy | FALSE | NA | 33.767 | -86.587 | Jumbo-Joy | 27.5914 |
| Keilers | FALSE | NA | 30.244 | -97.744 | Keilers | 29.0145 |
| Kelly | TRUE | TX_MX | 31.196 | -98.718 | Kelly | 28.9203 |
| Kennedy | FALSE | NA | 29.717 | -82.140 | IXZY | 39.395 |
| Kentucky | TRUE | NORTH_USA | 37.835 | -86.762 | Kentucky | 31.7014 |
| Kernodle | FALSE | NA | 32.800 | -85.654 | IXZZ | 53.3125 |
| Kibler | FALSE | NA | 42.186 | -86.308 | IYPT | 46.5346 |
| Kincaid | TRUE | TX_MX | 31.148 | -98.806 | IYAB | 45.0677 |
| Kiowa | FALSE | NA | NA | NA | Kiowa | 28.6823 |
| Koko | FALSE | NA | 30.125 | -91.833 | Koko | 25.8514 |
| KS 2-1 | FALSE | NA | NA | NA | KS-2-1 | 27.8397 |
| Kuykendall | TRUE | TX_MX | 32.763 | -98.718 | Kuykendall | 27.4684 |
| La Bahia | TRUE | TX_MX | 30.329 | -96.155 | IXFD | 62.374 |
| Lacina | TRUE | TX_MX | 30.203 | -96.398 | IXUL | 47.7568 |

| | | | | | | |
|---|---|---|---|---|---|---|
| Lane's Grandma | TRUE | TX_MX | 30.236 | -96.240 | IYPH | 47.2474 |
| Late | TRUE | TX_MX | 31.374 | -98.677 | IXUM | 40.9397 |
| Lineberger | FALSE | NA | 31.550 | -83.547 | Lineberger | 30.0232 |
| Lipan | FALSE | NA | NA | NA | Lipan | 28.1145 |
| Little Jewel | TRUE | TX_MX | 29.530 | -100.005 | IXFE | 41.9498 |
| Longfellow | TRUE | NORTH_MX_1 | 31.193 | -98.847 | Longfellow | 26.6666 |
| Lucas | TRUE | NORTH_USA | 40.705 | -82.418 | IXUN | 33.9526 |
| Magenta | TRUE | NORTH_USA | 32.519 | -91.966 | IYAD | 43.1085 |
| Mahan | FALSE | NA | 33.058 | -89.588 | IXWF | 33.1166 |
| Mahan Stuart | FALSE | NA | NA | NA | Mahan-Stuart | 29.7274 |
| Mandan | FALSE | NA | NA | NA | Mandan | 28.6512 |
| Maramec | FALSE | NA | 36.242 | -96.680 | Maramec | 29.219 |
| Martzahn | TRUE | NORTH_USA | 40.838 | -91.115 | IYAE | 36.5334 |
| Mason Deer | TRUE | TX_MX | 30.656 | -99.279 | Mason-Deer | 28.0897 |
| McCulley | TRUE | TX_MX | 31.723 | -98.958 | IYAF | 55.426 |
| McMillan | FALSE | NA | 30.790 | -87.780 | IYAG | 41.9599 |
| Meier | FALSE | NA | NA | NA | Meier | 30.2726 |
| Melrose | TRUE | NORTH_USA | 31.961 | -93.348 | Melrose | 25.3129 |
| Menchaca | FALSE | NA | NA | NA | Menchaca | 27.9897 |
| Meyers(TX) | FALSE | NA | 30.209 | -97.592 | IXWG | 48.18 |
| Miss L | FALSE | NA | NA | NA | Miss-L | 27.1749 |
| Mississippi 10 | FALSE | NA | 32.730 | -89.700 | Mississippi-10 | 27.8201 |
| MO-AES-2 | FALSE | NA | 38.959 | -92.343 | MO-AES-2 | 31.8882 |
| Mobile | FALSE | NA | 30.407 | -88.261 | Mobile | 26.7566 |
| Mohawk | FALSE | NA | NA | NA | IWWM | 51.8262 |
| Moneymaker | FALSE | NA | 32.339 | -91.024 | IXWH | 37.4135 |
| Montgomery | FALSE | NA | 32.516 | -93.732 | Montgomery | 27.6622 |
| Moore | FALSE | NA | 30.411 | -83.953 | IXKW | 40.4394 |
| Moreland | FALSE | NA | 31.873 | -93.201 | Moreland | 27.8472 |
| Mount | TRUE | NORTH_USA | 35.584 | -95.992 | IXWI | 47.9412 |
| Mullahy | TRUE | NORTH_USA | 41.172 | -90.996 | IXWJ | 51.4785 |

| | | | | | | |
|---|---|---|---|---|---|---|
| MX94-001-8 | TRUE | NORTH_MX_2 | 28.492 | -100.920 | IXFF | 37.5187 |
| MX94-002-9 | TRUE | NORTH_MX_2 | 27.929 | -101.304 | IWSB | 37.3389 |
| MX94-003-4 | TRUE | NORTH_MX_2 | 29.069 | -100.679 | IWSC | 48.1636 |
| MX94-004-5 | TRUE | NORTH_MX_2 | 27.604 | -100.728 | IWSD | 46.4795 |
| MX94-006-4 | TRUE | NORTH_MX_2 | 27.883 | -101.517 | MX94-006-4 | 27.0512 |
| MX94-007-6 | TRUE | NORTH_MX_2 | 26.826 | -102.255 | IXFG | 37.4068 |
| MX94-008-2 | TRUE | NORTH_MX_2 | 27.604 | -100.728 | IWSE | 39.3417 |
| MX94-009-2 | TRUE | NORTH_MX_2 | 26.785 | -101.430 | IXKM | 42.696 |
| MX94-010-8 | TRUE | NORTH_MX_2 | 26.826 | -102.255 | IWSF | 39.4291 |
| MX94-011-4 | TRUE | ADMIX | 25.350 | -103.133 | MX94-011-4 | 29.2064 |
| MX94-012-3 | TRUE | NORTH_MX_2 | 26.950 | -102.083 | 94-MX-012-3 | 28.2736 |
| MX94-012-9 | TRUE | NORTH_MX_2 | 26.826 | -102.255 | IXFH | 37.5821 |
| MX94-013-6 | TRUE | NORTH_MX_2 | 28.492 | -100.920 | IXKN | 35.1377 |
| MX94-014-5 | TRUE | ADMIX | 25.350 | -103.133 | IWUQ | 36.5852 |
| MX94-014-7 | TRUE | ADMIX | 28.492 | -100.920 | IXFI | 33.7057 |
| MX94-015-5 | TRUE | NORTH_MX_1 | 28.492 | -100.920 | IWSG | 40.2438 |
| MX94-016-4 | TRUE | NORTH_MX_2 | 25.350 | -103.133 | MX94-016-4 | 29.1066 |
| MX94-017-4 | TRUE | TX_MX | 26.900 | -101.417 | MX94-017-4 | 31.1581 |
| MX94-018-5 | TRUE | NORTH_MX_2 | 26.826 | -102.255 | IXUP | 56.8087 |
| MX94-019-3 | TRUE | NORTH_MX_1 | 27.033 | -101.800 | MX94-019-3 | 28.8009 |
| MX94-020-4 | TRUE | NORTH_MX_2 | 27.004 | -101.724 | IXUQ | 35.2941 |
| MX94-021-5 | TRUE | NORTH_MX_2 | 26.783 | -101.433 | MX94-021-5 | 28.4626 |
| MX94-022-4 | TRUE | NORTH_MX_2 | 26.783 | -101.433 | MX94-022-4 | 29.6397 |
| MX94-023-7 | TRUE | NORTH_MX_2 | 26.785 | -101.430 | IWSH | 54.3452 |
| MX94-024-5 | TRUE | NORTH_MX_2 | 26.833 | -100.667 | MX94-024-5 | 29.1805 |
| MX94-025-3 | TRUE | NORTH_MX_1 | 26.900 | -101.417 | MX94-025-3 | 28.3637 |
| MX94-026-5 | TRUE | NORTH_MX_2 | 27.311 | -102.397 | IWSI | 58.2497 |
| MX94-027-9 | TRUE | NORTH_MX_2 | 26.785 | -101.430 | IWSJ | 53.5471 |
| MX94-028-6 | TRUE | NORTH_MX_2 | 26.783 | -101.433 | MX94-028-6 | 30.38 |
| MX94-029-3 | TRUE | TX_MX | 25.350 | -103.133 | MX94-029-3 | 28.5356 |
| MX94-030-6 | TRUE | NORTH_MX_2 | 26.826 | -102.255 | IWSK | 38.5061 |

| | | | | | | |
|---|---|---|---|---|---|---|
| MX94-031-1 | TRUE | NORTH_MX_2 | 26.950 | -102.083 | MX94-031-1 | 29.3785 |
| MX94-032-6 | TRUE | TX_MX | 26.950 | -102.083 | MX94-032-6 | 27.9217 |
| MX94-033-4 | TRUE | NORTH_MX_2 | 26.950 | -102.083 | MX94-033-4 | 31.1672 |
| MX94-033-7 | TRUE | NORTH_MX_2 | 26.950 | -102.083 | 94-MX-033-7 | 25.4414 |
| MX94-034-3 | TRUE | NORTH_MX_2 | 26.950 | -102.083 | MX94-034-3 | 27.7127 |
| MX94-035-3 | TRUE | ADMIX | 26.950 | -102.083 | MX94-035-3 | 29.8186 |
| MX94-036-4 | TRUE | NORTH_MX_2 | 26.826 | -102.255 | IWSL | 39.1105 |
| MX94-036-6 | TRUE | NORTH_MX_2 | 26.826 | -102.255 | IWSM | 42.1047 |
| MX94-037-3 | TRUE | NORTH_MX_2 | 26.826 | -102.255 | IWSN | 45.9774 |
| MX94-038-6 | TRUE | NORTH_MX_2 | 26.826 | -102.255 | IWSP | 40.2133 |
| MX94-039-2 | TRUE | NORTH_MX_1 | 27.311 | -102.397 | IWSQ | 38.0555 |
| MX94-040-5 | TRUE | NORTH_MX_2 | 27.317 | -102.400 | MX94-040-5 | 31.294 |
| MX94-041-5 | TRUE | NORTH_MX_1 | 26.950 | -102.083 | MX94-041-5 | 28.6153 |
| MX94-041-9 | TRUE | NORTH_MX_2 | 26.950 | -102.083 | 94-MX-041-9 | 28.2213 |
| MX94-042-6 | TRUE | NORTH_MX_2 | 26.950 | -102.083 | MX94-042-6 | 29.5516 |
| MX94-043-1 | TRUE | NORTH_MX_2 | 26.826 | -102.255 | IYPC | 65.1743 |
| MX94-044-6 | TRUE | NORTH_MX_2 | 28.221 | -100.724 | IXUR | 43.1024 |
| MX94-044-9 | TRUE | NORTH_MX_2 | 28.250 | -100.717 | 94-MX-044-9 | 29.1359 |
| MX94-045-7 | TRUE | NORTH_MX_2 | 26.826 | -102.255 | IWSS | 43.8379 |
| MX94-046-1 | TRUE | NORTH_MX_2 | 26.950 | -102.083 | 94-MX-046-1 | 28.3616 |
| MX94-046-8 | TRUE | NORTH_MX_2 | 26.826 | -102.255 | IXUS | 46.7267 |
| MX94-047-2 | TRUE | NORTH_MX_2 | 26.950 | -102.083 | MX94-047-2 | 29.1787 |
| MX94-048-2 | TRUE | NORTH_MX_2 | 26.826 | -102.255 | IWST | 46.6495 |
| MX94-049-2 | TRUE | NORTH_MX_1 | 28.221 | -100.724 | IWSU | 39.3666 |
| MX94-050-4 | TRUE | ADMIX | 28.250 | -100.717 | MX94-050-4 | 26.8743 |
| MX94-051-9 | TRUE | NORTH_MX_2 | 28.221 | -100.724 | IXFJ | 60.704 |
| MX94-052-3 | TRUE | NORTH_MX_2 | 27.317 | -102.400 | 94-MX-052-3 | 27.6781 |
| MX94-052-6 | TRUE | NORTH_MX_2 | 27.317 | -102.400 | MX94-052-6 | 29.5212 |
| MX94-053-8 | TRUE | NORTH_MX_2 | 28.221 | -100.724 | IWSW | 51.8403 |
| MX94-054-8 | TRUE | NORTH_MX_1 | 28.221 | -100.724 | IWSX | 44.5341 |
| MX94-055-5 | TRUE | NORTH_MX_2 | 28.221 | -100.724 | IWSY | 42.24 |

| | | | | | | |
|---|---|---|---|---|---|---|
| MX94-056-5 | TRUE | NORTH_MX_2 | 28.221 | -100.724 | IWSZ | 50.0798 |
| MX94-057-3 | TRUE | NORTH_MX_2 | 28.221 | -100.724 | IXFK | 41.2452 |
| MX94-058-6 | TRUE | NORTH_MX_2 | 28.250 | -100.717 | MX94-058-6 | 31.7736 |
| MX94-059-9 | TRUE | NORTH_MX_2 | 28.221 | -100.724 | IXFL | 36.1315 |
| MX94-060-3 | TRUE | NORTH_MX_2 | 28.221 | -100.724 | IWTA | 46.3879 |
| MX94-061-7 | TRUE | NORTH_MX_2 | 28.221 | -100.724 | IXUT | 39.4358 |
| MX94-062-9 | TRUE | NORTH_MX_1 | 28.221 | -100.724 | IWTB | 41.9387 |
| MX94-063-7 | TRUE | NORTH_MX_2 | 28.221 | -100.724 | IXFM | 47.4902 |
| MX94-064-2 | TRUE | NORTH_MX_2 | 28.221 | -100.724 | IXFN | 35.4089 |
| MX94-064-8 | TRUE | NORTH_MX_2 | 28.250 | -100.717 | 94-MX-064-8 | 27.7347 |
| MX94-065-7 | TRUE | NORTH_MX_2 | 28.250 | -100.717 | MX94-065-7 | 30.058 |
| MX94-066-7 | TRUE | NORTH_MX_2 | 25.350 | -103.133 | MX94-066-7 | 29.2962 |
| MX94-067-2 | TRUE | NORTH_MX_2 | 28.492 | -100.920 | IXUU | 36.2577 |
| MX94-068-4 | TRUE | NORTH_MX_2 | 28.492 | -100.920 | IXKP | 37.1864 |
| MX94-069-3 | TRUE | NORTH_MX_2 | 28.409 | -100.889 | IWTC | 31.1887 |
| MX94-070-4 | TRUE | NORTH_MX_1 | 25.350 | -103.133 | MX94-070-4 | 27.867 |
| MX94-071-6 | TRUE | NORTH_MX_2 | 28.492 | -100.920 | IWTD | 47.1124 |
| MX94-072-5 | TRUE | NORTH_MX_1 | 28.492 | -100.920 | IWTE | 46.5837 |
| MX94-073-2 | TRUE | NORTH_MX_2 | 28.492 | -100.920 | IXFP | 54.5022 |
| MX94-074-2 | TRUE | NORTH_MX_2 | 28.492 | -100.920 | IYPE | 46.0856 |
| MX94-075-2 | TRUE | NORTH_MX_2 | 25.441 | -102.175 | IWTF | 33.9241 |
| MX94-076-4 | TRUE | NORTH_MX_2 | 25.441 | -102.175 | IWTG | 49.4027 |
| MX94-077-3 | TRUE | NORTH_MX_2 | 25.441 | -102.175 | IWTH | 40.4439 |
| MX94-078-3 | TRUE | ADMIX | 25.441 | -102.175 | IXUW | 45.2836 |
| MX94-079-3 | TRUE | ADMIX | 25.350 | -103.133 | MX94-079-3 | 29.3186 |
| MX94-080-4 | TRUE | NORTH_MX_2 | 27.033 | -101.800 | MX94-080-4 | 27.5262 |
| MX94-081-7 | TRUE | NORTH_USA | 27.882 | -101.514 | IXFQ | 40.3332 |
| MX94-082-6 | TRUE | NORTH_MX_2 | 25.377 | -101.477 | IWTI | 36.3699 |
| MX94-083-1 | TRUE | TX_MX | 25.441 | -102.175 | IXFR | 36.994 |
| MX94-084-4 | TRUE | NORTH_MX_2 | 26.952 | -106.350 | IXFS | 38.4683 |
| MX94-085-6 | TRUE | NORTH_MX_1 | 25.441 | -102.175 | IXFT | 34.2645 |

| | | | | | | |
|---|---|---|---|---|---|---|
| MX94-087-4 | TRUE | NORTH_MX_2 | 25.441 | -102.175 | IWTJ | 40.7024 |
| MX94-088-3 | TRUE | NORTH_MX_2 | 25.441 | -102.175 | IWTK | 46.8621 |
| MX94-089-2 | TRUE | NORTH_MX_2 | 25.417 | -102.183 | MX94-089-2 | 28.8018 |
| MX94-090-2 | TRUE | NORTH_MX_2 | 25.417 | -102.183 | MX94-090-2 | 28.2235 |
| MX94-091-1 | TRUE | TX_MX | 25.441 | -102.175 | IXUX | 44.0676 |
| MX94-092-3 | TRUE | NORTH_MX_2 | 25.441 | -102.175 | IWTL | 37.0533 |
| MX94-093-3 | TRUE | NORTH_MX_2 | 25.550 | -100.967 | 94-MX-093-3 | 29.4632 |
| MX94-093-6 | TRUE | NORTH_MX_2 | 25.540 | -100.943 | IXFU | 45.4583 |
| MX94-094-2 | TRUE | NORTH_MX_2 | 25.441 | -102.175 | IXFW | 41.2156 |
| MX94-095-8 | TRUE | NORTH_MX_1 | 25.441 | -102.175 | IXFX | 40.1245 |
| MX94-096-5 | TRUE | TX_MX | 25.441 | -102.175 | IXUY | 42.382 |
| MX94-097-3 | TRUE | TX_MX | 25.441 | -102.175 | IXFY | 52.8259 |
| MX94-098-5 | TRUE | NORTH_MX_2 | 25.417 | -102.183 | MX94-098-5 | 29.9465 |
| MX94-099-5 | TRUE | NORTH_MX_2 | 25.417 | -102.183 | MX94-099-5 | 27.8679 |
| MX94-100-6 | TRUE | NORTH_MX_2 | 25.540 | -100.943 | IWTM | 53.0689 |
| MX94-101-6 | TRUE | ADMIX | 25.550 | -100.967 | MX94-101-6 | 30.2744 |
| MX94-102-2 | TRUE | CENTRAL_MX | 20.917 | -99.933 | MX94-102-2 | 29.8338 |
| MX94-104-3 | TRUE | TX_MX | 25.441 | -102.175 | IXFZ | 59.9432 |
| MX94-105-3 | TRUE | NORTH_MX_2 | 25.540 | -100.943 | IXGA | 48.8087 |
| MX94-106-1 | TRUE | TX_MX | 25.467 | -100.850 | MX94-106-1 | 30.8518 |
| MX94-107-2 | TRUE | NORTH_MX_2 | 25.377 | -101.477 | IXGB | 34.034 |
| MX94-108-4 | TRUE | NORTH_MX_2 | 25.441 | -102.175 | IWTN | 33.4353 |
| MX94-109-4 | TRUE | CENTRAL_MX | 21.053 | -99.815 | IXGC | 47.2929 |
| MX94-110-1 | TRUE | NORTH_MX_1 | 23.411 | -99.379 | IWTP | 45.5322 |
| MX94-111-4 | TRUE | ADMIX | 23.411 | -99.379 | IXGD | 36.2855 |
| MX94-112-4 | TRUE | TX_MX | 23.411 | -99.379 | IXGE | 46.1644 |
| MX94-113-7 | TRUE | ADMIX | 23.411 | -99.379 | IXGF | 45.7655 |
| MX94-114-2 | TRUE | NORTH_MX_2 | 24.778 | -104.453 | IWTQ | 37.7371 |
| MX94-115-5 | TRUE | ADMIX | 24.790 | -104.037 | MX94-115-5 | 28.826 |
| MX94-116-5 | TRUE | TX_MX | 24.033 | -104.667 | MX94-116-5 | 28.4222 |
| MX94-117-7 | TRUE | NORTH_MX_2 | 24.778 | -104.453 | IWTR | 39.5723 |

| | | | | | | |
|---|---|---|---|---|---|---|
| MX94-118-4 | TRUE | NORTH_MX_2 | 24.783 | -104.450 | MX94-118-4 | 29.0674 |
| MX94-119-4 | TRUE | NORTH_MX_2 | 24.778 | -104.453 | IWTS | 36.2125 |
| MX94-120-4 | TRUE | NORTH_MX_2 | 28.492 | -100.920 | IXGG | 49.4318 |
| MX94-121-2 | TRUE | TX_MX | 25.540 | -100.943 | IWTT | 39.0115 |
| MX94-121-5 | TRUE | TX_MX | 25.550 | -100.967 | MX94-121-5 | 31.1561 |
| MX94-122-2 | TRUE | CENTRAL_MX | 21.050 | -99.817 | MX94-122-2 | 27.1701 |
| MX94-123-2 | TRUE | TX_MX | 25.550 | -100.967 | MX94-123-2 | 28.9892 |
| MX94-126-3 | TRUE | CENTRAL_MX | 21.050 | -99.817 | MX94-126-3 | 27.2976 |
| MX94-127-6 | TRUE | CENTRAL_MX | 21.053 | -99.815 | IXUZ | 35.6056 |
| MX94-128-6 | TRUE | CENTRAL_MX | 21.053 | -99.815 | IWTU | 31.7227 |
| MX94-129-4 | TRUE | CENTRAL_MX | 21.050 | -99.817 | MX94-129-4 | 28.8477 |
| MX94-232-3 | TRUE | NORTH_MX_2 | 27.017 | -101.600 | MX94-232-3 | 27.9056 |
| MX94-234-3 | TRUE | NORTH_MX_2 | 27.029 | -101.592 | IWTW | 35.4461 |
| MX94-235-2 | TRUE | NORTH_MX_2 | 27.052 | -101.796 | IWTX | 39.6824 |
| MX94-236-5 | TRUE | NORTH_MX_2 | 27.033 | -101.800 | MX94-236-5 | 30.184 |
| MX94-237-7 | TRUE | NORTH_MX_2 | 27.052 | -101.796 | IXWA | 43.4314 |
| MX94-239-4 | TRUE | TX_MX | 28.026 | -105.290 | IWTY | 49.7879 |
| MX94-240-6 | TRUE | NORTH_MX_2 | 28.201 | -105.472 | IWTZ | 45.2507 |
| MX94-241-4 | TRUE | NORTH_MX_1 | 26.833 | -100.667 | MX94-241-4 | 28.8536 |
| MX94-242-6 | TRUE | NORTH_MX_1 | 26.837 | -100.667 | IXGH | 59.2913 |
| MX94-243-2 | TRUE | NORTH_MX_2 | 27.052 | -101.796 | IWUA | 33.4596 |
| MX94-244-5 | TRUE | NORTH_MX_2 | 27.052 | -101.796 | IXGI | 71.9017 |
| MX94-245-6 | TRUE | NORTH_MX_2 | 27.052 | -101.796 | IWUB | 39.8051 |
| MX94-246-7 | TRUE | ADMIX | 25.441 | -102.175 | IYPD | 51.5389 |
| MX94-247-8 | TRUE | NORTH_MX_2 | 25.441 | -102.175 | IXKR | 45.5978 |
| MX94-248-5 | TRUE | NORTH_MX_2 | 25.441 | -102.175 | IXGJ | 50.0266 |
| MX94-249-8 | TRUE | NORTH_MX_1 | 25.441 | -102.175 | IWUD | 38.1512 |
| MX94-250-4 | TRUE | NORTH_MX_2 | 25.441 | -102.175 | IXGK | 41.751 |
| MX94-251-6 | TRUE | NORTH_MX_2 | 25.441 | -102.175 | IXGL | 38.8991 |
| MX94-253-3 | TRUE | CENTRAL_MX | 21.050 | -99.817 | MX94-253-3 | 26.4666 |
| MX94-256-3 | TRUE | CENTRAL_MX | 21.050 | -99.817 | MX94-256-3 | 33.4001 |

| | | | | | | |
|---|---|---|---|---|---|---|
| MX94-257-1 | TRUE | CENTRAL_MX | 21.053 | -99.815 | IWUE | 35.852 |
| MX94-259-1 | TRUE | NORTH_MX_1 | 26.937 | -105.393 | IWUF | 37.8954 |
| MX94-260-5 | TRUE | NORTH_MX_1 | 26.533 | -100.500 | MX94-260-5 | 27.439 |
| MX94-261-1 | TRUE | NORTH_MX_2 | 26.533 | -100.500 | MX94-261-1 | 30.6298 |
| MX94-262-4 | TRUE | TX_MX | 26.530 | -100.502 | IXGM | 52.0698 |
| MX94-262-7 | TRUE | TX_MX | 26.530 | -100.502 | IXGN | 43.1997 |
| MX94-263-6 | TRUE | NORTH_MX_2 | 25.308 | -103.636 | IWUG | 36.2667 |
| MX94-264-7 | TRUE | NORTH_MX_2 | 25.017 | -100.083 | MX94-264-7 | 31.2824 |
| MX94-265-6 | TRUE | CENTRAL_MX | 21.050 | -99.817 | MX94-265-6 | 28.6432 |
| MX94-267-6 | TRUE | NORTH_MX_1 | 21.050 | -99.817 | MX94-267-6 | 27.1506 |
| MX94-270-4 | TRUE | NORTH_MX_1 | 21.050 | -99.817 | MX94-270-4 | 28.407 |
| MX94-271-7 | TRUE | ADMIX | 25.437 | -100.994 | IWUH | 43.5825 |
| MX94-272-2 | TRUE | TX_MX | 26.837 | -100.667 | IWUI | 33.5405 |
| MX94-273-6 | TRUE | NORTH_MX_2 | 29.300 | -100.917 | MX94-273-6 | 27.3478 |
| MX94-274-3 | TRUE | NORTH_MX_1 | 25.540 | -100.943 | IWUJ | 43.2887 |
| MX94-275-6 | TRUE | NORTH_MX_2 | 26.826 | -102.255 | IXWB | 43.0833 |
| MX94-277-2 | TRUE | NORTH_MX_2 | 26.950 | -102.083 | MX94-277-2 | 29.903 |
| MX94-278-1 | TRUE | CENTRAL_MX | 21.050 | -99.817 | MX94-278-1 | 29.4233 |
| MX94-279-5 | TRUE | ADMIX | 26.900 | -101.417 | MX94-279-5 | 28.9323 |
| MX94-280-5 | TRUE | NORTH_MX_2 | 25.441 | -102.175 | IXGP | 56.0244 |
| MX94-281-6 | TRUE | NORTH_MX_2 | 25.417 | -102.183 | MX94-281-6 | 28.348 |
| MX94-281-7 | TRUE | TX_MX | 25.441 | -102.175 | IWUK | 36.6314 |
| MX94-282-2 | TRUE | NORTH_MX_2 | 25.417 | -102.183 | MX94-282-2 | 31.0525 |
| MX94-282-4 | TRUE | NORTH_MX_2 | 25.441 | -102.175 | IWUL | 37.0345 |
| MX94-283-7 | TRUE | TX_MX | 21.053 | -99.815 | IWUM | 43.0998 |
| MX94-284-4 | TRUE | CENTRAL_MX | 21.050 | -99.817 | MX94-284-4 | 28.0215 |
| MX94-285-3 | TRUE | CENTRAL_MX | 21.050 | -99.817 | MX94-285-3 | 26.012 |
| MX94-286-6 | TRUE | CENTRAL_MX | 21.050 | -99.817 | MX94-286-6 | 26.9022 |
| MX94-287-3 | TRUE | CENTRAL_MX | 21.050 | -99.817 | MX94-287-3 | 29.3064 |
| MX94-290-4 | TRUE | NORTH_MX_2 | 25.441 | -102.175 | IWUN | 39.6925 |
| MX94-291-5 | TRUE | NORTH_MX_1 | 25.441 | -102.175 | IWUP | 43.6173 |

| | | | | | | |
|---|---|---|---|---|---|---|
| MX94-293-5 | TRUE | NORTH_MX_2 | 25.417 | -102.183 | MX94-293-5 | 29.1599 |
| MX94-294-3 | TRUE | NORTH_MX_2 | 27.004 | -101.724 | IXGQ | 40.7907 |
| MX94-295-3 | TRUE | NORTH_MX_1 | 25.417 | -102.183 | MX94-295-3 | 29.8075 |
| MX94-296-2 | TRUE | ADMIX | 25.016 | -100.074 | IWUR | 39.8585 |
| MX94-297-7 | TRUE | NORTH_MX_2 | 27.004 | -101.724 | IXGR | 46.2933 |
| MX94-298-2 | TRUE | NORTH_MX_2 | 27.004 | -101.724 | IXGS | 50.4278 |
| MX94-299-6 | TRUE | NORTH_MX_2 | 27.311 | -102.397 | IWUS | 38.7174 |
| MX94-300-7 | TRUE | NORTH_MX_2 | 27.311 | -102.397 | IWUT | 52.3598 |
| MX94-301-5 | TRUE | NORTH_MX_2 | 27.311 | -102.397 | IWUU | 45.2145 |
| MX94-302-4 | TRUE | NORTH_MX_1 | 27.017 | -101.600 | MX94-302-4 | 30.3836 |
| MX94-303-5 | TRUE | NORTH_MX_2 | 27.004 | -101.724 | IWUW | 43.1417 |
| MX94-304-2 | TRUE | NORTH_MX_2 | 25.350 | -103.133 | MX94-304-2 | 30.671 |
| MX94-305-1 | TRUE | TX_MX | 25.441 | -102.175 | IXGT | 39.9047 |
| MX94-306-1 | TRUE | ADMIX | 25.441 | -102.175 | IXKS | 37.7379 |
| MX94-307-2 | TRUE | ADMIX | 25.441 | -102.175 | IWUX | 45.5042 |
| MX94-308-3 | TRUE | TX_MX | 25.441 | -102.175 | IXKT | 31.1097 |
| MX94-309-1 | TRUE | NORTH_MX_1 | 25.441 | -102.175 | IWUY | 35.7845 |
| MX94-310-7 | TRUE | NORTH_MX_2 | 25.441 | -102.175 | IWUZ | 54.4426 |
| MX94-311-1 | TRUE | NORTH_MX_2 | 25.441 | -102.175 | IXGU | 34.7714 |
| MX94-312-3 | TRUE | NORTH_MX_2 | 25.441 | -102.175 | IWWA | 44.3694 |
| MX94-313-3 | TRUE | NORTH_MX_2 | 25.417 | -102.183 | MX94-313-3 | 28.9887 |
| MX94-314-5 | TRUE | NORTH_MX_2 | 25.417 | -102.183 | MX94-314-5 | 29.7359 |
| MX94-315-1 | TRUE | TX_MX | 25.441 | -102.175 | IXGW | 40.933 |
| MX94-316-6 | TRUE | TX_MX | 25.016 | -100.074 | IXWC | 49.6773 |
| MX94-318-2 | TRUE | ADMIX | 25.016 | -100.074 | IXKU | 32.3192 |
| MX94-319-4 | TRUE | NORTH_MX_2 | 25.016 | -100.074 | IWWB | 50.2554 |
| MX94-320-3 | TRUE | ADMIX | 25.016 | -100.074 | IWWC | 52.7169 |
| MX94-320-5 | TRUE | ADMIX | 25.017 | -100.083 | 94-MX-320-5 | 27.6973 |
| MX94-321-4 | TRUE | TX_MX | 26.530 | -100.502 | IWWD | 43.4199 |
| MX94-322-5 | TRUE | NORTH_MX_2 | 25.016 | -100.074 | IWWE | 44.7645 |
| MX94-323-2 | TRUE | ADMIX | 26.530 | -100.502 | IXWD | 43.4276 |

| | | | | | | |
|---|---|---|---|---|---|---|
| MX94-324-3 | TRUE | NORTH_MX_1 | 26.530 | -100.502 | IWWF | 40.0417 |
| MX94-325-3 | TRUE | NORTH_MX_1 | 26.533 | -100.500 | MX94-325-3 | 29.8815 |
| MX94-326-1 | TRUE | NORTH_MX_1 | 24.608 | -100.490 | IWWG | 41.7025 |
| MX94-327-4 | TRUE | ADMIX | 25.016 | -100.074 | IWWH | 45.9342 |
| MX94-328-4 | TRUE | NORTH_MX_2 | 26.530 | -100.502 | IXWE | 40.5582 |
| MX94-329-6 | TRUE | ADMIX | 25.812 | -100.599 | IWWI | 39.273 |
| MX94-329-9 | TRUE | NORTH_MX_1 | 25.850 | -100.500 | 94-MX-329-9 | 31.382 |
| MX94-330-3 | TRUE | TX_MX | 25.812 | -100.599 | IXGX | 44.4256 |
| MX94-331-2 | TRUE | TX_MX | 24.608 | -100.490 | IWWJ | 52.5073 |
| MX94-332-2 | TRUE | NORTH_MX_2 | 26.837 | -100.667 | IWWK | 37.6031 |
| MX94-333-2 | TRUE | NORTH_MX_1 | 26.837 | -100.667 | IWWL | 40.8328 |
| MX94-334-1 | TRUE | NORTH_MX_2 | 25.222 | -104.113 | IXGY | 64.8935 |
| MX94-336-3 | TRUE | NORTH_MX_1 | 25.233 | -104.133 | MX94-336-3 | 27.4985 |
| Nacono | FALSE | NA | NA | NA | IWWN | 35.9377 |
| Navaho | FALSE | NA | NA | NA | Navaho | 29.8449 |
| NC-2A | FALSE | NA | NA | NA | IYPX | 43.0123 |
| NC-2B | FALSE | NA | NA | NA | IYPY | 80.0866 |
| NC-2B | FALSE | NA | NA | NA | NC-2B | 25.9979 |
| NC-4 | FALSE | NA | 43.255 | -79.077 | IYAH | 36.0834 |
| Nelson | FALSE | NA | 30.403 | -89.494 | Nelson | 37.7935 |
| Neuman | FALSE | NA | 29.820 | -95.958 | IXWK | 45.9121 |
| Niblack | TRUE | TX_MX | 38.679 | -87.419 | Niblack | 27.879 |
| Norton | TRUE | NORTH_USA | 39.313 | -90.820 | Norton | 25.8272 |
| Nueces | TRUE | NORTH_MX_2 | 28.716 | -99.811 | Nueces | 32.2457 |
| Nugget | TRUE | TX_MX | 31.837 | -98.396 | IXWL | 43.8561 |
| Number 54 | FALSE | NA | 34.360 | -106.100 | Number-54 | 29.3922 |
| Oconee | FALSE | NA | 28.593 | -98.991 | Oconee | 28.2963 |
| Odom | FALSE | NA | 33.532 | -93.561 | Odom | 29.4542 |
| Oklahoma | TRUE | NORTH_USA | 34.239 | -97.083 | IXWM | 43.3134 |
| Old Woman | TRUE | NORTH_USA | 40.862 | -91.073 | IYAJ | 51.7733 |
| Oliver | TRUE | TX_MX | 30.495 | -99.732 | IXKX | 48.2409 |

| | | | | | | |
|---|---|---|---|---|---|---|
| Onliwon | TRUE | TX_MX | 31.250 | -98.596 | IYAK | 41.1702 |
| Osborne | TRUE | TX_MX | 30.821 | -98.579 | IYAL | 37.2923 |
| Outpost | TRUE | TX_MX | 32.825 | -101.379 | IYPF | 58.4124 |
| Owens | FALSE | NA | 34.416 | -90.523 | Owens | 27.1542 |
| PA Big Tree | FALSE | NA | 40.212 | -77.013 | IYAM | 45.7901 |
| Pabst | FALSE | NA | NA | NA | Pabst | 27.4461 |
| Patrick | TRUE | NORTH_USA | 36.235 | -95.691 | Patrick | 27.4973 |
| Pawnee | FALSE | NA | NA | NA | JAED | 105.667 |
| Pearce | TRUE | TX_MX | 30.344 | -97.886 | IYAN | 41.4687 |
| Pensacola Cluster | FALSE | NA | 30.421 | -87.217 | Pensacola-Cluster | 28.0194 |
| Perfect | FALSE | NA | 29.667 | -96.000 | Perfect | 27.4409 |
| Peruque | TRUE | NORTH_USA | 38.879 | -90.626 | IYAP | 34.7819 |
| Pioneer (AL) | FALSE | NA | 30.399 | -87.776 | Pioneer-AL- | 31.0672 |
| Podsednik | FALSE | NA | 32.689 | -97.168 | IYAQ | 55.7409 |
| Pointe Coupee 2 | TRUE | NORTH_USA | 30.734 | -91.433 | IXWP | 53.7298 |
| Posey | TRUE | NORTH_USA | 38.389 | -87.555 | IXWQ | 37.7848 |
| Prilop | TRUE | TX_MX | 29.424 | -96.940 | IYAR | 46.5602 |
| Ramsey Mediumshell | FALSE | NA | 30.411 | -88.828 | Ramsey-Mediumshell | 32.2151 |
| Randall | FALSE | NA | 29.717 | -82.140 | IYAS | 36.2179 |
| RDM-LL-2 | FALSE | NA | NA | NA | IXKZ | 33.4341 |
| RDM-LL-8 | FALSE | NA | NA | NA | IXWR | 51.1172 |
| Red Cobb | TRUE | NORTH_USA | 34.603 | -92.009 | Red-Cobb | 28.2037 |
| Red Nut | FALSE | NA | NA | NA | Red-Nut | 27.3349 |
| RHS Special 1 | FALSE | NA | NA | NA | RHS-Special-1 | 26.4632 |
| RHS Special 2 | FALSE | NA | NA | NA | RHS-Special-2 | 27.2152 |
| Rice | FALSE | NA | NA | NA | Rice | 28.3635 |
| Ripe Early | TRUE | NORTH_USA | 35.520 | -97.240 | IYAT | 42.4071 |
| Risien 1 | FALSE | NA | 31.247 | -98.594 | Risien-1 | 27.0755 |
| Riverside | FALSE | NA | 31.379 | -98.661 | IXLA | 46.9173 |
| Rome | FALSE | NA | NA | NA | Rome | 30.8429 |

| | | | | | | |
|---|---|---|---|---|---|---|
| Roth | TRUE | TX_MX | 29.248 | -97.329 | IYAU | 32.4738 |
| RS 66-23 | FALSE | NA | 28.593 | -98.991 | RS-66-23 | 29.0455 |
| RS11-1 | FALSE | NA | 28.593 | -98.991 | IYCF | 45.2152 |
| RuCox | TRUE | NORTH_USA | 30.087 | -95.419 | RuCox | 26.6309 |
| Russell | FALSE | NA | 30.411 | -88.828 | Russell | 26.4848 |
| Salado | FALSE | NA | 28.556 | -98.983 | Salado | 27.9051 |
| Salopek | FALSE | NA | NA | NA | Salopek | 27.7092 |
| San Saba Improved | FALSE | NA | 31.250 | -98.596 | San-Saba-Improved | 27.0645 |
| SanFelipe | TRUE | NORTH_USA | 29.371 | -100.884 | IYPN | 60.2927 |
| SanSaba | TRUE | TX_MX | 31.250 | -98.596 | IYAW | 51.4132 |
| Schaeffer | FALSE | NA | 30.411 | -88.828 | Schaeffer | 39.6228 |
| Schley | FALSE | NA | 30.366 | -88.556 | IXWS | 39.2621 |
| Schutz 1 | TRUE | TX_MX | 29.482 | -97.583 | Schutz-1 | 29.3301 |
| Schutz 2 | TRUE | TX_MX | 29.485 | -97.586 | IYAX | 42.6978 |
| Seminole | FALSE | NA | 30.411 | -83.953 | Seminole | 28.7733 |
| Shawnee | FALSE | NA | 28.594 | -98.991 | Shawnee | 27.6178 |
| Shepherd | TRUE | NORTH_USA | 39.158 | -94.496 | IYAY | 38.4073 |
| Shoals West | FALSE | NA | NA | NA | ShoalsWest | 27.4463 |
| Shoshoni | FALSE | NA | NA | NA | IWWQ | 42.7564 |
| Siepmann | FALSE | NA | NA | NA | Siepmann- | 28.7534 |
| Silverback | FALSE | NA | 35.520 | -97.240 | Silverback | 29.086 |
| Sioux | FALSE | NA | NA | NA | Sioux | 28.6466 |
| Snaps | TRUE | NORTH_USA | 42.206 | -90.369 | IXWT | 49.8557 |
| Snodgrass | FALSE | NA | NA | NA | Snodgrass | 26.2949 |
| Soefje #1 | FALSE | NA | NA | NA | Soefje-1 | 29.4635 |
| Spence (MO) | TRUE | NORTH_USA | 39.426 | -93.131 | IXWW | 37.7212 |
| Squirrel's Delight | TRUE | TX_MX | 31.249 | -98.597 | IYAZ | 42.8157 |
| Starking_Hardy_Giant | TRUE | NORTH_USA | 39.371 | -93.065 | IYBA | 38.145 |
| Steuk | TRUE | NORTH_USA | 38.064 | -94.238 | Steuk | 27.6996 |
| Stuart | FALSE | NA | 30.683 | -88.150 | IXWX | 39.8459 |

| | | | | | | |
|---|---|---|---|---|---|---|
| Stubbs Early | FALSE | NA | NA | NA | Stubbs-Early | 26.3531 |
| Success | FALSE | NA | 30.411 | -88.828 | IYBB | 62.7575 |
| Sugar Loaf | FALSE | NA | NA | NA | Sugar-Loaf | 26.673 |
| Sumner | FALSE | NA | 31.444 | -83.516 | IYBC | 44.329 |
| Supreme | FALSE | NA | 28.593 | -98.991 | Supreme | 26.6188 |
| Surprize | FALSE | NA | 30.406 | -87.684 | Surprize | 26.4765 |
| Syrup Mill | FALSE | NA | 30.694 | -88.043 | Syrup-Mill | 26.5082 |
| Talbot | TRUE | TX_MX | 30.838 | -100.125 | IXGZ | 74.0236 |
| Tarrant | FALSE | NA | 30.982 | -97.516 | Tarrant | 27.0949 |
| Taylor Dwarf | FALSE | NA | 32.473 | -98.945 | Taylor-Dwarf | 29.2879 |
| Teche | FALSE | NA | 29.975 | -91.756 | Teche | 29.8654 |
| Tejas | FALSE | NA | 28.594 | -98.991 | Tejas | 28.3052 |
| Tetraploid 2 | FALSE | NA | NA | NA | IXLC | 50.7962 |
| Texas 60 | FALSE | NA | 31.250 | -98.596 | Texas-60 | 25.4231 |
| Tiemann | TRUE | TX_MX | 29.896 | -96.865 | IYBD | 38.0336 |
| Tinker | FALSE | NA | 31.180 | -85.280 | Tinker | 26.8812 |
| Tiny Tim | TRUE | NORTH_USA | 38.970 | -90.832 | IXWZ | 46.5193 |
| Tissue Paper | TRUE | NORTH_USA | 37.164 | -94.843 | IYBE | 46.2497 |
| TN Big Tree | FALSE | NA | 35.943 | -83.178 | IXWY | 55.0752 |
| Tobacco Barn | FALSE | NA | NA | NA | Tobacco-Barn | 25.4178 |
| U/E 2-8 | FALSE | NA | NA | NA | U-E-2-8 | 31.4179 |
| VanDeman | FALSE | NA | 30.087 | -90.905 | IYBF | 75.5696 |
| VC1-68 | FALSE | NA | 33.508 | -112.316 | IYPP | 43.9018 |
| W Iowa St. | FALSE | NA | NA | NA | W-Iowa-St- | 28.3736 |
| Waco | FALSE | NA | NA | NA | IWWR | 34.9577 |
| Waco Wonder | FALSE | NA | 31.541 | -97.191 | Waco-Wonder | 26.3909 |
| Wallops Island | FALSE | NA | 37.882 | -75.437 | IXXA | 37.6596 |
| Warden | FALSE | NA | NA | NA | Warden | 28.9692 |
| Warren 346 | TRUE | NORTH_USA | 39.779 | -93.370 | IYBG | 51.3503 |
| Waukeenah | FALSE | NA | 30.411 | -83.953 | IXLE | 36.6564 |
| Weinman | TRUE | TX_MX | 29.888 | -96.104 | IXLF | 32.4583 |

| | | | | | | |
|---|---|---|---|---|---|---|
| Weschcke Survivor | FALSE | NA | NA | NA | Weschcke-Survivor | 35.1238 |
| Western | FALSE | NA | 31.251 | -98.597 | IYBI | 48.4044 |
| Wichita | FALSE | NA | NA | NA | IYBJ | 41.5196 |
| Wiese | TRUE | NORTH_USA | 39.385 | -93.221 | IYBK | 59.2716 |
| Williamson | TRUE | TX_MX | 34.404 | -96.826 | IYBL | 39.0262 |
| Winner | TRUE | TX_MX | 31.375 | -98.671 | IYBM | 53.9194 |
| Witte (IA) | TRUE | NORTH_USA | 40.840 | -91.161 | Witte-IA- | 28.559 |
| WN-10 | TRUE | TX_MX | 33.039 | -98.600 | WN-10 | 25.8294 |
| WN-17 | TRUE | TX_MX | 33.039 | -98.600 | WN-17 | 28.5757 |
| WN-5 | TRUE | TX_MX | 33.039 | -98.600 | WN-5 | 27.0072 |
| WN-6 | TRUE | TX_MX | 33.039 | -98.600 | WN-6 | 28.9033 |
| Woodard | FALSE | NA | 31.475 | -83.643 | Woodard | 28.0332 |
| Woodroof | FALSE | NA | NA | NA | Woodroof | 28.32 |
| Woodside Early | FALSE | NA | 31.245 | -92.376 | Woodside-Early | 36.0363 |
| WS-10 | TRUE | TX_MX | 33.038 | -98.600 | WS-10 | 27.0016 |
| WS-15 | TRUE | TX_MX | 33.038 | -98.600 | WS-15 | 27.4248 |
| WS-21 | TRUE | TX_MX | 33.038 | -98.600 | WS-21 | 27.5194 |
| WS-5 | TRUE | TX_MX | 33.038 | -98.600 | WS-5 | 24.8348 |
| WSE | FALSE | NA | NA | NA | WSE | 25.973 |
| Zajicek | TRUE | TX_MX | 32.748 | -97.196 | Zajicek | 28.9621 |
| Zinner | FALSE | NA | 30.519 | -87.808 | IYBP | 41.5085 |
| Kanza | FALSE | NA | NA | NA | IWWS | 35.1842 |
| Lakota | FALSE | NA | NA | NA | IKFU | 73.6889 |
| Major | TRUE | NORTH_USA | 37.899 | -87.486 | IYBQ | 57.5862 |
| Osage | FALSE | NA | NA | NA | IWWP | 36.2837 |
| 2015-02-0001 | FALSE | NA | NA | NA | IYQK | 54.1986 |
| 2015-01-0001 | FALSE | NA | NA | NA | IYQH | 58.3984 |
| I-17-1 | FALSE | NA | NA | NA | I-17-1 | 29.6246 |

[1]Gene pool designation for native pecan samples based on ADMIXTURE results. Samples with <50% ancestry from a single gene pool are designated Admixed. Breeding samples not assigned a gene pool. CENTRAL_MX = Central Mexico; NORTH_MX_1 = Northern Mexico 1;

NORTH_MX_2 = Northern Mexico 2; NORTH_USA = Northern USA; TX_MX = Texas and Mexico.
[2]Sample ID in VCF genotype files.

**Table S4: Pecan samples genotyped using Genotype-by-Sequencing (GBS)**

| Plant ID | VCF_ID[1] | Native Sample | Latitude | Longitude | SNP Genotypes[2] |
|---|---|---|---|---|---|
| Success | NMSU0001 | FALSE | NA | NA | 19282 |
| Jackson | NMSU0002 | FALSE | NA | NA | 18995 |
| Curtis | NMSU0003 | FALSE | NA | NA | 19064 |
| Farley | NMSU0004 | FALSE | NA | NA | 21089 |
| A-93 | NMSU0005 | FALSE | NA | NA | 20625 |
| Brooks | NMSU0006 | FALSE | NA | NA | 19450 |
| Kellers | NMSU0007 | FALSE | NA | NA | 19368 |
| 1975-08-0005 | NMSU0008 | FALSE | NA | NA | 17779 |
| 1977-11-0028 | NMSU0009 | FALSE | NA | NA | 19518 |
| 1982-15-0002 | NMSU0010 | FALSE | NA | NA | 19178 |
| 1982-15-0021 | NMSU0011 | FALSE | NA | NA | 22224 |
| 1979-07-0044 | NMSU0012 | FALSE | NA | NA | 18707 |
| Shawnee | NMSU0020 | FALSE | NA | NA | 18088 |
| Cheyenne | NMSU0021 | FALSE | NA | NA | 22242 |
| Nacono | NMSU0022 | FALSE | NA | NA | 20955 |
| Cherokee | NMSU0023 | FALSE | NA | NA | 20076 |
| Chickasaw | NMSU0024 | FALSE | NA | NA | 22325 |
| Mandan | NMSU0027 | FALSE | NA | NA | 19667 |
| Apalachee | NMSU0028 | FALSE | NA | NA | 17495 |
| Tejas | NMSU0030 | FALSE | NA | NA | 22236 |
| Pawnee | NMSU0031 | FALSE | NA | NA | 21223 |
| Houma | NMSU0033 | FALSE | NA | NA | 21174 |
| Oconee | NMSU0034 | FALSE | NA | NA | 21013 |
| Navaho | NMSU0035 | FALSE | NA | NA | 20835 |
| Lipan | NMSU0037 | FALSE | NA | NA | 18545 |
| Creek | NMSU0038 | FALSE | NA | NA | 21006 |

| | | | | | |
|---|---|---|---|---|---|
| Hopi | NMSU0039 | FALSE | NA | NA | 14572 |
| MX94-108-4 | NMSU0040 | TRUE | 25.441 | -102.175 | 21024 |
| MX94-320-3 | NMSU0046 | TRUE | 25.016 | -100.074 | 19183 |
| MX94-043-1 | NMSU0041 | TRUE | 26.826 | -102.255 | 20750 |
| MX94-092-3 | NMSU0042 | TRUE | 25.441 | -102.175 | 18972 |
| Burkett1 | NMSU0043 | FALSE | NA | NA | 20284 |
| Giles1 | NMSU0044 | FALSE | NA | NA | 23453 |
| MX94-234-3 | NMSU0045 | TRUE | 27.029 | -101.592 | 23985 |
| MX94-100-6 | NMSU0047 | TRUE | 25.540 | -100.943 | 20042 |
| MX94-056-5 | NMSU0048 | TRUE | 28.221 | -100.724 | 20087 |
| MX94-299-6 | NMSU0049 | TRUE | 27.311 | -102.397 | 23820 |
| MX94-002-9 | NMSU0050 | TRUE | 27.929 | -101.304 | 24541 |
| MX94-245-6 | NMSU0051 | TRUE | 27.052 | -101.796 | 24413 |
| MX94-004-5 | NMSU0052 | TRUE | 27.604 | -100.728 | 24359 |
| MX94-327-4 | NMSU0053 | TRUE | 25.016 | -100.074 | 24097 |
| MX94-008-2 | NMSU0054 | TRUE | 27.604 | -100.728 | 23419 |
| MX94-076-4 | NMSU0055 | TRUE | 25.441 | -102.175 | 24017 |
| MX94-087-4 | NMSU0056 | TRUE | 25.441 | -102.175 | 23238 |
| MX94-271-7 | NMSU0057 | TRUE | 25.437 | -100.994 | 24698 |
| MX94-281-7 | NMSU0058 | TRUE | 25.441 | -102.175 | 19921 |
| MX94-048-2 | NMSU0059 | TRUE | 26.826 | -102.255 | 24039 |
| MX94-319-4 | NMSU0061 | TRUE | 25.016 | -100.074 | 23789 |
| MX94-072-5 | NMSU0063 | TRUE | 28.492 | -100.920 | 22023 |
| MX94-023-7 | NMSU0065 | TRUE | 26.784 | -101.430 | 22549 |
| MX94-303-5 | NMSU0066 | TRUE | 27.004 | -101.724 | 24345 |
| MX94-243-2 | NMSU0067 | TRUE | 27.052 | -101.796 | 22763 |
| MX94-246-7 | NMSU0068 | TRUE | 25.441 | -102.175 | 23265 |
| Major5 | NMSU0069 | FALSE | NA | NA | 23900 |
| MX94-249-8 | NMSU0071 | TRUE | 25.441 | -102.175 | 23791 |
| MX94-030-6 | NMSU0072 | TRUE | 26.826 | -102.255 | 24451 |
| MX94-054-8 | NMSU0073 | TRUE | 28.221 | -100.724 | 24290 |

| | | | | | |
|---|---|---|---|---|---|
| MX94-053-8 | NMSU0074 | TRUE | 28.221 | -100.724 | 24291 |
| MX94-322-5 | NMSU0076 | TRUE | 25.016 | -100.074 | 23321 |
| MX94-292-4 | NMSU0077 | TRUE | 27.311 | -102.397 | 24754 |
| MX94-307-2 | NMSU0078 | TRUE | 25.441 | -102.175 | 24319 |
| MX94-121-2 | NMSU0079 | TRUE | 25.540 | -100.943 | 23291 |
| MX94-291-5 | NMSU0080 | TRUE | 25.441 | -102.175 | 24330 |
| MX94-128-6 | NMSU0081 | TRUE | 21.053 | -99.815 | 24307 |
| MX94-326-1 | NMSU0082 | TRUE | 24.608 | -100.490 | 23851 |
| Peruque4 | NMSU0084 | FALSE | NA | NA | 23778 |
| MX94-062-9 | NMSU0085 | TRUE | 28.221 | -100.724 | 23375 |
| MX94-114-2 | NMSU0086 | TRUE | 24.778 | -104.453 | 23896 |
| MX94-003-4 | NMSU0087 | TRUE | 29.069 | -100.679 | 24252 |
| MX94-060-3 | NMSU0088 | TRUE | 28.221 | -100.724 | 24090 |
| MX94-300-7 | NMSU0089 | TRUE | 27.311 | -102.397 | 23868 |
| MX94-235-2 | NMSU0090 | TRUE | 27.052 | -101.796 | 22479 |
| MX94-088-3 | NMSU0091 | TRUE | 25.441 | -102.175 | 23397 |
| MX94-036-4 | NMSU0092 | TRUE | 26.826 | -102.255 | 23882 |
| MX94-257-1 | NMSU0095 | TRUE | 21.053 | -99.815 | 23504 |
| MX94-039-2 | NMSU0096 | TRUE | 27.311 | -102.397 | 24054 |
| MX94-049-2 | NMSU0097 | TRUE | 28.221 | -100.724 | 23951 |
| MX94-312-3 | NMSU0099 | TRUE | 25.441 | -102.175 | 21491 |
| MX94-037-3 | NMSU0102 | TRUE | 26.826 | -102.255 | 24195 |
| MX94-309-1 | NMSU0105 | TRUE | 25.441 | -102.175 | 23867 |
| MX94-119-4 | NMSU0107 | TRUE | 24.778 | -104.453 | 23894 |
| MX94-077-3 | NMSU0108 | TRUE | 25.441 | -102.175 | 22878 |
| MX94-329-6 | NMSU0109 | TRUE | 25.812 | -100.599 | 23097 |
| MX94-071-6 | NMSU0110 | TRUE | 28.492 | -100.920 | 24409 |
| MX94-026-5 | NMSU0117 | TRUE | 27.311 | -102.397 | 23229 |
| MX94-290-4 | NMSU0118 | TRUE | 25.441 | -102.175 | 23837 |
| MX94-075-2 | NMSU0119 | TRUE | 25.441 | -102.175 | 22892 |
| MX94-301-5 | NMSU0122 | TRUE | 27.311 | -102.397 | 24357 |

| | | | | | |
|---|---|---|---|---|---|
| MX94-263-6 | NMSU0123 | TRUE | 25.308 | -103.636 | 23843 |
| MX94-296-2 | NMSU0125 | TRUE | 25.016 | -100.074 | 24073 |
| MX94-007-6 | NMSU0127 | TRUE | 26.826 | -102.255 | 23751 |
| MX94-001-8 | NMSU0128 | TRUE | 28.492 | -100.920 | 22007 |
| MX94-326-4 | NMSU0132 | TRUE | 24.608 | -100.490 | 21385 |
| MX94-057-3 | NMSU0134 | TRUE | 28.221 | -100.724 | 12968 |
| MX94-078-3 | NMSU0135 | TRUE | 25.441 | -102.175 | 18519 |
| MX94-005-6 | NMSU0136 | TRUE | 27.929 | -101.304 | 13271 |
| MX94-315-1 | NMSU0137 | TRUE | 25.441 | -102.175 | 15614 |
| MX94-242-6 | NMSU0140 | TRUE | 26.837 | -100.667 | 15115 |
| MX94-044-6 | NMSU0141 | TRUE | 28.221 | -100.724 | 15708 |
| LittleJewel3 | NMSU0142 | FALSE | NA | NA | 16214 |
| MX94-105-3 | NMSU0145 | TRUE | 25.540 | -100.943 | 18949 |
| MX94-308-3 | NMSU0146 | TRUE | 25.441 | -102.175 | 14692 |
| MX94-113-7 | NMSU0147 | TRUE | 23.411 | -99.379 | 13733 |
| MX94-258-6 | NMSU0149 | TRUE | 21.053 | -99.815 | 16527 |
| MX94-104-3 | NMSU0152 | TRUE | 25.441 | -102.175 | 22553 |
| MX94-280-5 | NMSU0153 | TRUE | 25.441 | -102.175 | 14370 |
| MX94-316-6 | NMSU0155 | TRUE | 25.016 | -100.074 | 16413 |
| MX94-334-2 | NMSU0156 | TRUE | 25.222 | -104.113 | 13862 |
| MX94-009-2 | NMSU0157 | TRUE | 26.784 | -101.430 | 21089 |
| MX94-127-6 | NMSU0158 | TRUE | 21.053 | -99.815 | 15372 |
| MX94-250-4 | NMSU0159 | TRUE | 25.441 | -102.175 | 16151 |
| MX94-018-5 | NMSU0160 | TRUE | 26.826 | -102.255 | 15699 |
| MX94-083-1 | NMSU0161 | TRUE | 25.441 | -102.175 | 13180 |
| MX94-328-4 | NMSU0162 | TRUE | 26.530 | -100.502 | 21679 |
| MX94-237-7 | NMSU0163 | TRUE | 27.052 | -101.796 | 16474 |
| MX94-107-2 | NMSU0164 | TRUE | 25.377 | -101.477 | 17260 |
| MX94-248-5 | NMSU0165 | TRUE | 25.441 | -102.175 | 17285 |
| MX94-073-2 | NMSU0167 | TRUE | 28.492 | -100.920 | 14454 |
| MX94-311-1 | NMSU0169 | TRUE | 25.441 | -102.175 | 14377 |

| | | | | | |
|---|---|---|---|---|---|
| MX94-067-2 | NMSU0170 | TRUE | 28.492 | -100.920 | 21938 |
| MX94-064-2 | NMSU0171 | TRUE | 28.221 | -100.724 | 14443 |
| MX94-091-1 | NMSU0172 | TRUE | 25.441 | -102.175 | 20410 |
| MX94-305-1 | NMSU0174 | TRUE | 25.441 | -102.175 | 17412 |
| MX94-323-2 | NMSU0175 | TRUE | 26.530 | -100.502 | 20802 |
| MX94-020-4 | NMSU0177 | TRUE | 27.004 | -101.724 | 14373 |
| MX94-013-6 | NMSU0182 | TRUE | 28.492 | -100.920 | 14072 |
| MX94-095-8 | NMSU0183 | TRUE | 25.441 | -102.175 | 13210 |
| MX94-063-7 | NMSU0186 | TRUE | 28.221 | -100.724 | 21235 |
| MX94-094-2 | NMSU0188 | TRUE | 25.441 | -102.175 | 18585 |
| MX94-093-6 | NMSU0190 | TRUE | 25.540 | -100.943 | 19398 |
| MX94-120-4 | NMSU0191 | TRUE | 28.492 | -100.920 | 19315 |
| MX94-247-8 | NMSU0192 | TRUE | 25.441 | -102.175 | 15725 |
| VC1-68-14 | NMSU0193 | FALSE | NA | NA | 17115 |
| MX94-298-2 | NMSU0194 | TRUE | 27.004 | -101.724 | 16154 |
| MX94-074-2 | NMSU0195 | TRUE | 28.492 | -100.920 | 15758 |
| MX94-109-4 | NMSU0196 | TRUE | 21.053 | -99.815 | 13562 |
| 87-MX-4-3.1 | NMSU0199 | TRUE | 20.485 | -99.220 | 14619 |
| 86-TX-1-3.1 | NMSU0200 | TRUE | 29.486 | -97.466 | 18469 |
| 86-TX-1-4.1 | NMSU0201 | TRUE | 29.490 | -97.466 | 21696 |
| 87-MX-1-1.1 | NMSU0202 | TRUE | 21.667 | -99.539 | 21175 |
| 86-TX-1-5.1 | NMSU0203 | TRUE | 29.495 | -97.460 | 22001 |
| 86-TX-5-5.1 | NMSU0204 | TRUE | 31.481 | -100.467 | 17830 |
| 87-MX-4-5.1 | NMSU0205 | TRUE | 20.486 | -99.221 | 19425 |
| 87-MX-1-1.2 | NMSU0206 | TRUE | 21.667 | -99.539 | 23510 |
| 87-MX-1-2.1 | NMSU0207 | TRUE | 21.672 | -99.539 | 22768 |
| 87-MX-5-4.1 | NMSU0208 | TRUE | 23.368 | -99.479 | 24124 |
| 87-MX-4-2.1 | NMSU0209 | TRUE | 20.484 | -99.222 | 23820 |
| 87-MX-4-3.2 | NMSU0210 | TRUE | 20.485 | -99.220 | 24290 |
| 87-MX-1-1.3 | NMSU0211 | TRUE | 21.667 | -99.539 | 22350 |
| 87MX-5-3.1 | NMSU0212 | TRUE | 23.404 | -99.382 | 23555 |

| | | | | | |
|---|---|---|---|---|---|
| 86-TX-5-5.2 | NMSU0213 | TRUE | 31.481 | -100.467 | 20189 |
| 86-KS-2-4.1 | NMSU0216 | TRUE | 37.062 | -95.060 | 19965 |
| 87-MX-1-1.4 | NMSU0217 | TRUE | 21.667 | -99.539 | 23657 |
| 87-MX-2-1.1 | NMSU0218 | TRUE | 19.891 | -103.586 | 24068 |
| 86-TX-5-4.1 | NMSU0219 | TRUE | 31.482 | -100.471 | 22441 |
| 86-TX-5-2.1 | NMSU0221 | TRUE | 31.354 | -100.209 | 22531 |
| 89-ILL-OK-5.1 | NMSU0222 | TRUE | 34.361 | -96.635 | 24057 |
| 86-TX-5-1.1 | NMSU0223 | TRUE | 31.352 | -100.207 | 23353 |
| 86-KS-2-5.2 | NMSU0224 | TRUE | 37.025 | -95.046 | 20251 |
| 86-TX-5-4.2 | NMSU0225 | TRUE | 31.482 | -100.471 | 23707 |
| 86-TX-2-2.1 | NMSU0226 | TRUE | 28.910 | -99.781 | 24250 |
| 87-MX-2-2.1 | NMSU0227 | TRUE | 19.893 | -103.588 | 24029 |
| 94-MX-093-3 | NMSU0228 | TRUE | 25.540 | -100.943 | 22886 |
| 86-MO-1-1.1 | NMSU0229 | TRUE | 39.758 | -93.608 | 24093 |
| 86-TN-1-1.1 | NMSU0230 | TRUE | 36.377 | -89.501 | 24328 |
| 94-MX-041-9 | NMSU0231 | TRUE | 26.826 | -102.255 | 23397 |
| 87-MX-1-4.1 | NMSU0233 | TRUE | 21.678 | -99.543 | 23506 |
| 87-MX-1-2.2 | NMSU0234 | TRUE | 21.672 | -99.539 | 19418 |
| 86-TX-4-5.1 | NMSU0235 | TRUE | 30.025 | -101.169 | 23515 |
| 86-TX-2-2.2 | NMSU0236 | TRUE | 28.910 | -99.781 | 22204 |
| 86-TX-3-1.1 | NMSU0237 | TRUE | 29.230 | -100.482 | 18362 |
| 87-MX-4-2.2 | NMSU0238 | TRUE | 20.484 | -99.222 | 22944 |
| 86-TX-2-1.1 | NMSU0239 | TRUE | 28.918 | -99.795 | 23451 |
| 89-ILL-ILL-4.4 | NMSU0240 | TRUE | 39.301 | -90.595 | 15547 |
| 87-MX-2-2.2 | NMSU0241 | TRUE | 19.893 | -103.588 | 20570 |
| 87-MX-1-2.3 | NMSU0242 | TRUE | 21.672 | -99.539 | 21279 |
| 86-TX-1-4.2 | NMSU0243 | TRUE | 29.490 | -97.466 | 22643 |
| 87-MX-5-5.1 | NMSU0244 | TRUE | 23.367 | -99.475 | 22238 |
| 87-MX-1-1.5 | NMSU0245 | TRUE | 21.667 | -99.539 | 22317 |
| 86-TX-5-2.2 | NMSU0246 | TRUE | 31.354 | -100.209 | 24067 |
| 87-MX-1-4.2 | NMSU0247 | TRUE | 21.678 | -99.543 | 23522 |

| | | | | | |
|---|---|---|---|---|---|
| 87-MX-5-5.2 | NMSU0249 | TRUE | 23.367 | -99.475 | 20428 |
| 94-MX-320-5 | NMSU0250 | TRUE | 25.016 | -100.074 | 20951 |
| 94-MX-033-7 | NMSU0251 | TRUE | 26.826 | -102.255 | 22481 |
| 86-IL-1-3.2 | NMSU0252 | TRUE | 39.086 | -90.616 | 21530 |
| 87-MX-5-1.1 | NMSU0253 | TRUE | 23.405 | -99.381 | 21087 |
| 87MX-5-3.2 | NMSU0254 | TRUE | 23.404 | -99.382 | 22246 |
| 94-MX-329-9 | NMSU0255 | TRUE | 25.812 | -100.599 | 22605 |
| 94-MX-052-3 | NMSU0256 | TRUE | 27.311 | -102.397 | 21237 |
| 86-KS-2-4.2 | NMSU0257 | TRUE | 37.062 | -95.060 | 19866 |
| 87-MX-5-1.2 | NMSU0258 | TRUE | 23.405 | -99.381 | 22707 |
| 86-TX-3-1.2 | NMSU0259 | TRUE | 29.230 | -100.482 | 22350 |
| 94-MX-012-3 | NMSU0262 | TRUE | 26.826 | -102.255 | 22339 |
| 86-TX-1-3.2 | NMSU0263 | TRUE | 29.486 | -97.466 | 20151 |
| 87-MX-2-1.2 | NMSU0264 | TRUE | 19.891 | -103.586 | 21302 |
| 86-TX-5-1.2 | NMSU0266 | TRUE | 31.352 | -100.207 | 23975 |
| 87-MX-1-2.4 | NMSU0268 | TRUE | 21.672 | -99.539 | 21816 |
| 86-TX-2-2.4 | NMSU0270 | TRUE | 28.910 | -99.781 | 20291 |
| 87-MX-5-5.3 | NMSU0271 | TRUE | 23.367 | -99.475 | 23827 |
| 87-MX-5-1.3 | NMSU0272 | TRUE | 23.405 | -99.381 | 18849 |
| 87MX-5-3.3 | NMSU0273 | TRUE | 23.404 | -99.382 | 22517 |
| 86-TX-2-1.2 | NMSU0274 | TRUE | 28.918 | -99.795 | 22910 |
| 86-KS-2-4.3 | NMSU0275 | TRUE | 37.062 | -95.060 | 24033 |
| 86-TX-5-5.3 | NMSU0277 | TRUE | 31.481 | -100.467 | 23267 |
| 87-MX-5-1.4 | NMSU0279 | TRUE | 23.405 | -99.381 | 18777 |
| 87-MX-4-5.3 | NMSU0280 | TRUE | 20.486 | -99.221 | 23423 |
| 86-TX-1-4.4 | NMSU0281 | TRUE | 29.490 | -97.466 | 23299 |
| 86-TX-5-1.3 | NMSU0282 | TRUE | 31.352 | -100.207 | 21708 |
| 86-MO-1-1.2 | NMSU0283 | TRUE | 39.758 | -93.608 | 23724 |
| 86-MO-1-1.3 | NMSU0285 | TRUE | 39.758 | -93.608 | 21440 |
| 86-IL-1-3.4 | NMSU0286 | TRUE | 39.086 | -90.616 | 14677 |
| 87-MX-3-2.1 | NMSU0287 | TRUE | 16.951 | -96.756 | 22888 |

| | | | | | |
|---|---|---|---|---|---|
| 87-MX-4-3.3 | NMSU0288 | TRUE | 20.485 | -99.220 | 22381 |
| 86-TX-1-5.2 | NMSU0289 | TRUE | 29.495 | -97.460 | 18867 |
| 89-ILL-ALA-4.1 | NMSU0290 | TRUE | 32.497 | -87.716 | 17715 |
| 87-MX-5-4.4 | NMSU0291 | TRUE | 23.368 | -99.479 | 22149 |
| 87-MX-4-2.3 | NMSU0292 | TRUE | 20.484 | -99.222 | 23933 |
| 87-MX-1-2.5 | NMSU0293 | TRUE | 21.672 | -99.539 | 20702 |
| 86-TX-5-2.3 | NMSU0294 | TRUE | 31.354 | -100.209 | 23010 |
| 87-MX-5-5.4 | NMSU0295 | TRUE | 23.367 | -99.475 | 22776 |
| 87MX-5-3.4 | NMSU0298 | TRUE | 23.404 | -99.382 | 23306 |
| 89-ILL-UV-4.9 | NMSU0299 | TRUE | 29.530 | -100.005 | 22595 |
| 86-TX-4-5.3 | NMSU0300 | TRUE | 30.025 | -101.169 | 15220 |
| 87-MX-1-1.6 | NMSU0301 | TRUE | 21.667 | -99.539 | 19765 |
| 86-TX-5-4.3 | NMSU0302 | TRUE | 31.482 | -100.471 | 20948 |
| 87-MX-3-2.2 | NMSU0303 | TRUE | 16.951 | -96.756 | 23195 |
| 86-TX-5-5.4 | NMSU0304 | TRUE | 31.481 | -100.467 | 22129 |
| 86-TX-1-3.4 | NMSU0305 | TRUE | 29.486 | -97.466 | 21253 |
| 87-MX-2-1.3 | NMSU0306 | TRUE | 19.891 | -103.586 | 21474 |
| 86-MO-1-5.3 | NMSU0309 | TRUE | 39.701 | -93.305 | 16109 |
| 86-TX-5-3.3 | NMSU0312 | TRUE | 31.357 | -100.209 | 21032 |
| 86-TX-5-4.4 | NMSU0315 | TRUE | 31.482 | -100.471 | 20300 |
| 86-MO-1-1.4 | NMSU0316 | TRUE | 39.758 | -93.608 | 24413 |
| 87-MX-4-3.4 | NMSU0317 | TRUE | 20.485 | -99.220 | 18162 |
| 86-TX-5-1.4 | NMSU0318 | TRUE | 31.352 | -100.207 | 20318 |
| 86-TX-4-5.4 | NMSU0319 | TRUE | 30.025 | -101.169 | 14711 |
| 86-TX-5-2.6 | NMSU0322 | TRUE | 31.354 | -100.209 | 14349 |
| 86-KS-2-4.4 | NMSU0325 | TRUE | 37.062 | -95.060 | 18941 |
| 89-ILL-OK-5.3 | NMSU0326 | TRUE | 34.361 | -96.635 | 13990 |
| 87-MX-5-1.5 | NMSU0327 | TRUE | 23.405 | -99.381 | 19103 |
| 86-TX-5-5.6 | NMSU0328 | TRUE | 31.481 | -100.467 | 22712 |
| 86-TX-1-3.6 | NMSU0329 | TRUE | 29.486 | -97.466 | 21707 |
| 87-MX-1-1.7 | NMSU0330 | TRUE | 21.667 | -99.539 | 22157 |

| | | | | | |
|---|---|---|---|---|---|
| 86-TX-5-1.5 | NMSU0331 | TRUE | 31.352 | -100.207 | 17384 |
| 87-MX-4-2.4 | NMSU0335 | TRUE | 20.484 | -99.222 | 24053 |
| 86-TX-2-1.4 | NMSU0336 | TRUE | 28.918 | -99.795 | 14284 |
| 87-MX-1-2.6 | NMSU0337 | TRUE | 21.672 | -99.539 | 19271 |
| 87-MX-4-2.5 | NMSU0338 | TRUE | 20.484 | -99.222 | 21544 |
| 87-MX-5-1.6 | NMSU0339 | TRUE | 23.405 | -99.381 | 23354 |
| 86-TX-4-5.5 | NMSU0340 | TRUE | 30.025 | -101.169 | 14238 |
| 86-TX-2-1.5 | NMSU0341 | TRUE | 28.918 | -99.795 | 19888 |
| 86-TX-2-2.6 | NMSU0342 | TRUE | 28.910 | -99.781 | 20159 |
| 86-MO-1-1.5 | NMSU0343 | TRUE | 39.758 | -93.608 | 17025 |
| 86-TX-1-5.5 | NMSU0344 | TRUE | 29.495 | -97.460 | 14137 |
| 87-MX-5-4.6 | NMSU0345 | TRUE | 23.368 | -99.479 | 24219 |
| 87-MX-1-1.8 | NMSU0347 | TRUE | 21.667 | -99.539 | 18915 |
| 86-TX-5-4.6 | NMSU0348 | TRUE | 31.482 | -100.471 | 19641 |
| 86-MO-1-1.6 | NMSU0349 | TRUE | 39.758 | -93.608 | 19857 |
| 86-TX-4-5.6 | NMSU0350 | TRUE | 30.025 | -101.169 | 19908 |
| 87-MX-1-2.7 | NMSU0351 | TRUE | 21.672 | -99.539 | 20181 |
| 86-KS-2-5.5 | NMSU0352 | TRUE | 37.025 | -95.046 | 24174 |
| 87-MX-5-3.6 | NMSU0353 | TRUE | 23.404 | -99.382 | 21464 |
| 87-MX-4-3.6 | NMSU0355 | TRUE | 20.485 | -99.220 | 22262 |
| 86-IL-1-3.5 | NMSU0356 | TRUE | 39.086 | -90.616 | 24097 |
| 87MX-5-3.7 | NMSU0357 | TRUE | 23.404 | -99.382 | 18377 |
| 86-TX-3-1.6 | NMSU0358 | TRUE | 29.230 | -100.482 | 23093 |
| 87-MX-4-2.7 | NMSU0359 | TRUE | 20.484 | -99.222 | 22344 |
| 86-TX-2-1.6 | NMSU0360 | TRUE | 28.918 | -99.795 | 23838 |
| 86-TX-5-1.6 | NMSU0361 | TRUE | 31.352 | -100.207 | 24319 |
| 87-MX-3-2.4 | NMSU0362 | TRUE | 16.951 | -96.756 | 21809 |
| 86-TX-5-2.7 | NMSU0363 | TRUE | 31.354 | -100.209 | 22815 |
| 86-KS-2-4.5 | NMSU0364 | TRUE | 37.062 | -95.060 | 19309 |
| 86-TX-2-1.7 | NMSU0365 | TRUE | 28.918 | -99.795 | 18656 |
| 86-TX-1-5.6 | NMSU0368 | TRUE | 29.495 | -97.460 | 20072 |

| | | | | | |
|---|---|---|---|---|---|
| 87-MX-5-4.7 | NMSU0369 | TRUE | 23.368 | -99.479 | 21825 |
| 86-TX-5-5.7 | NMSU0370 | TRUE | 31.481 | -100.467 | 19017 |
| 89-ILL-ALA-4.2 | NMSU0371 | TRUE | 32.497 | -87.716 | 20228 |
| 87-MX-1-1.9 | NMSU0372 | TRUE | 21.667 | -99.539 | 14829 |
| 87-MX-4-5.5 | NMSU0373 | TRUE | 20.486 | -99.221 | 23244 |
| 87-MX-5-1.7 | NMSU0374 | TRUE | 23.405 | -99.381 | 17821 |
| 87-MX-1-1.10 | NMSU0375 | TRUE | 21.667 | -99.539 | 22744 |
| 86-TX-5-1.7 | NMSU0376 | TRUE | 31.352 | -100.207 | 20040 |
| 86-TX-5-5.8 | NMSU0377 | TRUE | 31.481 | -100.467 | 21782 |
| 86-MO-1-5.4 | NMSU0379 | TRUE | 39.701 | -93.305 | 17092 |
| 87-MX-5-5.5 | NMSU0380 | TRUE | 23.367 | -99.475 | 17763 |
| 87-MX-5-1.8 | NMSU0381 | TRUE | 23.405 | -99.381 | 16425 |
| 86-TX-1-5.7 | NMSU0382 | TRUE | 29.495 | -97.460 | 22532 |
| 86-TX-2-2.8 | NMSU0386 | TRUE | 28.910 | -99.781 | 23760 |
| 87-MX-5-5.6 | NMSU0387 | TRUE | 23.367 | -99.475 | 18937 |
| 86-TX-4-5.7 | NMSU0388 | TRUE | 30.025 | -101.169 | 21729 |
| 86-TX-5-2.8 | NMSU0389 | TRUE | 31.354 | -100.209 | 23324 |
| 87-MX-4-5.6 | NMSU0390 | TRUE | 20.486 | -99.221 | 14080 |
| 86-TX-1-4.8 | NMSU0391 | TRUE | 29.490 | -97.466 | 19264 |
| 86-TX-1-3.8 | NMSU0392 | TRUE | 29.486 | -97.466 | 17564 |
| 86-KS-2-4.6 | NMSU0393 | TRUE | 37.062 | -95.060 | 20551 |
| 86-TX-5-4.7 | NMSU0394 | TRUE | 31.482 | -100.471 | 22019 |
| Outpost | NMSU0395 | TRUE | 32.825 | -101.379 | 22584 |
| Duck Creek 25 | NMSU0396 | TRUE | 33.683 | -100.939 | 14950 |
| Duck Creek 16 | NMSU0397 | TRUE | 33.684 | -100.939 | 23934 |
| 86TX5-1 | NMSU0398 | TRUE | 31.352 | -100.207 | 23068 |
| Little Jewel | NMSU0400 | TRUE | 29.530 | -100.005 | 14035 |
| La Bahia | NMSU0402 | TRUE | 30.329 | -96.155 | 22365 |
| Anson Jones | NMSU0403 | TRUE | 30.322 | -96.147 | 18393 |
| Meier | NMSU0404 | FALSE | NA | NA | 20660 |
| 87MX3-2.11 | NMSU0405 | TRUE | 16.951 | -96.756 | 23249 |

| | | | | | |
|---|---|---|---|---|---|
| Fangue | NMSU0406 | TRUE | 29.945 | -96.120 | 21952 |
| Lane's Grandma | NMSU0407 | TRUE | 30.236 | -96.240 | 23794 |
| TN Big Tree | NMSU0408 | FALSE | NA | NA | 24139 |
| Weinman | NMSU0409 | TRUE | 29.888 | -96.104 | 23707 |
| 89-ILL-LA-3.1 | NMSU0410 | TRUE | 29.785 | -93.086 | 21734 |
| 89-ILL-LA-1.1 | NMSU0411 | TRUE | 29.784 | -93.087 | 23818 |
| 89-ILL-SBRA-2.3 | NMSU0412 | TRUE | 33.214 | -96.109 | 23618 |
| 89-ILL-LIPAN-3.1 | NMSU0415 | TRUE | 31.357 | -100.209 | 21964 |
| 89-ILL-CHAMP-2.1 | NMSU0416 | TRUE | 32.308 | -100.801 | 21832 |
| 89-ILL-OK-1.3 | NMSU0417 | TRUE | 34.090 | -96.960 | 20238 |
| 89-ILL-CON-3.4 | NMSU0418 | FALSE | NA | NA | 22981 |
| 89-ILL-IL-4.1 | NMSU0419 | TRUE | 39.301 | -90.595 | 21027 |
| 89-ILL-CON-6.1 | NMSU0420 | FALSE | NA | NA | 23726 |
| 89-ILL-LIPAN-1.1 | NMSU0421 | TRUE | 31.357 | -100.209 | 22133 |
| 89-ILL-ED-6.1 | NMSU0422 | TRUE | 29.792 | -100.415 | 23744 |
| 89-ILL-LMO-4.1 | NMSU0423 | TRUE | 29.290 | -100.424 | 22302 |
| 89-ILL-LA-3.2 | NMSU0424 | TRUE | 29.785 | -93.086 | 18492 |
| 89-ILL-SFE-3.1 | NMSU0425 | TRUE | 29.361 | -100.890 | 19593 |
| 89-ILL-CON-6.2 | NMSU0426 | FALSE | NA | NA | 19734 |
| 89-ILL-ALA-3.1 | NMSU0427 | TRUE | 32.497 | -87.716 | 23793 |
| 89-ILL-CON-2.1 | NMSU0428 | FALSE | NA | NA | 22839 |
| 89-ILL-SFE-3.2 | NMSU0429 | TRUE | 29.361 | -100.890 | 24494 |
| 89-ILL-DEV-1.6 | NMSU0430 | TRUE | 29.967 | -101.149 | 21125 |
| 89-ILL-SBRA-4.1 | NMSU0431 | TRUE | 32.746 | -96.101 | 16763 |
| 94-MX-064-8 | NMSU0432 | TRUE | 28.221 | -100.724 | 19835 |
| 89-ILL-LMO-2.1 | NMSU0433 | TRUE | 29.290 | -100.425 | 21557 |
| 89-ILL-DEV-3.1 | NMSU0434 | TRUE | 29.981 | -101.168 | 19101 |
| 89-ILL-SBRA-4.2 | NMSU0435 | TRUE | 32.746 | -96.101 | 18364 |
| 89-ILL-OK-5.1 | NMSU0436 | TRUE | 34.361 | -96.635 | 22318 |
| 89-ILL-SBRA-2.4 | NMSU0437 | TRUE | 33.214 | -96.109 | 21087 |
| 89-ILL-LMO-2.2 | NMSU0438 | TRUE | 29.290 | -100.425 | 23745 |

| | | | | | |
|---|---|---|---|---|---|
| 89-ILL-DEV-1.2 | NMSU0439 | TRUE | 29.967 | -101.149 | 23266 |
| 89-ILL-SFE-1.1 | NMSU0440 | TRUE | 29.371 | -100.884 | 24213 |
| Elliott | NMSU0709 | FALSE | NA | NA | 22876 |
| Riverside | NMSU0711 | FALSE | NA | NA | 20360 |
| Mahan | NMSU0732 | FALSE | NA | NA | 22409 |
| Spence | NMSU0744 | FALSE | NA | NA | 20988 |
| Elgin | NMSU0781 | FALSE | NA | NA | 18966 |
| Wichita | NMSU0837 | FALSE | NA | NA | 24672 |
| Curtis | NMSU0849 | FALSE | NA | NA | 24143 |
| Silverback | NMSU0850 | FALSE | NA | NA | 24073 |
| Giles | NMSU0852 | TRUE | 37.062 | -95.060 | 23879 |
| Jenkins | NMSU0854 | FALSE | NA | NA | 24107 |
| RuCox | NMSU0858 | FALSE | NA | NA | 16894 |
| Branch | NMSU0859 | FALSE | NA | NA | 23018 |
| Forey | NMSU0860 | FALSE | NA | NA | 20221 |
| Davis | NMSU0861 | FALSE | NA | NA | 13345 |
| Deakle's Special | NMSU0862 | FALSE | NA | NA | 23478 |
| Carman | NMSU0864 | FALSE | NA | NA | 21665 |
| Syrup Mill | NMSU0865 | FALSE | NA | NA | 23491 |
| Woodroof | NMSU0866 | FALSE | NA | NA | 14019 |
| Odom | NMSU0868 | FALSE | NA | NA | 21897 |
| Magenta | NMSU0869 | TRUE | 32.519 | -91.966 | 21481 |
| Carole Leigh | NMSU0870 | FALSE | NA | NA | 23144 |
| Mobile | NMSU0872 | FALSE | NA | NA | 23714 |
| Esnuel | NMSU0873 | FALSE | NA | NA | 23607 |
| Melrose | NMSU0874 | TRUE | 31.961 | -93.348 | 19471 |
| Koko | NMSU0875 | FALSE | NA | NA | 22605 |
| Iago | NMSU0877 | FALSE | NA | NA | 22630 |
| Sumner | NMSU0878 | FALSE | NA | NA | 23444 |
| El Mart | NMSU0879 | FALSE | NA | NA | 24201 |
| Kernodle | NMSU0880 | FALSE | NA | NA | 20268 |

| | | | | | |
|---|---|---|---|---|---|
| Woodard | NMSU0881 | FALSE | NA | NA | 23752 |
| Harris Super | NMSU0882 | FALSE | NA | NA | 23209 |
| Russell | NMSU0884 | FALSE | NA | NA | 23177 |
| UC7-15 | NMSU0885 | FALSE | NA | NA | 21821 |
| 99-ILL-SC-2 | NMSU0886 | FALSE | NA | NA | 23249 |
| Hughes | NMSU0887 | FALSE | NA | NA | 23853 |
| Broyles Jackson | NMSU0888 | FALSE | NA | NA | 23771 |
| Guenther | NMSU0889 | FALSE | NA | NA | 24142 |
| Starking Hardy Giant | NMSU0890 | TRUE | 39.371 | -93.065 | 22696 |
| Canton | NMSU0891 | TRUE | 40.132 | -91.530 | 24232 |
| Faith | NMSU0893 | FALSE | NA | NA | 18742 |
| Hodge | NMSU0894 | TRUE | 39.174 | -87.635 | 21792 |
| Peruque | NMSU0895 | TRUE | 38.879 | -90.626 | 22254 |
| Chief | NMSU0896 | TRUE | 37.806 | -88.262 | 23359 |
| Donald Grotian | NMSU0897 | TRUE | 39.423 | -93.152 | 22677 |
| MO-AES-2 | NMSU0899 | TRUE | 38.959 | -92.343 | 23868 |
| Hirschi | NMSU0900 | TRUE | 38.064 | -94.238 | 22692 |
| NC-4 | NMSU0901 | FALSE | NA | NA | 22240 |
| Dooley | NMSU0903 | TRUE | 35.622 | -96.014 | 23558 |
| Flack | NMSU0906 | FALSE | NA | NA | 20297 |
| Kibler | NMSU0907 | FALSE | NA | NA | 20080 |
| Warren 346 | NMSU0908 | TRUE | 39.779 | -93.370 | 19375 |
| Shepherd | NMSU0909 | TRUE | 39.158 | -94.496 | 20375 |
| Wiese | NMSU0910 | TRUE | 39.384 | -93.221 | 20681 |
| Goosepond | NMSU0911 | TRUE | 39.326 | -92.959 | 23554 |
| Busseron | NMSU0913 | TRUE | 38.835 | -87.449 | 20211 |
| GraZona | NMSU0914 | FALSE | NA | NA | 14919 |
| McCulley | NMSU0915 | TRUE | 31.723 | -98.958 | 20302 |
| Maramec | NMSU0916 | FALSE | NA | NA | 22699 |
| GraCross | NMSU0917 | FALSE | NA | NA | 19718 |
| Reece | NMSU0919 | FALSE | NA | NA | 17930 |

| | | | | | |
|---|---|---|---|---|---|
| RHS Special 1 | NMSU0921 | FALSE | NA | NA | 23615 |
| RHS Special 2 | NMSU0922 | FALSE | NA | NA | 22827 |
| Western | NMSU0923 | FALSE | NA | NA | 18156 |
| Kincaid | NMSU0924 | TRUE | 31.148 | -98.806 | 22206 |
| Carden | NMSU0925 | TRUE | 31.713 | -96.163 | 22793 |
| Brown's Leaning Tree | NMSU0927 | TRUE | 29.531 | -100.006 | 22336 |
| E Z Peel | NMSU0929 | FALSE | NA | NA | 21180 |
| Fifth Row | NMSU0930 | FALSE | NA | NA | 22996 |
| Fritz (TX) | NMSU0931 | FALSE | NA | NA | 22965 |
| Salado | NMSU0932 | FALSE | NA | NA | 24499 |
| F.W.Anderson | NMSU0933 | FALSE | NA | NA | 20873 |
| Salopek | NMSU0934 | FALSE | NA | NA | 19856 |
| Ames 25281.1 | NMSU0935 | FALSE | NA | NA | 23198 |
| Bryce | NMSU0936 | TRUE | 39.756 | -93.560 | 20341 |
| Biggs | NMSU0938 | FALSE | NA | NA | 22847 |
| Dumbell Lake Small | NMSU0940 | TRUE | 40.862 | -91.072 | 20607 |
| MX94-069-3 | NMSU0060 | TRUE | 28.409 | -100.889 | 18138 |
| MX94-010-8 | NMSU0062 | TRUE | 26.826 | -102.255 | 19785 |
| MX94-038-6 | NMSU0093 | TRUE | 26.826 | -102.255 | 20294 |
| MX94-240-6 | NMSU0094 | TRUE | 28.201 | -105.472 | 14712 |
| MX94-324-3 | NMSU0098 | TRUE | 26.530 | -100.502 | 21078 |
| MX94-332-2 | NMSU0100 | TRUE | 26.837 | -100.667 | 20310 |
| MX94-274-3 | NMSU0101 | TRUE | 25.540 | -100.943 | 23158 |
| MX94-310-7 | NMSU0104 | TRUE | 25.441 | -102.175 | 21200 |
| MX94-282-4 | NMSU0106 | TRUE | 25.441 | -102.175 | 20267 |
| MX94-036-6 | NMSU0115 | TRUE | 26.826 | -102.255 | 21988 |
| MX94-283-7 | NMSU0116 | TRUE | 21.053 | -99.815 | 22478 |
| MX94-331-2 | NMSU0120 | TRUE | 24.608 | -100.490 | 20555 |
| MX94-015-5 | NMSU0121 | TRUE | 28.492 | -100.920 | 22800 |
| MX94-082-6 | NMSU0124 | TRUE | 25.377 | -101.477 | 21291 |
| MX94-085-6 | NMSU0129 | TRUE | 25.441 | -102.175 | 22668 |

| | | | | | |
|---|---|---|---|---|---|
| MX94-051-9 | NMSU0131 | TRUE | 28.221 | -100.724 | 20283 |
| MX94-096-5 | NMSU0133 | TRUE | 25.441 | -102.175 | 20109 |
| MX94-112-4 | NMSU0139 | TRUE | 23.411 | -99.379 | 22080 |
| MX94-262-7 | NMSU0154 | TRUE | 26.530 | -100.502 | 22029 |
| MX94-244-5 | NMSU0181 | TRUE | 27.052 | -101.796 | 20233 |
| 86-TX-1-3.3 | NMSU0267 | TRUE | 29.486 | -97.466 | 22088 |
| 02-MYR-LA-LS2 | NMSU0307 | TRUE | 32.539 | -93.850 | 21621 |
| 86-TX-1-4.6 | NMSU0346 | TRUE | 29.490 | -97.466 | 22081 |
| 87-MX-4-2.6 | NMSU0354 | TRUE | 20.484 | -99.222 | 20359 |
| 86-TX-3-1.7 | NMSU0366 | TRUE | 29.230 | -100.482 | 19193 |
| 86-TX-2-2.7 | NMSU0378 | TRUE | 28.910 | -99.781 | 20473 |
| 86-MO-1-5.5 | NMSU0384 | TRUE | 39.701 | -93.305 | 22049 |
| 89-ILL-UV-3.4 | NMSU0441 | TRUE | 29.531 | -100.006 | 21787 |
| 89-ILL-SFE-1.2 | NMSU0442 | TRUE | 29.371 | -100.884 | 16430 |
| 89-ILL-UV-4.4 | NMSU0443 | TRUE | 29.530 | -100.005 | 23560 |
| 89-ILL-CONCHO-2 | NMSU0445 | TRUE | 31.481 | -100.467 | 23141 |
| 89-ILL-LIPAN-1.2 | NMSU0446 | TRUE | 31.357 | -100.209 | 23590 |
| 89-ILL-OK-5.2 | NMSU0447 | TRUE | 34.361 | -96.635 | 19686 |
| 89-ILL-STATON-1.1 | NMSU0448 | TRUE | 32.570 | -100.905 | 22913 |
| 87-MX-2-2.3 | NMSU0449 | TRUE | 19.893 | -103.588 | 21640 |
| 89-ILL-UV-4.3 | NMSU0450 | TRUE | 29.530 | -100.005 | 23821 |
| 89-ILL-LA-1.4 | NMSU0451 | TRUE | 29.784 | -93.087 | 20882 |
| 89-ILL-ALA-3.2 | NMSU0452 | TRUE | 32.497 | -87.716 | 23396 |
| 89-ILL-CON-2.2 | NMSU0453 | FALSE | NA | NA | 21606 |
| 89-ILL-CONCH-4 | NMSU0454 | TRUE | 31.482 | -100.471 | 20659 |
| 89-ILL-STATON-1.2 | NMSU0455 | TRUE | 32.570 | -100.905 | 21223 |
| 89-ILL-LIPAN-3.2 | NMSU0456 | TRUE | 31.357 | -100.209 | 21669 |
| 89-ILL-ED-1.1 | NMSU0457 | TRUE | 29.842 | -100.160 | 23335 |
| 89-ILL-ED-1.2 | NMSU0458 | TRUE | 29.842 | -100.160 | 22176 |
| 89-ILL-CHAMP-1.1 | NMSU0459 | TRUE | 32.309 | -100.800 | 21693 |
| 89-ILL-ED-6.2 | NMSU0461 | TRUE | 29.792 | -100.415 | 22628 |

| | | | | | |
|---|---|---|---|---|---|
| 89-ILL-CONCHO-4 | NMSU0463 | TRUE | 31.482 | -100.471 | 20137 |
| 94-MX-044-9 | NMSU0465 | TRUE | 28.221 | -100.724 | 23316 |
| 89-ILL-CHAMP-2.2 | NMSU0466 | TRUE | 32.308 | -100.801 | 21774 |
| 89-ILL-ILL-6.2 | NMSU0467 | TRUE | 41.978 | -90.552 | 19086 |
| 89-ILL-STATON-2.2 | NMSU0468 | TRUE | 32.573 | -100.907 | 21245 |
| 89-ILL-CONCH-2.1 | NMSU0469 | TRUE | 31.481 | -100.467 | 22487 |
| 89-ILL-CHAMP-1.2 | NMSU0471 | TRUE | 32.309 | -100.800 | 21395 |
| 89-ILL-HAYES-3.1 | NMSU0472 | TRUE | 32.796 | -101.006 | 20801 |
| 89-ILL-LA-1.5 | NMSU0474 | TRUE | 29.784 | -93.087 | 21194 |
| 89-ILL-ALA-3.3 | NMSU0475 | TRUE | 32.497 | -87.716 | 20143 |
| 89-ILL-UV-4.1 | NMSU0476 | TRUE | 29.530 | -100.005 | 22153 |
| 89-ILL-SFE-1.3 | NMSU0477 | TRUE | 29.371 | -100.884 | 21635 |
| 89-ILL-HAYES-1.3 | NMSU0478 | TRUE | 32.797 | -101.010 | 21750 |
| 89-ILL-UV-3.1 | NMSU0479 | TRUE | 29.531 | -100.006 | 21378 |
| 89-ILL-OK-5.4 | NMSU0480 | TRUE | 34.361 | -96.635 | 21666 |
| 89-ILL-HAYES-1.4 | NMSU0481 | TRUE | 32.797 | -101.010 | 20647 |
| 89-ILL-SCHAMP-1.1 | NMSU0482 | TRUE | 32.309 | -100.800 | 22804 |
| 89-ILL-CON-6.3 | NMSU0483 | FALSE | NA | NA | 20503 |
| 89-ILL-LA-3.3 | NMSU0484 | TRUE | 29.785 | -93.086 | 22197 |
| 89-ILL-ALA-3.4 | NMSU0485 | TRUE | 32.497 | -87.716 | 22862 |
| 89-ILL-OK-5.3 | NMSU0486 | TRUE | 34.361 | -96.635 | 22522 |
| 89-ILL-CON-3.1 | NMSU0487 | FALSE | NA | NA | 21938 |
| 89-ILL-CON-2.3 | NMSU0488 | FALSE | NA | NA | 18644 |
| 89-ILL-LA-3.4 | NMSU0489 | TRUE | 29.785 | -93.086 | 22650 |
| CSPA1-15 | NMSU0491 | FALSE | NA | NA | 21007 |
| 89-ILL-CHAMP-1.3 | NMSU0492 | TRUE | 32.309 | -100.800 | 16317 |
| 89-ILL-CON-3.5 | NMSU0493 | FALSE | NA | NA | 21099 |
| 89-ILL-CON-6.4 | NMSU0494 | FALSE | NA | NA | 21914 |
| 89-ILL-CON-2.4 | NMSU0495 | FALSE | NA | NA | 22211 |
| 89-ILL-HAYES-3.2 | NMSU0496 | TRUE | 32.796 | -101.006 | 21619 |
| 89-ILL-STATON-1.3 | NMSU0497 | TRUE | 32.570 | -100.905 | 20919 |

| | | | | | |
|---|---|---|---|---|---|
| 89-ILL-ILL-6.3 | NMSU0499 | TRUE | 41.978 | -90.552 | 21256 |
| 89-ILL-LA-1.2 | NMSU0500 | TRUE | 29.784 | -93.087 | 21140 |
| 89-ILL-STATON-2.4 | NMSU0501 | TRUE | 32.573 | -100.907 | 20795 |
| 89-ILL-SBRA-2.5 | NMSU0502 | TRUE | 33.214 | -96.109 | 22212 |
| 89-ILL-LIPAN-1.3 | NMSU0503 | TRUE | 31.357 | -100.209 | 22475 |
| 89-ILL-ED-6.4 | NMSU0504 | TRUE | 29.792 | -100.415 | 19553 |
| 89-ILL-DEV-1.7 | NMSU0506 | TRUE | 29.967 | -101.149 | 21938 |
| 89-ILL-ED-1.3 | NMSU0510 | TRUE | 29.842 | -100.160 | 22119 |
| 89-ILL-SBRA-2.1 | NMSU0511 | TRUE | 33.214 | -96.109 | 19917 |
| 89-ILL-SBRA-4.4 | NMSU0512 | TRUE | 32.746 | -96.101 | 20231 |
| 89-ILL-LMO-4.3 | NMSU0513 | TRUE | 29.290 | -100.424 | 21749 |
| 89-ILL-DEV-3.3 | NMSU0514 | TRUE | 29.981 | -101.168 | 19057 |
| 89-ILL-SFE-1.4 | NMSU0515 | TRUE | 29.371 | -100.884 | 21636 |
| 89-ILL-UV-3.6 | NMSU0516 | TRUE | 29.531 | -100.006 | 18230 |
| 89-ILL-ED-6.3 | NMSU0517 | TRUE | 29.792 | -100.415 | 18627 |
| 89-ILL-CONCH-2.2 | NMSU0518 | TRUE | 31.481 | -100.467 | 21339 |
| 89-ILL-LIPAN-3.3 | NMSU0521 | TRUE | 31.357 | -100.209 | 19142 |
| 89-ILL-ILL-6.4 | NMSU0522 | TRUE | 41.978 | -90.552 | 19755 |
| 89-ILL-LIPAN-1.4 | NMSU0524 | TRUE | 31.357 | -100.209 | 19943 |
| 89-ILL-LMO-2.3 | NMSU0526 | TRUE | 29.290 | -100.425 | 22081 |
| 89-ILL-CHAMP-2.4 | NMSU0530 | TRUE | 32.308 | -100.801 | 20373 |
| 89-ILL-CON-6.5 | NMSU0531 | FALSE | NA | NA | 19806 |
| 89-ILL-UV-3.7 | NMSU0532 | TRUE | 29.531 | -100.006 | 23018 |
| 89-ILL-LA-1.6 | NMSU0533 | TRUE | 29.784 | -93.087 | 21608 |
| 89-ILL-STATON-2.5 | NMSU0534 | TRUE | 32.573 | -100.907 | 23950 |
| 89-ILL-CON-2.5 | NMSU0535 | FALSE | NA | NA | 17529 |
| 89-ILL-LA-3.5 | NMSU0536 | TRUE | 29.785 | -93.086 | 22778 |
| 89-ILL-DEV-3.4 | NMSU0537 | TRUE | 29.981 | -101.168 | 21779 |
| 89-ILL-CON-3.2 | NMSU0538 | FALSE | NA | NA | 22300 |
| 89-ILL-DEV-1.9 | NMSU0542 | TRUE | 29.967 | -101.149 | 21860 |
| 89-ILL-SFE-1.6 | NMSU0543 | TRUE | 29.371 | -100.884 | 22587 |

| | | | | | |
|---|---|---|---|---|---|
| 89-ILL-CONCHO-2 | NMSU0544 | TRUE | 31.481 | -100.467 | 22405 |
| 89-ILL-ED-6.5 | NMSU0545 | TRUE | 29.792 | -100.415 | 23589 |
| 89-ILL-CON-3.6 | NMSU0546 | FALSE | NA | NA | 20194 |
| 89-ILL-CONCHO-4 | NMSU0547 | TRUE | 31.482 | -100.471 | 20192 |
| 89-ILL-HAYES-3.4 | NMSU0548 | TRUE | 32.796 | -101.006 | 20007 |
| CSPA18-10 | NMSU0553 | FALSE | NA | NA | 22592 |
| 89-ILL-ALA-3.5 | NMSU0555 | TRUE | 32.497 | -87.716 | 22852 |
| 89-ILL-SFE-3.5 | NMSU0556 | TRUE | 29.361 | -100.890 | 12939 |
| 89-ILL-LMO-4.5 | NMSU0558 | TRUE | 29.290 | -100.424 | 21324 |
| 89-ILL-LIPAN-1.5 | NMSU0559 | TRUE | 31.357 | -100.209 | 22300 |
| 89-ILL-LIPAN-3.4 | NMSU0560 | TRUE | 31.357 | -100.209 | 22560 |
| 89-ILL-CHAMP-1.4 | NMSU0561 | TRUE | 32.309 | -100.800 | 15907 |
| 89-ILL-ED-1.6 | NMSU0562 | TRUE | 29.842 | -100.160 | 22506 |
| CSPA18-13 | NMSU0563 | FALSE | NA | NA | 14888 |
| 89-ILL-LIPAN-3.5 | NMSU0564 | TRUE | 31.357 | -100.209 | 22155 |
| 89-ILL-ED-6.6 | NMSU0565 | TRUE | 29.792 | -100.415 | 21749 |
| 89-ILL-CHAMP-1.5 | NMSU0567 | TRUE | 32.309 | -100.800 | 21362 |
| 89-ILL-ED-1.5 | NMSU0568 | TRUE | 29.842 | -100.160 | 22759 |
| CSPA18-15 | NMSU0569 | FALSE | NA | NA | 15285 |
| 89-ILL-HAYES-1.5 | NMSU0571 | TRUE | 32.797 | -101.010 | 22874 |
| 89-ILL-ALA-3.6 | NMSU0572 | TRUE | 32.497 | -87.716 | 21465 |
| 89-ILL-OK-1.6 | NMSU0573 | TRUE | 34.090 | -96.960 | 21307 |
| 89-ILL-SBRA-2.6 | NMSU0574 | TRUE | 33.214 | -96.109 | 21203 |
| 89-ILL-SBRA-6.1 | NMSU0575 | FALSE | NA | NA | 18358 |
| 89-ILL-STATON-1.4 | NMSU0576 | TRUE | 32.570 | -100.905 | 22573 |
| 89-ILL-UV-4.6 | NMSU0577 | TRUE | 29.530 | -100.005 | 21565 |
| 89-ILL-DEV-1.10 | NMSU0578 | TRUE | 29.967 | -101.149 | 20981 |
| 89-ILL-OK-1.2 | NMSU0579 | TRUE | 34.090 | -96.960 | 21357 |
| 89-ILL-HAYES-3.5 | NMSU0580 | TRUE | 32.796 | -101.006 | 16561 |
| 89-ILL-STATON-1.5 | NMSU0581 | TRUE | 32.570 | -100.905 | 20035 |
| 89-ILL-CON-3.7 | NMSU0582 | FALSE | NA | NA | 21373 |

| | | | | | |
|---|---|---|---|---|---|
| 89-ILL-LMO-4.6 | NMSU0583 | TRUE | 29.290 | -100.424 | 20153 |
| 89-ILL-LA-1.7 | NMSU0584 | TRUE | 29.784 | -93.087 | 23061 |
| 89-ILL-IL-4.2 | NMSU0585 | TRUE | 39.301 | -90.595 | 21184 |
| 89-ILL-CONCHO-2 | NMSU0587 | TRUE | 31.481 | -100.467 | 22997 |
| 89-ILL-STATON-2.6 | NMSU0589 | TRUE | 32.573 | -100.907 | 20099 |
| 89-ILL-LA-3.6 | NMSU0590 | TRUE | 29.785 | -93.086 | 21160 |
| 89-ILL-UV-3.3 | NMSU0591 | TRUE | 29.531 | -100.006 | 17625 |
| 89-ILL-HAYES-1.6 | NMSU0592 | TRUE | 32.797 | -101.010 | 19619 |
| 89-ILL-SFE-1.7 | NMSU0593 | TRUE | 29.371 | -100.884 | 21516 |
| 89-ILL-ALA-3.7 | NMSU0595 | TRUE | 32.497 | -87.716 | 21485 |
| 89-ILL-CON-2.7 | NMSU0596 | FALSE | NA | NA | 23844 |
| 89-ILL-SCHAMP-2.1 | NMSU0597 | TRUE | 32.309 | -100.800 | 21608 |
| 89-ILL-ED-6.8 | NMSU0598 | TRUE | 29.792 | -100.415 | 18288 |
| 89-ILL-LA-3.7 | NMSU0599 | TRUE | 29.785 | -93.086 | 21040 |
| 89-ILL-HAYES-3.7 | NMSU0600 | TRUE | 32.796 | -101.006 | 20844 |
| 89-ILL-UV-3.8 | NMSU0601 | TRUE | 29.531 | -100.006 | 20643 |
| 89-ILL-SBRA-4.7 | NMSU0602 | TRUE | 32.746 | -96.101 | 19955 |
| 89-ILL-LMO-2.5 | NMSU0603 | TRUE | 29.290 | -100.425 | 20373 |
| 89-ILL-CHAMP-1.6 | NMSU0604 | TRUE | 32.309 | -100.800 | 22473 |
| 89-ILL-SBRA-4.3 | NMSU0605 | TRUE | 32.746 | -96.101 | 19979 |
| 89-ILL-ILL-6 | NMSU0606 | TRUE | 41.978 | -90.552 | 21044 |
| 89-ILL-SFE-3.6 | NMSU0607 | TRUE | 29.361 | -100.890 | 22261 |
| 89-ILL-CHAMP-4.1 | NMSU0609 | TRUE | 32.320 | -100.746 | 20686 |
| 89-ILL-CONCHO-4 | NMSU0610 | TRUE | 31.482 | -100.471 | 18953 |
| 89-ILL-CON-6.7 | NMSU0611 | FALSE | NA | NA | 22127 |
| 89-ILL-CON-6.8 | NMSU0612 | FALSE | NA | NA | 22343 |
| 89-ILL-SFE-3.7 | NMSU0613 | TRUE | 29.361 | -100.890 | 19218 |
| 89-ILL-CONCH-4 | NMSU0614 | TRUE | 31.482 | -100.471 | 20370 |
| 89-ILL-UV-4.8 | NMSU0615 | TRUE | 29.530 | -100.005 | 21429 |
| 89-ILL-LIPAN-1.7 | NMSU0616 | TRUE | 31.357 | -100.209 | 23186 |
| 89-ILL-OK-5.8 | NMSU0617 | TRUE | 34.361 | -96.635 | 19815 |

| | | | | | |
|---|---|---|---|---|---|
| 89-ILL-SFE-1.8 | NMSU0618 | TRUE | 29.371 | -100.884 | 22748 |
| 89-ILL-HAYES-1.7 | NMSU0619 | TRUE | 32.797 | -101.010 | 18568 |
| 89-ILL-CONCH-2.5 | NMSU0620 | TRUE | 31.481 | -100.467 | 20405 |
| 89-ILL-LMO-4.7 | NMSU0621 | TRUE | 29.290 | -100.424 | 15199 |
| 89-ILL-UV-4.7 | NMSU0622 | TRUE | 29.530 | -100.005 | 18153 |
| 89-ILL-DEV-1.11 | NMSU0623 | TRUE | 29.967 | -101.149 | 21863 |
| 89-ILL-OK-5.7 | NMSU0625 | TRUE | 34.361 | -96.635 | 18636 |
| 89-ILL-ED-1.7 | NMSU0626 | TRUE | 29.842 | -100.160 | 22870 |
| 89-ILL-OK-1.7 | NMSU0627 | TRUE | 34.090 | -96.960 | 23010 |
| 89-ILL-SBRA-2.7 | NMSU0628 | TRUE | 33.214 | -96.109 | 21596 |
| 89-ILL-STATON-1.6 | NMSU0629 | TRUE | 32.570 | -100.905 | 22004 |
| 89-ILL-CON-2.8 | NMSU0630 | FALSE | NA | NA | 21876 |
| 89-ILL-DEV-1.12 | NMSU0631 | TRUE | 29.967 | -101.149 | 20958 |
| 89-ILL-LIPAN-3.7 | NMSU0632 | TRUE | 31.357 | -100.209 | 19966 |
| 89-ILL-ALA-3.8 | NMSU0633 | TRUE | 32.497 | -87.716 | 22019 |
| 89-ILL-SBRA-2 | NMSU0634 | TRUE | 33.214 | -96.109 | 22756 |
| 89-ILL-LA-1.8 | NMSU0635 | TRUE | 29.784 | -93.087 | 19084 |
| 89-ILL-CON-3.8 | NMSU0637 | FALSE | NA | NA | 19742 |
| 89-ILL-STATON-2.7 | NMSU0638 | TRUE | 32.573 | -100.907 | 18968 |
| RDM-LL-10 | NMSU0639 | FALSE | NA | NA | 19238 |
| RDM-LL-15 | NMSU0640 | FALSE | NA | NA | 18447 |
| Silverback-14 | NMSU0641 | FALSE | NA | NA | 19973 |
| 94-ILL-ARK-1.3 | NMSU0642 | TRUE | 34.610 | -92.185 | 22664 |
| 94-ILL-RR-2.2 | NMSU0644 | TRUE | 33.571 | -94.207 | 21735 |
| 94-ILL-RR-4.3 | NMSU0645 | TRUE | 33.567 | -94.207 | 20294 |
| 94-ILL-STF-2.1 | NMSU0646 | TRUE | 35.522 | -90.432 | 19664 |
| 94-ILL-STF-5.4 | NMSU0647 | TRUE | 35.511 | -90.444 | 16926 |
| 94-ILL-ARK-4.6 | NMSU0648 | TRUE | 34.598 | -92.192 | 18955 |
| 94-ILL-ARK-2.4 | NMSU0649 | TRUE | 34.604 | -92.189 | 21021 |
| 94-ILL-STF-4.5 | NMSU0650 | TRUE | 35.511 | -90.446 | 23542 |
| 94-ILL-RR-5.6 | NMSU0651 | TRUE | 33.566 | -94.209 | 18833 |

| | | | | | |
|---|---|---|---|---|---|
| 94-ILL-STF-1.6 | NMSU0652 | TRUE | 35.522 | -90.431 | 20482 |
| 97-26-0010 | NMSU0653 | FALSE | NA | NA | 18928 |
| 94-ILL-RR-1.8 | NMSU0654 | TRUE | 33.573 | -94.203 | 21099 |
| 94-ILL-ARK-3.9 | NMSU0655 | TRUE | 34.599 | -92.192 | 19319 |
| 94-ILL-STF-3.8 | NMSU0657 | TRUE | 35.511 | -90.446 | 20507 |
| 94-ILL-RR-6.7 | NMSU0658 | TRUE | 33.569 | -94.204 | 20869 |
| 97-ILL-LA-1.12 | NMSU0660 | TRUE | 30.065 | -93.351 | 21600 |
| 97-ILL-LA-3.1 | NMSU0661 | TRUE | 30.065 | -93.351 | 21803 |
| 97-ILL-LA-WB1.14 | NMSU0662 | TRUE | 30.094 | -93.391 | 18689 |
| 97-ILL-LA-WB2.3 | NMSU0663 | TRUE | 30.094 | -93.392 | 18800 |
| 97-ILL-LA-WB3.1 | NMSU0664 | TRUE | 30.095 | -93.392 | 21220 |
| 97-ILL-CAT-1.1 | NMSU0665 | TRUE | 29.571 | -92.239 | 20192 |
| 97-ILL-CAT-7.17 | NMSU0666 | TRUE | 29.570 | -92.224 | 19493 |
| 97-ILL-CAT-8.5 | NMSU0667 | TRUE | 29.570 | -92.223 | 21550 |
| 97-ILL-CAT-9.4 | NMSU0668 | TRUE | 29.570 | -92.222 | 20978 |
| 97-ILL-CAT-11.3 | NMSU0669 | TRUE | 29.568 | -92.204 | 22193 |
| 97-ILL-CAT-13.9 | NMSU0671 | TRUE | 29.569 | -92.203 | 22574 |
| 97-ILL-CAT-14.13 | NMSU0672 | TRUE | 29.569 | -92.203 | 21533 |
| 97-ILL-CAT-15.15 | NMSU0673 | TRUE | 29.569 | -92.203 | 21077 |
| 97-ILL-CAT-16.2 | NMSU0674 | TRUE | 29.569 | -92.203 | 18715 |
| Womack Dwarf | NMSU0684 | FALSE | NA | NA | 12817 |
| Apache | NMSU0707 | FALSE | NA | NA | 24320 |
| Centennial | NMSU0708 | TRUE | 30.004 | -90.776 | 21413 |
| Moore | NMSU0710 | FALSE | NA | NA | 18469 |
| San Felipe 1 | NMSU0712 | TRUE | 29.371 | -100.884 | 20916 |
| VC1-68 | NMSU0713 | FALSE | NA | NA | 21415 |
| Oliver | NMSU0715 | TRUE | 30.495 | -99.732 | 22351 |
| Delmas | NMSU0716 | FALSE | NA | NA | 22622 |
| Altman | NMSU0717 | TRUE | 30.769 | -98.264 | 21753 |
| Rome | NMSU0718 | FALSE | NA | NA | 23295 |
| Carter | NMSU0719 | FALSE | NA | NA | 20564 |

| | | | | | |
|---|---|---|---|---|---|
| Desirable | NMSU0720 | FALSE | NA | NA | 18984 |
| Alley | NMSU0721 | FALSE | NA | NA | 22445 |
| Baker | NMSU0722 | FALSE | NA | NA | 20696 |
| Barton | NMSU0724 | FALSE | NA | NA | 22551 |
| James(LA) | NMSU0725 | FALSE | NA | NA | 21797 |
| Cooper | NMSU0726 | FALSE | NA | NA | 22268 |
| Brake | NMSU0727 | FALSE | NA | NA | 21754 |
| Jack Doby | NMSU0728 | FALSE | NA | NA | 19526 |
| Barton (Catwalk) | NMSU0729 | FALSE | NA | NA | 19398 |
| Ramsey Mediumshell | NMSU0730 | FALSE | NA | NA | 21181 |
| Farley | NMSU0731 | FALSE | NA | NA | 19929 |
| Nelson | NMSU0733 | FALSE | NA | NA | 23706 |
| UE 4-13 | NMSU0734 | FALSE | NA | NA | 20823 |
| Philema | NMSU0735 | FALSE | NA | NA | 23125 |
| Schley | NMSU0736 | FALSE | NA | NA | 22163 |
| UE 1-13 | NMSU0737 | FALSE | NA | NA | 20690 |
| Woodside Early | NMSU0738 | FALSE | NA | NA | 21074 |
| Cherryle | NMSU0739 | FALSE | NA | NA | 22678 |
| Mississippi 10 | NMSU0740 | FALSE | NA | NA | 23293 |
| Schaeffer | NMSU0741 | FALSE | NA | NA | 22021 |
| Moneymaker | NMSU0742 | FALSE | NA | NA | 23561 |
| Stuart | NMSU0743 | FALSE | NA | NA | 25044 |
| Surprize | NMSU0746 | FALSE | NA | NA | 24192 |
| Teche | NMSU0747 | FALSE | NA | NA | 22413 |
| Linberger | NMSU0748 | FALSE | NA | NA | 21891 |
| Moreland | NMSU0749 | FALSE | NA | NA | 23548 |
| Pointe Coupee 2 | NMSU0751 | TRUE | 30.734 | -91.433 | 21385 |
| Dependable | NMSU0752 | FALSE | NA | NA | 14830 |
| Mahan Stuart | NMSU0753 | FALSE | NA | NA | 19670 |
| Pensacola Cluster | NMSU0754 | FALSE | NA | NA | 23901 |
| Neuman | NMSU0755 | FALSE | NA | NA | 22939 |

| | | | | | |
|---|---|---|---|---|---|
| Jackson (LA) | NMSU0756 | FALSE | NA | NA | 19161 |
| Amling | NMSU0757 | FALSE | NA | NA | 20612 |
| A-93 | NMSU0758 | FALSE | NA | NA | 21685 |
| Hipps | NMSU0759 | FALSE | NA | NA | 22322 |
| 1950-12-0090 | NMSU0760 | FALSE | NA | NA | 24008 |
| Jubilee | NMSU0761 | FALSE | NA | NA | 22423 |
| Candy | NMSU0762 | FALSE | NA | NA | 23870 |
| Colby | NMSU0763 | TRUE | 38.679 | -89.309 | 23764 |
| Chetopa (K-112) | NMSU0764 | FALSE | NA | NA | 23945 |
| Frisbie | NMSU0765 | TRUE | 39.754 | -93.549 | 21644 |
| Tiny Tim | NMSU0766 | TRUE | 38.970 | -90.832 | 23271 |
| Posey | NMSU0767 | TRUE | 38.389 | -87.555 | 21563 |
| Mullahy | NMSU0768 | TRUE | 41.171 | -90.996 | 21822 |
| Spence (MO) | NMSU0770 | TRUE | 39.426 | -93.131 | 23796 |
| Witte (IA) | NMSU0771 | TRUE | 40.840 | -91.161 | 21275 |
| Weschcke Survivor | NMSU0772 | FALSE | NA | NA | 22223 |
| Aquilla | NMSU0773 | FALSE | NA | NA | 22758 |
| Lucas | NMSU0774 | TRUE | 40.705 | -82.418 | 22115 |
| Buchel 1 | NMSU0778 | TRUE | 29.098 | -97.285 | 21505 |
| Carmichael | NMSU0779 | TRUE | 29.655 | -97.368 | 22997 |
| Clark | NMSU0780 | TRUE | 31.105 | -98.505 | 21741 |
| Evans | NMSU0782 | TRUE | 29.867 | -96.817 | 22187 |
| Evers | NMSU0783 | TRUE | 33.206 | -97.080 | 20942 |
| Fayette | NMSU0785 | FALSE | NA | NA | 23475 |
| Meyers(TX) | NMSU0786 | TRUE | 30.209 | -97.592 | 20370 |
| Gloria Grande | NMSU0787 | FALSE | NA | NA | 21870 |
| N1-62 | NMSU0788 | FALSE | NA | NA | 24497 |
| Foster | NMSU0790 | TRUE | 29.461 | -97.658 | 23619 |
| Freeman | NMSU0791 | TRUE | 32.816 | -95.664 | 23319 |
| Gay | NMSU0792 | TRUE | 29.465 | -96.539 | 15891 |
| Govett | NMSU0794 | TRUE | 29.568 | -97.885 | 21915 |

| | | | | | |
|---|---|---|---|---|---|
| GraPark Giant | NMSU0795 | FALSE | NA | NA | 23533 |
| Halsly | NMSU0796 | FALSE | NA | NA | 24290 |
| Hensel | NMSU0797 | TRUE | 29.893 | -96.871 | 23354 |
| Hollis | NMSU0798 | TRUE | 31.100 | -98.514 | 22397 |
| Humble | NMSU0799 | TRUE | 29.087 | -99.875 | 23365 |
| Ideal | NMSU0800 | TRUE | 31.196 | -98.718 | 22201 |
| Jersey | NMSU0801 | TRUE | 31.250 | -98.596 | 23140 |
| Johnson(KS) | NMSU0802 | TRUE | 37.176 | -94.820 | 21318 |
| Jin Hua 1 | NMSU0803 | FALSE | NA | NA | 22697 |
| Coy | NMSU0804 | TRUE | 37.169 | -94.844 | 23887 |
| Late | NMSU0805 | TRUE | 31.374 | -98.677 | 20802 |
| Number 54 | NMSU0807 | FALSE | NA | NA | 24054 |
| Mount | NMSU0808 | TRUE | 35.584 | -95.992 | 20625 |
| Nugget | NMSU0809 | TRUE | 31.837 | -98.396 | 21127 |
| Die Guet | NMSU0810 | TRUE | 29.411 | -98.905 | 23262 |
| Oklahoma | NMSU0811 | TRUE | 34.239 | -97.083 | 23237 |
| GraKing | NMSU0812 | FALSE | NA | NA | 13350 |
| Osborne | NMSU0813 | TRUE | 30.821 | -98.579 | 23038 |
| Pearce | NMSU0814 | TRUE | 30.344 | -97.886 | 23317 |
| Ripe Early | NMSU0815 | TRUE | 35.520 | -97.240 | 22699 |
| Risien 1 | NMSU0816 | TRUE | 31.247 | -98.594 | 21541 |
| Roth | NMSU0817 | TRUE | 29.248 | -97.329 | 22756 |
| Schutz 1 | NMSU0818 | TRUE | 29.482 | -97.583 | 22191 |
| Schutz 2 | NMSU0819 | TRUE | 29.485 | -97.586 | 24228 |
| Squirrel's Delight | NMSU0820 | TRUE | 31.249 | -98.597 | 23969 |
| Tarrant | NMSU0822 | FALSE | NA | NA | 23356 |
| Texas 60 | NMSU0823 | TRUE | 31.250 | -98.596 | 24582 |
| Kiowa | NMSU0824 | FALSE | NA | NA | 20404 |
| Tiemann | NMSU0825 | TRUE | 29.896 | -96.865 | 22374 |
| Tissue Paper | NMSU0826 | TRUE | 37.164 | -94.843 | 22028 |
| Williamson | NMSU0827 | TRUE | 34.404 | -96.826 | 20667 |

| | | | | | |
|---|---|---|---|---|---|
| Winner | NMSU0828 | TRUE | 31.375 | -98.670 | 22269 |
| Barraza 3 | NMSU0830 | FALSE | NA | NA | 23986 |
| Brzozowski-Vlasek | NMSU0831 | TRUE | 29.522 | -97.491 | 23903 |
| Keilers (Stockbauer) | NMSU0832 | FALSE | NA | NA | 23696 |
| San Saba Improved | NMSU0834 | FALSE | NA | NA | 22651 |
| Prilop | NMSU0835 | TRUE | 29.424 | -96.940 | 22902 |
| Sioux | NMSU0836 | FALSE | NA | NA | 23720 |
| Hogan | NMSU0853 | FALSE | NA | NA | 20374 |
| Allbritton | NMSU0857 | TRUE | 32.120 | -93.520 | 22581 |
| Tinker | NMSU0871 | FALSE | NA | NA | 21599 |
| Greenriver | NMSU0898 | TRUE | 37.898 | -87.491 | 18644 |
| Best's Early | NMSU0902 | TRUE | 39.288 | -90.552 | 20122 |
| Carlson #3 | NMSU0904 | TRUE | 41.176 | -91.000 | 18312 |
| Shag | NMSU0905 | TRUE | 37.534 | -95.826 | 20954 |
| GraTex | NMSU0918 | FALSE | NA | NA | 21227 |
| Limpia Creek | NMSU0920 | TRUE | 30.640 | -103.846 | 20599 |
| Champ 15 | NMSU0928 | FALSE | NA | NA | 20811 |
| Duck Creek 1 | NMSU0937 | TRUE | 33.684 | -100.939 | 20930 |
| Green Island Beaver | NMSU0941 | TRUE | 42.177 | -90.301 | 15666 |
| Green Island Hackberry | NMSU0942 | TRUE | 42.182 | -90.299 | 17286 |
| Podsednik | NMSU0943 | FALSE | NA | NA | 18539 |
| Old Woman | NMSU0944 | TRUE | 40.862 | -91.073 | 21451 |
| Red Nut | NMSU0945 | FALSE | NA | NA | 19443 |
| Snodgrass | NMSU0946 | FALSE | NA | NA | 19680 |
| Warsaw No. 1 | NMSU0947 | TRUE | 40.359 | -91.435 | 16691 |
| Excel | NMSU0954 | FALSE | NA | NA | 21536 |
| Gafford | NMSU0955 | FALSE | NA | NA | 20682 |
| GRE 14-20 | NMSU0956 | FALSE | NA | NA | 20611 |
| Miss L | NMSU0957 | FALSE | NA | NA | 14271 |
| Martzahn | NMSU0958 | TRUE | 40.838 | -91.115 | 18453 |
| PA Big Tree | NMSU0959 | FALSE | NA | NA | 16981 |

| | | | | | |
|---|---|---|---|---|---|
| CSPB13-35 | NMSU0968 | FALSE | NA | NA | 21076 |
| Norton | NMSU0982 | TRUE | 39.313 | -90.820 | 21628 |
| Jefferson | NMSU0984 | FALSE | NA | NA | 16189 |
| RS11-1 | NMSU0985 | FALSE | NA | NA | 19747 |
| Pabst | NMSU0996 | FALSE | NA | NA | 17955 |
| 10-ILL-IA-4.1 | NMSU0999 | TRUE | 41.228 | -91.113 | 21764 |
| MX94-318-2 | NMSU0150 | FALSE | NA | NA | 15593 |
| 94-MX-046-1 | NMSU0260 | FALSE | NA | NA | 19638 |
| 87-MX-4-5.2 | NMSU0261 | FALSE | NA | NA | 18901 |

[1] Sample ID in VCF genotype files.
[2]Number of SNPs with genotypes for each sample.

**Table S5: Fst values between pecan gene pools**

| Gene Pool | Central Mexico | Northern Mexico 1 | Northern Mexico 2 | Texas and Mexico | Northern USA |
|---|---|---|---|---|---|
| Central Mexico | - | 0.119[1] | 0.065[1] | 0.087[1] | 0.089[1] |
| Northern Mexico 1 | 0.136[2] | - | 0.069[1] | 0.063[1] | 0.077[1] |
| Northern Mexico 2 | 0.063[2] | 0.064[2] | - | 0.033[1] | 0.042[1] |
| Texas and Mexico | 0.102[2] | 0.058[2] | 0.029[2] | - | 0.032[1] |
| Northern USA | 0.115[2] | 0.082[2] | 0.049[2] | 0.025[2] | - |

[1] Fst values from ADMIXTURE (Alexander, Novembre, and Lange 2009) K=5 results using the 466 native pecan samples genotyped with Illumina whole genome resequencing.
[2] Fst values calculated with pixy (Korunes and Samuk 2021). Samples assigned to a gene pool when they have >50% ancestry from that gene pool according to the K=5 ADMIXTURE results.

**Table S6: Gene pool divergence times estimated from *dadi***

| Gene Pools in Model | Model[1] | Optimization Method | Divergence 1; 15 year generations (Years in past) | Divergence 2;15 year generations (Years in past) | Divergence 1; 20 year generations (Years in past) | Divergence 2; 20 year generations (Years in past) |
|---|---|---|---|---|---|---|
| Central Mexico, Northern Mexico 1, Northern | divergence, no migration | standard | 191,187.5 | 179,714.4 | 254,916.6 | 239,619.1 |

| | | | | | | |
|---|---|---|---|---|---|---|
| USA | | | | | | |
| Central Mexico, Northern Mexico 1, Northern USA | divergence, no migration | log | 177,252.8 | 173,496.8 | 236,337.1 | 231,329.0 |
| Central Mexico, Northern Mexico 1, Northern USA | divergence, no migration, exponential growth | standard | 235,436.0 | 169,972.0 | 313,914.6 | 226,629.3 |
| Central Mexico, Northern Mexico 1, Northern USA | divergence, no migration, exponential growth | log | 153,781.8 | 147,115.8 | 205,042.4 | 196,154.4 |
| Northern Mexico 2, Texas and Mexico, Northern USA | divergence, no migration | standard | 88,174.2 | 87,415.1 | 117,565.6 | 116,553.4 |
| Northern Mexico 2, Texas and Mexico, Northern USA | divergence, no migration | log | 82,341.1 | 81,705.0 | 109,788.2 | 108,940.0 |
| Northern Mexico 2, Texas and Mexico, Northern USA | divergence, no migration, exponential growth | standard | 100,506.7 | 98,843.0 | 134,008.9 | 131,790.7 |
| Northern Mexico 2, Texas and Mexico, Northern USA | divergence, no migration, exponential growth | log | 74,414.4 | 67,459.3 | 99,219.2 | 89,945.7 |

[1] Demographic estimates using *dadi* (Gutenkunst et al. 2009).

**Table S7: Partial RDA (pRDA) forward selection table for the identification of non-redundant variables**

| Bioclimate Variable | Adjusted R-squared | F-value | P-value |
|---|---|---|---|
| Isothermality (Isotherm) | 0.01470842 | 4.9559 | 0.002 ** |
| Mean Temperature of the Warmest Quarter (Mean.Tmp.Wrm.Q) | 0.02030499 | 2.5081 | 0.002 ** |
| Annual Precipitation (Ann.Prc) | 0.02530542 | 2.3493 | 0.002 ** |
| Precipitation of the Warmest Quarter (Prc.Wrm.Q) | 0.027192 | 1.5082 | 0.002 ** |
| Precipitation of the Coldest Quarter Prc.Cld.Q) | 0.028535 | 1.3607 | 0.002 ** |
| Precipitation of the Wettest Quarter (Prc.Wet.Q) | 0.029731 | 1.3206 | 0.002 ** |
| Precipitation Seasonality (Prc.Seas) | 0.031047 | 1.3517 | 0.002 ** |
| Elevation | 0.032248 | 1.3202 | 0.002 ** |
| Mean Temperature of the Wettest Quarter (Mean.Tmp.Wet.Q) | 0.033323 | 1.2858 | 0.002 ** |
| Mean Temperature of the Driest Quarter (Mean.Tmp.Dry.Q) | 0.034179 | 1.2269 | 0.002 ** |
| Precipitation of the Wettest Quarter (Prc.Dry.Q) | 0.035013 | 1.2203 | 0.002 ** |
| Precipitation of the Driest Month (Prc.Dry.M) | 0.035981 | 1.2553 | 0.002 ** |

**Table S8: Partial RDA (pRDA) variance partitioning for decomposing the association of environment, geography, and demography on genetic variation of native pecan**

| Model | Inertia/Variance | R2<br>Adjusted R2 | F (df) p (>F) | Proportion of explainable variance |
|---|---|---|---|---|
| Full model | 7083 | 0.119691<br>0.05934723 | 1.9835 (17)<br>0.001*** | 1 |

| | | | | |
|---|---|---|---|---|
| Climate model | 3121 | 0.05273758 0.01033702 | 1.2381 (12) 0.001 *** | 0.44 |
| Neutral pop structure model | 1792 | 0.03029068 0.02073733 | 2.8445 (3) 0.001 *** | 0.25 |
| Geography model | 505 | 0.008530995 0.001517632 | 1.2017 (2) 0.001*** | 0.07 |

**Table S9: Genomic libraries included in 'Oaxaca' genome assemblies and assembled sequence coverage in final 'Oaxaca' V2 release**

| Library | Sequencing Platform | Average Read/Insert Size | Read Number | Assembled Sequence Coverage (x) |
|---|---|---|---|---|
| IPHL | Illumina-HiC (2x150) | N/A | 1,093,417,634 | 120.70 |
| IIMT | Illumina (2x150) | 400 | 931,146,802 | 101.90 |
| IIMU | Illumina (2x150) | 400 | 858,776,556 | 93.31 |
| PFBW | PACBIO/REVIO | 6,132* | 5,092,472 | 25.71 |
| **Total** | | N/A | 2,888,433,464 | 341.62 |

*Avg read length of PACBIO reads

**Table S10: PACBIO CCS library statistics and respective assembled sequence coverage levels**

| Cutoff | Number of Reads | Basepairs | Average Read Length | Coverage |
|---|---|---|---|---|
| 0 | 5,092,472 | 34,699,946,234 | 6,132 | 25.71x |
| 1,000 | 5,086,007 | 34,694,512,002 | 6,136 | 25.70x |
| 2,000 | 4,978,818 | 34,518,414,588 | 6,209 | 25.57x |
| 3,000 | 4,721,309 | 33,870,584,888 | 6,390 | 25.09x |
| 4,000 | 4,390,569 | 32,695,365,147 | 6,637 | 24.22x |
| 5,000 | 3,540,868 | 28,828,174,875 | 7,357 | 21.35x |
| 6,000 | 2,646,307 | 23,929,905,547 | 8,298 | 17.73x |
| 7,000 | 1,971,330 | 19,558,101,831 | 9,207 | 14.49x |
| 8,000 | 1,453,005 | 15,682,511,890 | 10,098 | 11.62x |
| 9,000 | 1,055,501 | 12,312,659,748 | 10,987 | 9.12x |

| | | | | |
|---|---|---|---|---|
| 10,000 | 751,752 | 9,434,161,665 | 11,890 | 6.99x |
| 11,000 | 525,249 | 7,061,674,385 | 12,799 | 5.23x |
| 12,000 | 360,151 | 5,167,503,146 | 13,709 | 3.83x |
| 13,000 | 241,857 | 3,692,230,117 | 14,639 | 2.73x |
| 14,000 | 159,149 | 2,578,315,573 | 15,585 | 1.91x |
| 15,000 | 103,264 | 1,769,778,436 | 16,528 | 1.31x |
| 16,000 | 65,787 | 1,190,166,417 | 17,491 | 0.88x |
| 17,000 | 41,383 | 788,352,221 | 18,464 | 0.59x |
| 18,000 | 25,903 | 518,070,090 | 19,410 | 0.39x |
| 19,000 | 15,936 | 334,086,529 | 20,375 | 0.25x |

**Table S11: Summary statistics of initial output of the 'Oaxaca' V2 HAP1 RACON polished HiFiAsm+HIC assembly**

| Minimum Scaffold Length | Number of Scaffolds | Number of Contigs | Scaffold Size | Basepairs | % Non-gap Basepairs |
|---|---|---|---|---|---|
| 5 Mb | 39 | 39 | 532,784,029 | 532,784,029 | 100.00% |
| 2.5 Mb | 62 | 62 | 622,368,601 | 622,368,601 | 100.00% |
| 1 Mb | 83 | 83 | 656,922,702 | 656,922,702 | 100.00% |
| 500 Kb | 90 | 90 | 662,568,589 | 662,568,589 | 100.00% |
| 250 Kb | 96 | 96 | 664,953,311 | 664,953,311 | 100.00% |
| 100 Kb | 119 | 119 | 668,648,347 | 668,648,347 | 100.00% |
| 50 Kb | 161 | 161 | 671,547,376 | 671,547,376 | 100.00% |
| 25 Kb | 161 | 161 | 671,547,376 | 671,547,376 | 100.00% |
| 10 Kb | 161 | 161 | 671,547,376 | 671,547,376 | 100.00% |
| 5 Kb | 161 | 161 | 671,547,376 | 671,547,376 | 100.00% |
| 2.5 Kb | 161 | 161 | 671,547,376 | 671,547,376 | 100.00% |

| | | | | | |
|---|---|---|---|---|---|
| 1 Kb | 161 | 161 | 671,547,376 | 671,547,376 | 100.00% |
| 0 bp | 161 | 161 | 671,547,376 | 671,547,376 | 100.00% |

**Table S12: Summary statistics of initial output of the 'Oaxaca' V2 HAP2 RACON polished HiFiAsm+HIC assembly**

| Minimum Scaffold Length | Number of Scaffolds | Number of Contigs | Scaffold Size | Basepairs | % Non-gap Basepairs |
|---|---|---|---|---|---|
| **5 Mb** | **48** | **48** | **559,017,296** | **559,017,296** | **100.00%** |
| **2.5 Mb** | **64** | **64** | **622,374,938** | **622,374,938** | **100.00%** |
| **1 Mb** | **83** | **83** | **659,814,800** | **659,814,800** | **100.00%** |
| **500 Kb** | **91** | **91** | **666,115,169** | **666,115,169** | **100.00%** |
| **250 Kb** | **100** | **100** | **669,215,956** | **669,215,956** | **100.00%** |
| **100 Kb** | **121** | **121** | **672,288,322** | **672,288,322** | **100.00%** |
| **50 Kb** | **158** | **158** | **674,730,332** | **674,730,332** | **100.00%** |
| **25 Kb** | **158** | **158** | **674,730,332** | **674,730,332** | **100.00%** |
| **10 Kb** | **158** | **158** | **674,730,332** | **674,730,332** | **100.00%** |
| **5 Kb** | **158** | **158** | **674,730,332** | **674,730,332** | **100.00%** |
| **2.5 Kb** | **158** | **158** | **674,730,332** | **674,730,332** | **100.00%** |
| **1 Kb** | **158** | **158** | **674,730,332** | **674,730,332** | **100.00%** |
| **0 bp** | **158** | **158** | **674,730,332** | **674,730,332** | **100.00%** |
